# Expanding the HDAC druggable landscape beyond enzymatic activity

**DOI:** 10.1101/2022.12.07.519454

**Authors:** Julien Olivet, Soon Gang Choi, Salvador Sierra, Tina M. O’Grady, Mario de la Fuente Revenga, Florent Laval, Vladimir V. Botchkarev, Christoph Gorgulla, Paul W. Coote, Jérémy Blavier, Ezekiel A. Geffken, Jimit Lakhani, Kijun Song, Zoe C. Yeoh, Bin Hu, Anthony C. Varca, Jonathan Bruyr, Samira Ibrahim, Tasneem Jivanjee, Joshua D. Bromley, Sarah K. Nyquist, Aaron Richardson, Hong Yue, Yang Wang, Natalia Calonghi, Alessandra Stefan, Kerstin Spirohn, Didier Vertommen, Maria F. Baietti, Irma Lemmens, Hyuk-Soo Seo, Mikhail G. Dozmorov, Luc Willems, Jan Tavernier, Kalyan Das, Eleonora Leucci, Alejandro Hochkoeppler, Zhen-Yu Jim Sun, Michael A. Calderwood, Tong Hao, Alex K. Shalek, David E. Hill, Andras Boeszoermenyi, Haribabu Arthanari, Sara J. Buhrlage, Sirano Dhe-Paganon, Javier González-Maeso, Franck Dequiedt, Jean-Claude Twizere, Marc Vidal

**Affiliations:** Center for Cancer Systems Biology, Dana-Farber Cancer Institute, Boston, MA, USA; Department of Genetics, Blavatnik Institute, Harvard Medical School, Boston, MA, USA; Department of Cancer Biology, Dana-Farber Cancer Institute, Boston, MA, USA; Laboratory of Viral Interactomes, Interdisciplinary Cluster for Applied Genoproteomics (GIGA)-Molecular Biology of Diseases, University of Liège, Liège, Belgium; Structural Biology Unit, Laboratory of Virology and Chemotherapy, Rega Institute for Medical Research, Department of Microbiology, Immunology and Transplantation, Katholieke Universiteit Leuven, Leuven, Belgium; Department of Physiology and Biophysics, Virginia Commonwealth University School of Medicine, Richmond, VA, USA; Laboratory of Gene Expression and Cancer, Interdisciplinary Cluster for Applied Genoproteomics (GIGA)-Molecular Biology of Diseases, University of Liège, Liège, Belgium; Laboratory of Cellular and Molecular Epigenetics, Interdisciplinary Cluster for Applied Genoproteomics (GIGA)-Cancer, University of Liège, Liège, Belgium; TERRA Teaching and Research Center, Gembloux Agro-Bio Tech, University of Liège, Gembloux, Belgium; Department of Biological Chemistry and Molecular Pharmacology, Blavatnik Institute, Harvard Medical School, Boston, MA, USA; Department of Physics, Faculty of Arts and Sciences, Harvard University, Cambridge, MA, USA; Broad Institute of Harvard and Massachusetts Institute of Technology, Cambridge, MA, USA; Institute for Medical Engineering and Science, Department of Chemistry and Koch Institute for Integrative Cancer Research, Massachusetts Institute of Technology, Cambridge, MA, USA; Ragon Institute of Massachusetts General Hospital, Massachusetts Institute of Technology and Harvard University, Cambridge, MA, USA; Laboratory of Biochemistry, Department of Pharmacy and Biotechnology, University of Bologna, Bologna, Italy; Laboratory of Biotechnology, Department of Pharmacy and Biotechnology, University of Bologna, Bologna, Italy; Consorzio per lo Sviluppo dei Sistemi a Grande Interfase, University of Firenze, Florence, Italy; de Duve Institute, Université Catholique de Louvain, Brussels, Belgium; TRACE Patient Derived Xenograft Platform, Department of Oncology, Leuven Cancer Institute, Katholieke Universiteit Leuven, Leuven, Belgium; Laboratory for RNA Cancer Biology, Department of Oncology, Leuven Cancer Institute, Katholieke Universiteit Leuven, Leuven, Belgium; Center for Medical Biotechnology, Vlaams Instituut voor Biotechnologie, Ghent, Belgium; Cytokine Receptor Laboratory, Department of Biomolecular Medicine, Faculty of Medicine and Health Sciences, Ghent University, Ghent, Belgium; Department of Biostatistics, Virginia Commonwealth University School of Medicine, Richmond, VA, USA; Division of Science and Math, New York University Abu Dhabi, Abu Dhabi, UAE

**Author notes:** These co-first authors contributed equally (Julien Olivet, Soon Gang Choi, Salvador Sierra, Tina M. O’Grady). These second authors contributed equally (Mario de la Fuente Revenga, Florent Laval, Vladimir V. Botchkarev, Christoph Gorgulla). Corresponding authors (Alex K. Shalek), (David E. Hill), (Andras Boeszoermenyi), (Haribabu Arthanari), (Sara J. Buhrlage), (Sirano Dhe-Paganon), (Javier González-Maeso), (Franck Dequiedt), (Jean-Claude Twizere), (Marc Vidal).

## Abstract

Enzymatic pockets such as those of histone deacetylases (HDACs) are among the most favored targets for drug development. However, enzymatic inhibitors often exhibit low selectivity and high toxicity due to targeting multiple enzyme paralogs, which are often involved in distinct multisubunit complexes. Here, we report the discovery and characterization of a non-enzymatic small molecule inhibitor of HDAC transcriptional repression functions with comparable anti-tumor activity to the enzymatic HDAC inhibitor Vorinostat, and anti-psychedelic activity of an *HDAC2* knockout *in vivo*. We highlight that these phenotypes are achieved while modulating the expression of 20- and 80-fold fewer genes than enzymatic and genetic inhibition in the respective models. Thus, by achieving the same biological outcomes as established therapeutics while impacting a dramatically smaller number of genes, inhibitors of protein-protein interactions can offer important advantages in improving the selectivity of epigenetic modulators.

**GRAPHICAL ABSTRACT:** 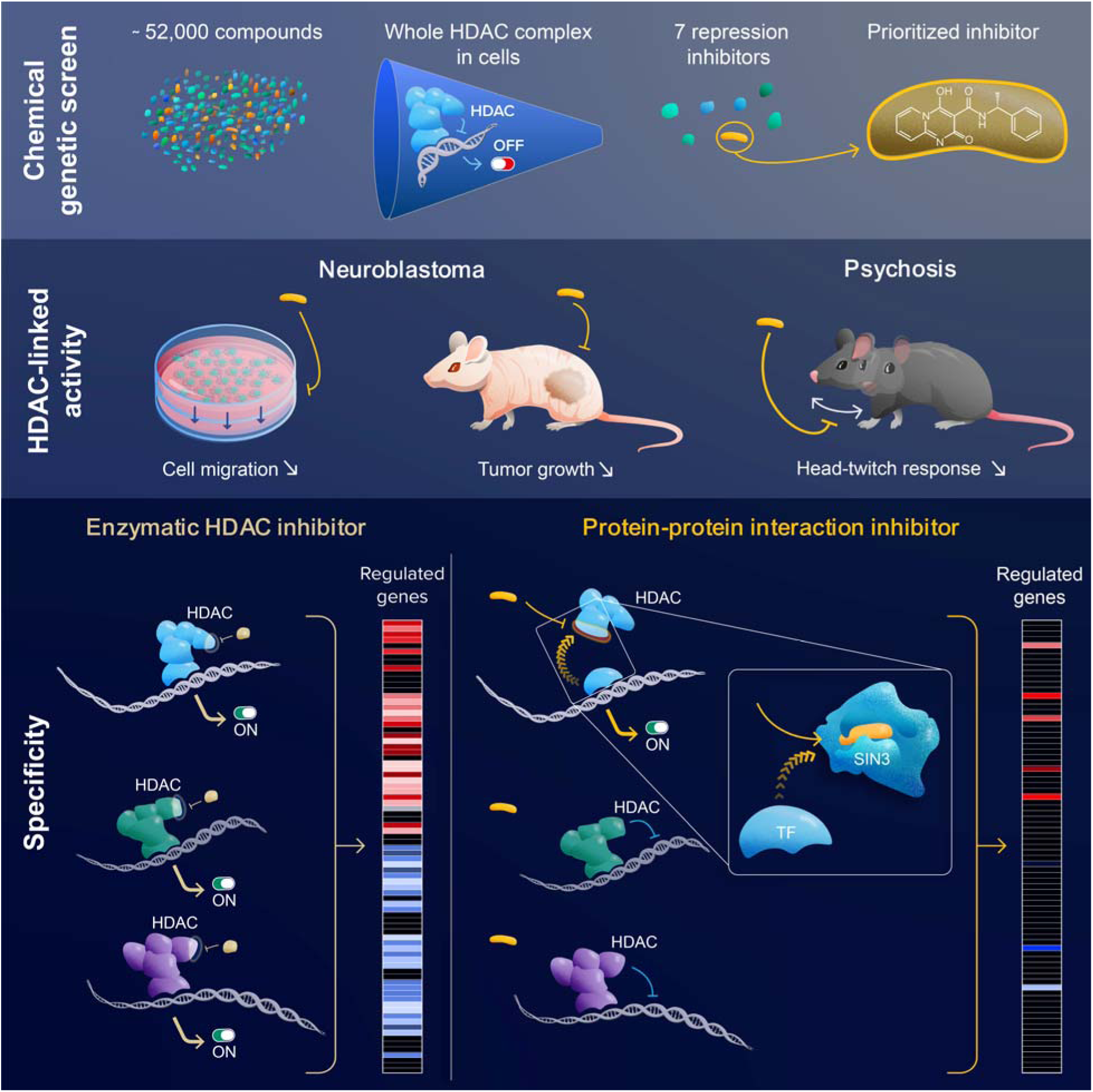

## INTRODUCTION

Strategies to discover novel therapeutics typically include targeting enzymatic activities involved in relevant biological pathways, with the goal of identifying potent, safe, and bioavailable drugs^1^. While catalytic pockets are attractive binding sites to identify such compounds, specific perturbation of enzymatic activity can be complicated by: i) the co-existence of numerous enzyme paralogs with distinct functions in unrelated pathways; and ii) the fact that many enzymes work as part of different multisubunit complexes for which associated proteins, and not the enzyme itself, determine substrate specificity and regulate enzymatic functions.

Human protein interactomes are emerging frontiers for the development of new therapeutics and the number of identified protein-protein interactions (PPIs) has already surpassed that of the druggable proteome by several orders of magnitude^2^. Yet, experimental tools and concepts are lagging behind to elevate PPI targets as a mainstream drug development ground, especially for high-order multisubunit complexes such as enzymatic complexes, which are abundant in the human proteome^3^. They thus represent attractive targets for emerging PPI modulator drugs with potentially greater selectivity.

This is particularly relevant for histone deacetylase (HDAC) complexes such as Sin3, CoREST, NuRD, SMRT/NCoR, and MiDAC, which are major epigenetic transcriptional regulators and for which identification of inhibitors originally raised great therapeutic hopes, especially for cancer treatments^4^. Unfortunately, despite huge resources and intense research invested in the last 30 years, and nearly a thousand clinical trials, most results have been disappointing and failed to meet the original promises. To date, only a handful of HDAC inhibitors have been approved for treatments of T-cell lymphoma, multiple myeloma, and breast cancer in combinatorial therapy, i.e. none of these molecules are currently used as first-line medicines^5–7^. While HDAC inhibitors were originally discovered via functional cell-based experiments^8^, the universal strategy used in the last decades has consisted of screening molecules in cell-free enzymatic assays^6^. This approach has delivered compounds that potently target deacetylase enzymatic pockets but display low selectivity *in vivo*, in that they tend to confer widespread effects on the transcriptome of treated cells, leading to derepression and in many cases deactivation of thousands of genes^9,10^. Such wide transcriptomic effects are thought to be at least in part responsible for serious adverse reactions^11^.

As alluded to above, most currently available inhibitors bind indiscriminately to the enzymatic pockets of several if not all of the 11 zinc-dependent HDAC paralogs encoded by the human genome^12^, which makes selective inhibition challenging^13,14^. In addition, even if an inhibitor specifically were able to target a single HDAC paralog, it would still disrupt the many complexes in which that particular enzyme is involved^15–17^, suggesting that higher selectivity is likely not achievable by solely targeting enzymatic activity.

Since the *in vivo* functions of all HDAC paralogs are carried on in the context of large protein complexes generally containing dozens of subunits and organized by large numbers of PPIs^18^, alternative strategies have emerged based on targeting particular domains involved in intra-complex interactions, such as the paired amphipathic helix (PAH) domains of SIN3, a scaffold protein of HDAC complexes^19^. This has allowed scientists to identify a handful of peptidomimetics^20,21^ and small molecule inhibitors with implications in Huntington’s disease^22^, breast cancer^23,24^, chronic pain^25^ or medulloblastoma^26^. Although these focused studies demonstrate that inhibiting intra-complex PPIs is a viable strategy, systematic PPI targeting within HDAC complexes has not been extensively investigated and the global effects of PPI inhibitors on whole cellular transcriptomes remain unknown.

Here, we have probed the druggable landscape of the conserved Sin3 HDAC complex by unbiasedly targeting its transcriptional repression functions with small molecules *in vivo*. We have identified structurally unique repression inhibitors that leave HDAC enzymatic activity unperturbed using a genetically engineered growth selection. We show that one of these molecules binds to the conserved PAH2 domain of the SIN3 scaffold protein, disrupting its interaction with a DNA-binding protein and recruitment of the HDAC complex to specific genes. This PPI inhibitor achieves anti-tumor and anti-psychedelic activities that parallel those of the FDA-approved HDAC enzymatic inhibitor Vorinostat, and of an *HDAC2* knockout mouse model, respectively. Remarkably, these beneficial effects are achieved while altering the expression of up to two orders of magnitude fewer genes compared to classical HDAC enzymatic inhibition. Our findings pave the way towards the expansion of the druggable space of macromolecular complexes, and will help improving the identification of alternative targets for the development of more selective and effective epigenetic modulators.

## RESULTS

### Targeting core subunits of the Sin3 HDAC complex

To interrogate the effect of small molecules on the repressive activity of an HDAC complex *in vivo,* we needed a model system that conserves critical features observed in human but in which positive selection for cell-permeable and non-toxic compounds could easily be implemented at large scale. Inspired by the original genetic identification of key genes involved in HDAC complexes, we selected yeast as a model system (**Figure 1A**). Indeed, the first HDAC-encoding gene ever identified, *RPD3,* was found, together with *SIN3,* in an unbiased yeast genetic screen based on *in vivo* growth selection for mutants that derepress the transcription of *TRK2,* a K transporter gene^27–30^ Similar screens or selections using a handful of other regulated genes such as *HO, INO1, SPO13, IME2* or *CAR1* also identified *RPD3* and/or *SIN3*^31^, in addition to other genes such as *UME6, UME1,* or *DEP1,* now known to encode subunits of an HDAC complex referred to here as the Sin3/Rpd3 Large (Sin3/Rpd3L) complex altogether containing 12 subunits^32^. The Sin3/Rpd3L HDAC complex is recruited to specific promoters through Ume6, a transcription factor (TF) that binds to DNA motifs such as the upstream repressing sequence (*URS*)^33,34^ (**Figure 1A**).

**Figure 1.**
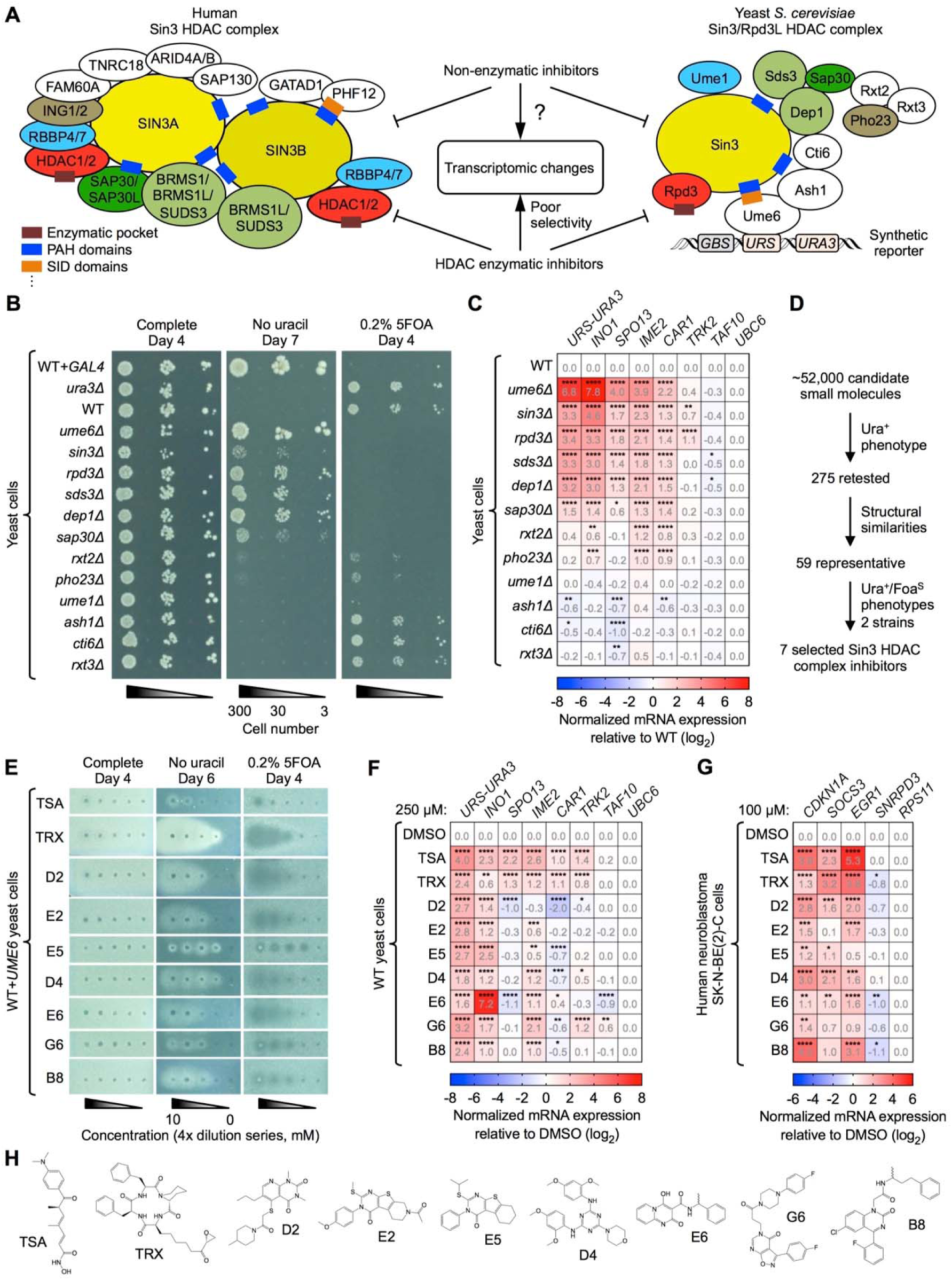
Identification of structurally unique inhibitors derepressing Sin3 HDAC complex-regulated genes. (**A**) Models of the human Sin3 HDAC^18^ and yeast *S. cerevisiae* Sin3/Rpd3L HDAC^32^ complexes with some protein domains indicated. The synthetic *URS-URA3* reporter gene in yeast is represented. Matching colored subunits indicate orthologous proteins. White subunit proteins lack full-length orthologs. **(B)** Phenotypes of the yeast Sin3/Rpd3L HDAC complex subunit deletion mutants in the *URS-URA3* reporter assay. **(C)** RT-qPCR analysis of yeast Sin3/Rpd3-regulated genes *(INO1, SPO13, IME2, CAR1, TRK2)* and genes not known to be regulated by Rpd3 *(TAF10, UBC6),* compared to *URS-URA3* for the deletion mutants presented in (B) (n = 3 biologically independent samples per group). **(D)** Small-molecule screening pipeline using the *URS-URA3* reporter assay in *WT+UME6* yeast cells. **(E)** Phenotypes of TSA, TRX, and the seven Sin3 HDAC complex inhibitors in the *URS-URA3* reporter assay. Uncropped plate pictures (E) are provided as **Figure S1F**. **(F)** RT-qPCR analysis of genes from (C) for TSA, TRX and the seven Sin3 HDAC complex inhibitors (n = 4 biologically independent samples per group). **(G)** RT-qPCR analysis of known human Sin3 HDAC complex-regulated genes *(CDKN1A, SOCS3, EGR1)* and genes not known to be regulated by a Sin3 HDAC complex *(SNRPD3, RPS11)* in neuroblastoma SK-N-BE(2)-C cells for TSA, TRX and the seven Sin3 HDAC complex inhibitors (n ≥ 3 biologically independent samples per group). **(H)** Structure of TSA, TRX and the seven small molecule inhibitors used in (E-G). RT-qPCR values for *UBC6* (C and F), and *RPS11* (G) were used to normalize data. Values represent means of replicates (C, F and G). Statistical analyses, (C, F and G) two-way Analysis of Variance (ANOVA) with Dunnett’s multiple comparison post-test relative to WT (C) or DMSO (F and G). Raw data are provided in **Source Data**.

To study Sin3/Rpd3L HDAC complex inhibition at a specific locus and identify small molecules that perturb such functions, we engineered a screening assay using the synthetic *SPAL10::URA3* reporter gene from MaV yeast strains^33^. An *URS* DNA motif was integrated downstream of Gal4-binding sites *(GBS)* and upstream of a chimeric *URA3* cassette (*URS-URA3*)^33^ to allow recruitment of the Sin3/Rpd3L HDAC complex via Ume6 (**Figure 1A**). In this context, a functional Sin3/Rpd3L HDAC complex represses *URA3,* inducing uracil auxotrophy in yeast cells. Thus, comparable to cells lacking the *URA3* gene *(ura3Δ*), yeast cells carrying the *URS-URA3* cassette and defined as wild-type (WT) in this study, do not grow on media lacking uracil (Ura_−_ phenotype) but do grow on media containing 5-fluoroorotic acid (5FOA) (Foa^R^ phenotype) (**Figure 1B**). We then evaluated the contribution of each component to the repressive activity of the complex by generating yeast strains in which subunit encoding-genes were individually deleted. We assessed associated cell growth on media lacking uracil or containing 5FOA and observed that six core subunits of the Sin3/Rpd3L HDAC complex are required to maintain *URS-URA3* repression: Ume6, Sin3, Rpd3, Sds3, Dep1, and Sap30 (**Figure 1B**). Indeed, their deletions confer strong Ura^+^/Foa-sensitive (Foa^S^) phenotypes, similar to cells in which *URA3* is activated via ectopic expression of the Gal4 TF *(WT+GAL4).* Interestingly, knocking out *RXT2* or *PHO23* was associated with intermediate Ura_+_/Foa^S^ phenotypes while deletions of other subunits led to Ura^-^/Foa^R^ phenotypes, suggesting that they are not required for repression at *URS-URA3.* Reverse transcription-quantitative PCR (RT-qPCR) further confirmed that repression of the *URS-URA3* locus is relieved when deleting any of the six core subunits of the Sin3/Rpd3L HDAC complex (**Figure 1C**), mirroring the observed growth phenotypes.

To compare the impact of the deletions on the synthetic *URS-URA3* reporter gene to established endogenous Sin3/Rpd3-regulated genes^34^, we measured expression of *INO1, SPO13, IME2, CAR1* and *TRK2* in the different deletion mutant strains. To control for specificity, two genes not known to be regulated by Rpd3, *UBC6* and *TAF10,* were also included. *INO1, SPO13, IME2* and *CAR1* were all significantly derepressed by the same six deletions as the synthetic *URS-URA3* locus, while *TRK2* was derepressed only in *sın3Δ* and *rpd3Δ* strains, consistent with published observations^28^ (**Figure 1C**). Interestingly, deleting *UME6* increased *INO1* expression by ∼230-fold, while deleting *RPD3* only had a ∼10-fold effect. In addition, deleting *ASH1, CTI6,* or *RXT3* decreased expression levels of *SPO13* and in some cases *CAR1,* suggesting their potential involvement in transcriptional activation mechanisms. Together, these results indicate that the Sin3/Rpd3L HDAC complex can adopt different compositions at different loci, and that using *URS-URA3* as a reporter gene should allow for the identification of compounds targeting core subunits of the complex.

To evaluate if the synthetic *URS-URA3* reporter gene would respond to established HDAC enzymatic inhibitors, we tested Trichostatin A (TSA) and Trapoxin A (TRX)^6,35^ We used the MaV208 yeast strain mutated for *PDR5* and *SNQ2,* two genes encoding drug efflux pumps known to be responsible for most pleiotropic drug resistance in *S. cerevisiae*^36^ (**Figure S1A**). To maximize the signal-to-noise ratio of the *URS-URA3* reporter assay, a plasmid encoding the Ume6 TF was transformed into WT haploid cells (WT+*UME6*), which decreased the frequency of Ura_+_ mutant colonies spontaneously growing on uracil dropout media by ∼100-fold (**Figure S1B**). Notably, this extra copy of *UME6* did not modify expression at *URS-URA3* (**Figure S1C**). Using an agar diffusion assay (**Note S1**), we found that both TSA and TRX led to Ura^+^/Foa^S^ phenotypes while having little or no effect on overall cell viability on complete media (**Figure S1D**). The effect was stronger for TRX than for TSA, which might be related to their different binding modes to HDAC catalytic pockets^35^.

### Selected small molecules derepress Sin3 HDAC complex-regulated genes

To test the ability of various chemical structures to disrupt the repressive activity of the yeast Sin3/Rpd3L HDAC complex at the *URS-URA3* locus, we interrogated ∼52,000 compounds using the JOY134 (WT+*UME6*) strain (**Figure 1D**). We selected compounds from diverse, high-quality chemical libraries and included known bioactives such as commercially-available drugs, natural products, molecules with PPI inhibition properties, diverse synthesized compounds containing complex three-dimensional chemotypes, and annotated molecules with drug-like properties and potential for structure-activity relationship (SAR) experiments (**Table S1**). Of the ∼52,000 tested compounds, 275 led to Ura_+_ phenotypes of varying intensities, resulting in a ∼0.5% hit rate. Following clustering based on structural similarities, 59 representative compounds were prioritized according to criteria including: i) potency in the primary assay, ii) absence of reactive chemical groups, and iii) absence of metabolic liabilities such as functional groups being prone to first pass metabolism by cytochrome P450 enzymes. To select for the most robust compounds, we retested them in the original JOY134 yeast cells, as well as in the JOY201 (WT2+*UME6*) strain in which the *SPAL10::URA3* reporter system was reconstructed *de novo* (**Figure S1E**). Seven compounds consistently conferred strong Ura^+^/Foa^S^ phenotypes in both strains and in a dose-dependent manner (**Figure 1E and Figures S1F and S1G**), and were thus selected for further studies.

To compare the transcriptional responses of the selected molecules to those of TSA and TRX, we measured their effects on the expression of the same genes used to assess the contribution of each subunit on the repressive activity of the yeast Sin3/Rpd3L HDAC complex (**Figure 1C**). As expected, all compounds significantly derepressed the *URS-URA3* reporter gene with effects ranging from ∼3 to 16-fold (**Figure 1F**). While TSA and TRX derepressed the representative Sin3/Rpd3-regulated loci as expected, compounds from the screen showed higher selectivity and derepressed only some genes. Interestingly, while *INO1* expression was generally increased by ∼1.5 to 6-fold, the effect of compound E6 was two orders of magnitude higher (∼150-fold) than any other treatment, including TSA and TRX. These results indicate that in contrast to established enzymatic HDAC inhibitors, these new chemical probes may affect only specific Sin3/Rpd3L HDAC complex-regulated loci.

Finally, to evaluate if the selected compounds would target the conserved Sin3 HDAC complex repressive activity in human cells, we measured their transcriptional effects in neuroblastoma SK-N-BE(2)-C cells shown to be sensitive to HDAC inhibition^37,38^. To benchmark compounds, we used *CDKN1A*, *SOCS3*, and *EGR1*, three genes known to be regulated by a Sin3 HDAC complex. To control for specificity, two genes not known to be regulated by HDAC, *RPS11* and *SNRPD3*, were also included. As expected, TSA and TRX both derepressed all the tested human Sin3 HDAC complex-regulated loci while compounds identified with the *URS-URA3* reporter assay also induced derepression responses (**Figure 1G**). Interestingly, none of these molecules have structural features characteristic of HDAC enzymatic pocket binders such as TSA or TRX^4^ (**Figure 1H**), suggesting that they might constitute a new class of non-enzymatic inhibitors targeting evolutionary conserved subunits of the Sin3 HDAC complex.

### Selected small molecules act as non-enzymatic HDAC complex inhibitors

To understand the mechanisms by which the new compounds might perturb Sin3 HDAC complex repression functions, we first tested their ability to inhibit enzymatic activity using an established luminescence-based assay (**Note S2**) and three different sources of HDAC enzymes: i) live MaV208 WT yeast cells, ii) human HeLa nuclear extracts, and iii) human HDAC1 complexes purified from HEK293T cells and containing the Sin3 HDAC complex as verified by mass-spectrometry (**Table S2**)^16^ While known HDAC enzymatic inhibitors including TSA and TRX systematically reduced HDAC activity from all three sources, none of our prioritized compounds showed any effect (**Figure 2A**). Together with the lack of structural features typically found in HDAC enzymatic pocket binders (**Figure 1H**), these experiments strongly suggest that the compounds identified in the phenotypic screen do not act by inhibiting HDAC enzymatic activity.

**Figure 2.**
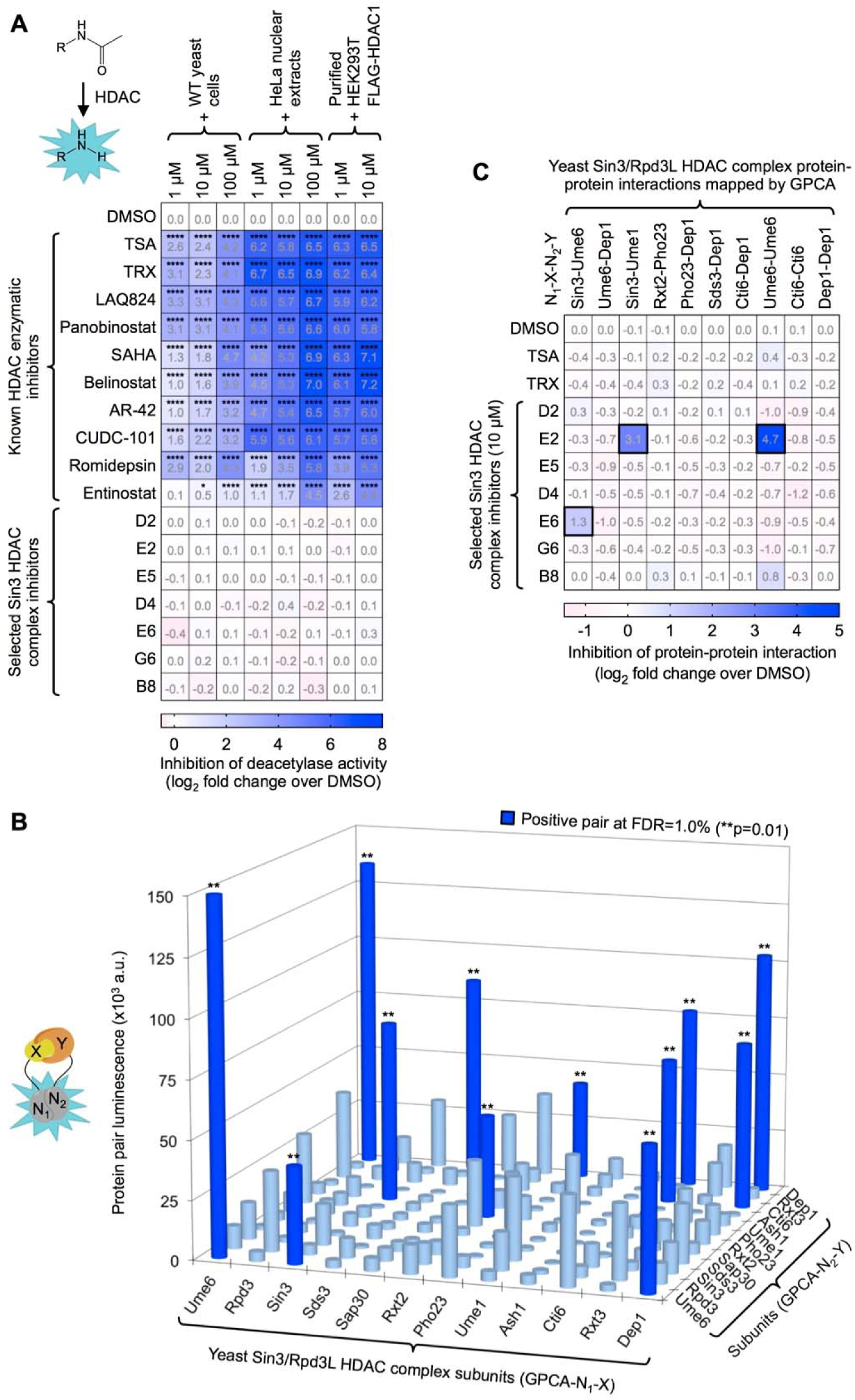
Selected small molecules act as non-enzymatic HDAC complex inhibitors. **(A)** Inhibition of deacetylase enzymatic activity by ten classical HDAC enzymatic inhibitors and the seven selected Sin3 HDAC complex inhibitors (n ≥ 3 biologically independent samples per group). Values represent means of replicates. **(B)** Mapping binary Sin3/Rpd3L HDAC complex subunit-subunit interactions in a all-by­all format (12 x 12 matrix) using the N_1_N_2_ GPCA versions. Protein pairs scoring above the FDR cutoff of 1.0% are indicated by dark blue cylinders, and light blue cylinders correspond to pairs scoring below the cutoff. Bars represent individual data points. The y-axis scale is cut at 150×10^3^ arbitrary units (a.u.). **(C)** Inhibition of binary PPIs mapped in (B) by TSA, TRX, and the seven Sin3 HDAC complex inhibitors assessed by GPCA (n ≥ 7 biologically independent samples per group for DMSO, n = 1 for the tested small molecules). Values represent means of DMSO replicates or individual data points for the tested small molecules. A black frame indicates a data point scoring below three standard deviations from the DMSO mean for a given PPI. Statistical analyses, (A) two-way ordinary ANOVA with Dunnett’s multiple comparisons post-test relative to DMSO, and (B) based on empirical p value. Raw data are provided in **Source Data**.

To test for alternative mechanisms, we investigated whether our compounds could inhibit Sin3/Rpd3L HDAC complex-mediated transcriptional repression in yeast by perturbing intra-complex PPIs. First, we systematically mapped binary interactions between the 12 full-length subunits of the complex under highly specific and sensitive conditions^39^ using the mammalian cell-based *Gaussia princeps* complementation assay (GPCA) (**Figures S2A and S2B**). Ten PPIs were identified by GPCA, including seven heterodimeric and three homodimeric interactions (**Figure 2B**), among which 50% were validated with the orthogonal kinase substrate sensor (KISS) assay (**Figures S2C and S2D**). We then used GPCA to test the ability of TSA, TRX, and each of the selected compounds to perturb binary interactions within the Sin3/Rpd3L HDAC complex. We found that only two molecules disrupted subunit-subunit interactions, including the racemic compound E6 that selectively inhibits the well-documented Sin3-Ume6 PPI (**Figures 2C and S2E**) driving recruitment of the complex to specific genes^40^. Due to its potential higher selectivity towards the disruption of the Sin3/Rpd3L HDAC complex at loci containing Ume6 TF binding site(s), compound E6 was prioritized for further studies.

To evaluate if E6 disruption of the Sin3-Ume6 interaction prevents recruitment of the HDAC complex to its target loci, we performed chromatin immunoprecipitation (ChIP) experiments in a *sın3Δ* yeast strain expressing an HA-tagged Sin3^40^ (**Figure S2F**). Using ChIP-qPCR we verified that E6 significantly decreased the presence of HA-tagged Sin3 at *URS-URA3* and *INO1* (**Figure S2G**). Interestingly, TSA also appeared to prevent recruitment of Sin3, in agreement with studies suggesting that enzymatic inhibitors can lead to dynamic remodeling of HDAC complexes^41^. To compare the effects of enzymatic (TSA) and PPI (E6) inhibitors on the recruitment of Sin3 on a genome-wide scale, we performed ChIP-sequencing (ChIP-seq) experiments. In the absence of inhibitors, Sin3 was detected at the promoter of 545 genes (**Figure 3A**). Importantly, there were significant overlaps between Sin3-occupied genes and the 132 genes containing at least one *URS*-binding site motif in their promoter extracted from a published database^42^, and the 361 genes bound by the Ume6 TF obtained from the literature^43^ (**Figure 3A**). Treating yeast cells with TSA significantly decreased the presence of Sin3 at 295 genes, while compound E6 reduced Sin3 occupancy at 450 genes (**Figure 3B**). Interestingly, upon examination of the *URS*-containing and/or Ume6 bound genes, we found that E6 did not dissociate Sin3-containing complexes at every locus (**Figure 3C-E**), as illustrated by ChIP-seq tracks for three Ume6-bound genes that also contain a *URS* motif in their promoter (**Figure 3F**).

**Figure 3.**
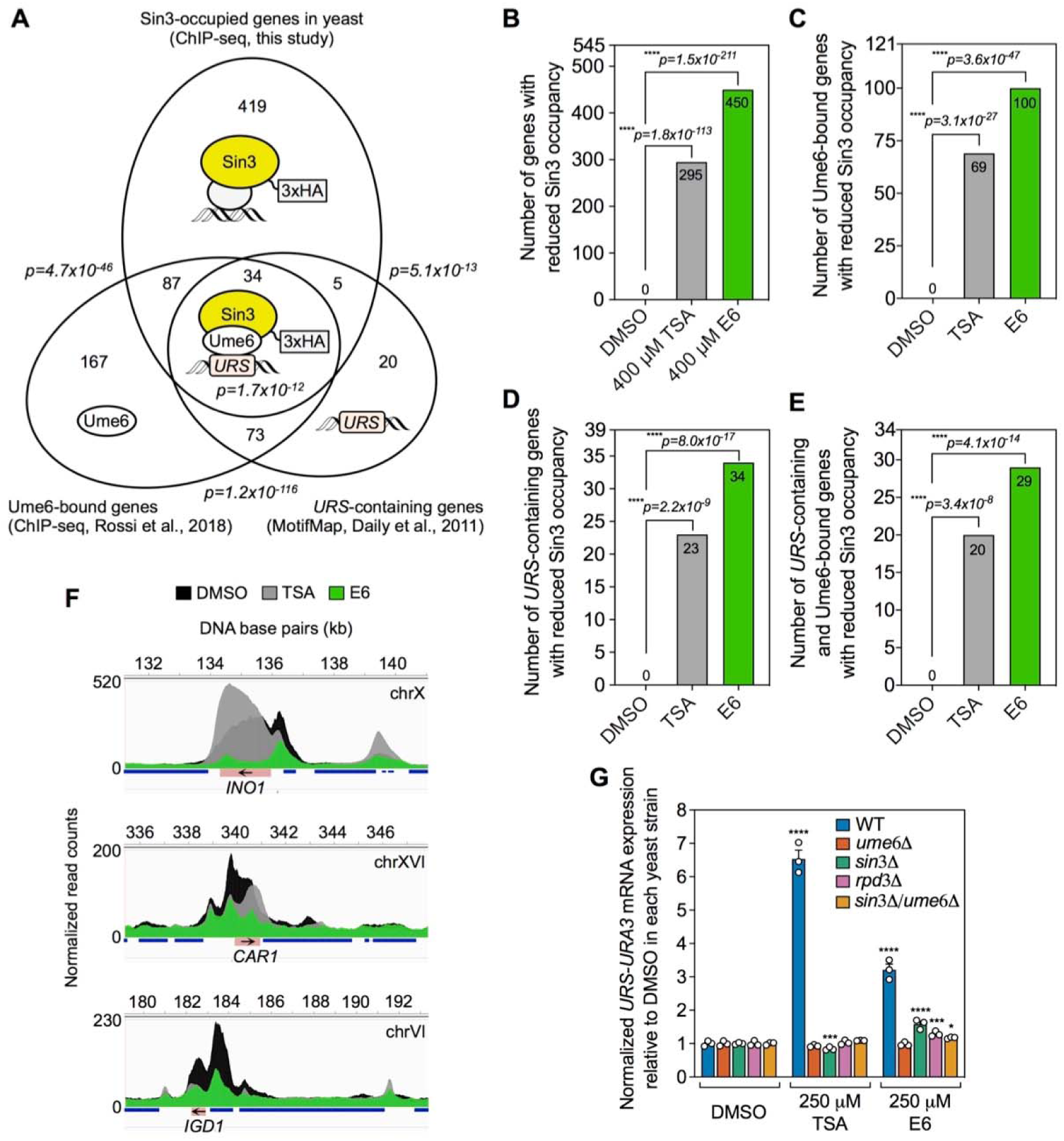
E6 inhibitor prevents recruitment of the yeast Sin3/Rpd3L HDAC complex at specific genes. (**A**) Venn diagrams between Sin3-occupied genes from ChIP-seq experiments in this study (n = 2 biologically independent samples per group; overlap between replicates were used to analyze data), *URS*-containing, and Ume6-bound genes from the literature. (**B-E**) Number of (B) yeast genes, (C) Ume6-bound genes, (D) *URS*-containing genes, and (E) Ume6-bound and *URS*-containing genes where Sin3 occupancy is reduced by TSA or E6 treatment compared to the DMSO control. (**F**) Examples of ChIP-seq tracks for three Ume6-bound and *URS*-containing genes represented by red rectangles. Arrows indicate the sense of the DNA strand and surrounding genes are represented by blue rectangles. **(G)** RT-qPCR analysis of TSA and E6 effects in the WT, *ume6Δ*, *sin3Δ*, *rpd3Δ*, or *sin3Δlume6Δ* yeast strains (n = 3 biologically independent samples per group). Symbols represent independent repeats, bars represent means of replicates and error bars SEM. RT-qPCR values for *UBC6* were used to normalize data. Statistical analyses, (G) two-way ordinary ANOVA with Dunnett’s multiple comparisons post-test relative to DMSO. Raw data are provided in **Source Data**.

Finally, to determine if derepression of the *URS-URA3* reporter gene by compound E6 was dependent on a functional, fully-assembled Sin3/Rpd3L HDAC complex, we tested its effect in the following yeast strain backgrounds: WT, *rpd3Δ*, *sın3Δ*, *ume6Δ* and *sin3Δlume6Δ* (**Figure 3G**). As expected, TSA had no additive effect on *URS-URA3* expression in the mutant strains, further demonstrating the epistasicity of the tested deletions with this inhibitor. Similarly, the absence of these subunits generally abolished the ability of E6 to derepress the *URS-URA3* locus, suggesting that the transcriptional activation effects of this compound rely on a fully assembled, functional Sin3/Rpd3L HDAC complex.

### E6R perturbs transcription by binding to the SIN3 PAH2 domain and inhibiting the PAH2-SID interaction

The yeast Sin3-Ume6 interaction is mediated by the paired amphipathic helix 2 (PAH2) of Sin3 and the Sin3-interacting domain (SID) of Ume6^44^. Both domains are highly conserved in mammalian proteins and allow similar assemblies between SID-containing transcription factors and the SIN3A and SIN3B scaffold subunits^19,45,46^. To characterize this interaction interface, we determined the crystal structures of the yeast Sin3 PAH2 domain alone (**Figure S2H and Table S3**), or in complex with the Ume6 SID peptide (**Figure 4A-C and Table S3**). Structures were determined at 2.2 and 1.8 Å resolutions, respectively. Similar to published solution structures of mammalian SIN3A PAH2^45^ and SIN3B PAH2^46^, the PAH2 domain of yeast *S. cerevisiae* Sin3 adopts a left-handed, anti-parallel, four-helix bundle configuration, two of which (helices α1 and α2) define a deep hydrophobic pocket that accommodates the amphipathic Ume6 α helix peptide (**Figure 4A-C**). With a root-mean-square deviation (RMSD) between backbone atoms of 1.7 Å^2^, the free and complexed structures did not reveal significant conformational changes.

**Figure 4.**
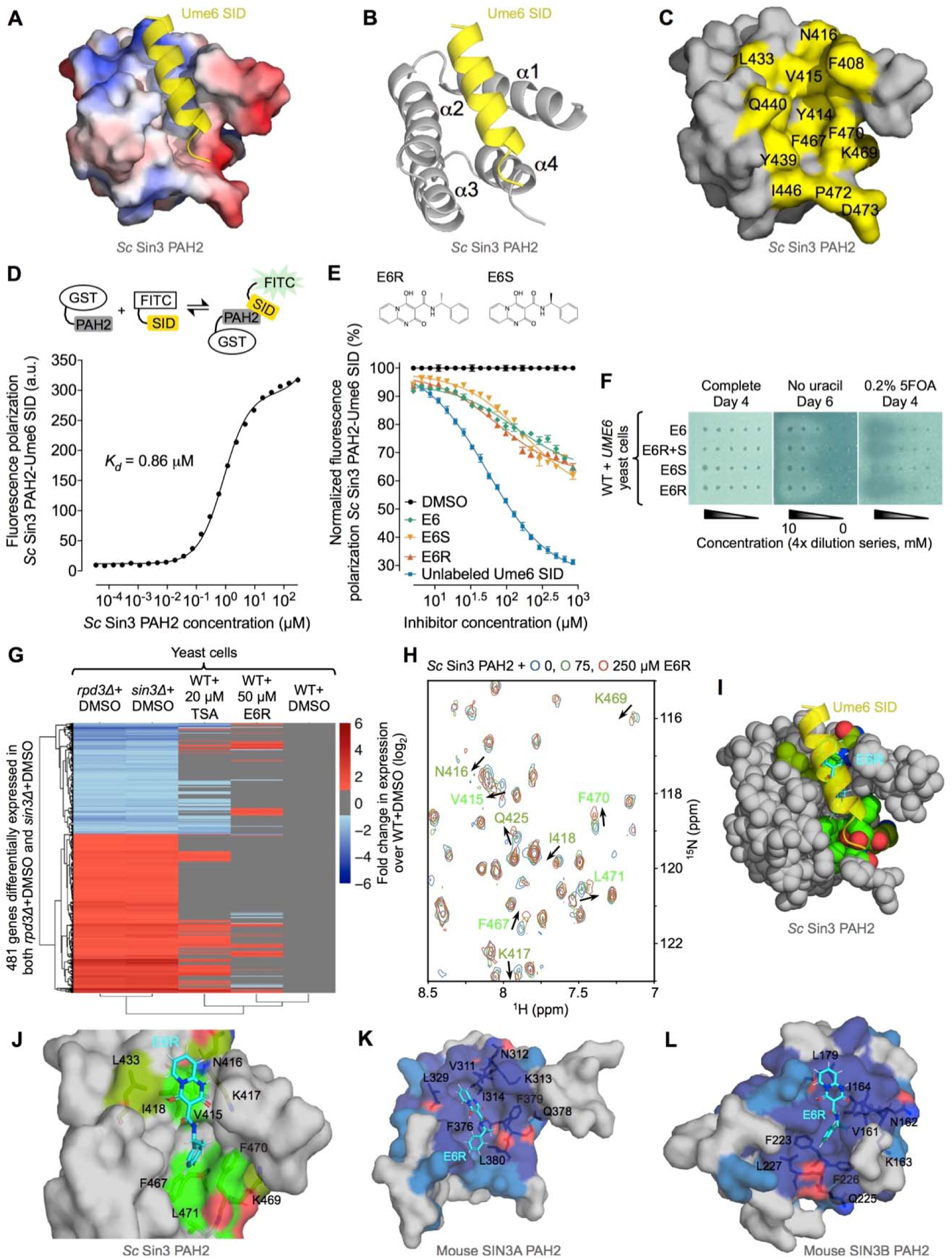
E6R perturbs transcription by binding to the SIN3 PAH2 domain and inhibiting the PAH2-SID interaction. **(A)** Surface representation of the yeast *S. cerevisiae (Sc)* Sin3 PAH2 domain in complex with the Ume6 SID peptide at 1.8 Å resolution. The electrostatic surface is presented with a gradient from -3 (red) to +3 (blue) KT/e. **(B)** Ribbon diagram of the *Sc* Sin3 PAH2-Ume6 SID structure with alpha helices indicated. **(C)** Surface representation of the *Sc* Sin3 PAH2 domain with residues in contact with the Ume6 SID peptide indicated in yellow. (**D and E**) FP titration curves showing (D) the interaction of *Sc* Sin3 PAH2 domain (GST-tagged) with Ume6 SID (FITC-labeled) (n = 4 biologically independent samples per group), and (E) inhibition of the *Sc* Sin3 PAH2-Ume6 SID interaction by unlabeled Ume6 SID, E6, E6S, or E6R (n = 3 biologically independent samples per group). Symbols represent means with SEM, and lines fitted curves. **(F)** Phenotypes of commercial E6, synthesized E6R+S, E6S and E6R in the *URS-URA3* reporter assay. **(G)** Effect of TSA and E6R on Sin3/Rpd3L HDAC complex-regulated genes (RNA-seq) in yeast (n = 3 biologically independent samples per group). For each condition, a row represents a gene log_2_(fold change). **(H)** ^15^N-SOFAST HMQC NMR spectra of untagged *Sc* Sin3 PAH2 (25 µM) with or without E6R. A black arrow indicates the direction of a peak shift with increasing E6R concentrations. **(I)** Space filling model of the *Sc* Sin3 PAH2 domain (grey) where carbon atoms of residues experiencing E6R chemical shift perturbations above 0.05 and 0.025 ppm in (H) are colored in light and dark green, respectively. Oxygen and nitrogen atoms in perturbed residues are shown in red and blue, respectively. The Ume6 SID peptide is visualized as a semi-transparent yellow helix and superposed with the docked E6R structure with a cyan carbon backbone where the red, blue and white colors correspond to oxygen, nitrogen and hydrogen atoms, respectively. **(J)** Semi-transparent surface representation of the *Sc* Sin3 PAH2 domain with E6R perturbed residues shown as sticks following the color codes from (I). (**K and L**) Docking of E6R into the mouse SIN3A PAH2 (PDB: 2L9S) (K), and SIN3B PAH2 (PDB: 2F05) (L) domains where deep blue corresponds to residues conserved across all SIN3 PAH2 homologs and sky blue to semi-conserved residues. Amino acids shown as sticks correspond to NMR perturbed residues by E6R in yeast and their mouse pendants. Raw data are provided in **Source Data**.

To examine inhibition of the isolated Sin3-Ume6 interaction by compound E6, we used a fluorescence polarization (FP) assay. By adding increasing amounts of GST-Sin3 PAH2 to a fixed concentration of FITC-Ume6 SID, we estimated the interaction K_D_ as 0.86 ± 0.02 µM (**Figure 4D**). We then used FP to test the effect of E6 and synthesized E6R and E6S enantiomers on the Sin3 PAH2-Ume6 SID interaction (**Figure 4E**). The unlabeled Ume6 SID control inhibited the interaction in a dose-dependent manner, with an IC_50_ of 53.3 ± 2.4 µM. The E6 racemic mixture was also able to displace the FITC-Ume6 SID peptide from Sin3 PAH2 with an estimated IC_50_ of 132.5 ± 19.8 µM, calculated as previously reported^47^. The E6R enantiomer was more potent than E6S, with estimated IC_50_ values of 74.5 ± 7.2 µM versus 174.5 ± 21.9 µM, respectively. With an assumption of competitive inhibition, these IC_50_ values correspond to inhibitory concentrations (K_i_) of 11.4, 29.1, 38.5 and 16.1 µM for unlabeled Ume6 SID, E6, E6S and E6R, respectively. Similar differences between E6R and E6S were observed when labeling the Ume6 SID peptide with TAMRA, a different fluorescent probe (**Figures S2I and S2J**). Taken together, these results led us to conclude that E6R can compete for the Ume6 SID binding site on the Sin3 PAH2 domain. These findings were consistent with the *URS-URA3* phenotypic assay as, compared to E6S, the E6R enantiomer conferred: i) stronger rings of growth on media lacking uracil; ii) stronger growth inhibition on media containing 5FOA; and iii) stronger derepression of *URS-URA3* (**Figures 4F and S3A**). Based on these observations, E6R was prioritized for follow-up studies and its transcriptional effects were compared to that of TSA on the subset of Sin3/Rpd3L HDAC complex-regulated genes in yeast following compound titrations (**Figure S3A**). Interestingly, while RNA-sequencing (RNA-seq) analyses revealed that TSA altered expression of 45% of the Sin3/Rpd3-dependent genes, E6R significantly affected expression of only 20% of these genes, indicating a greater selectivity towards Sin3/Rpd3L HDAC complex-regulated loci (**Figures 4G and S3B-D**).

To test the binding of E6R to the PAH2 domain of yeast Sin3, we used three ligand-detected nuclear magnetic resonance (NMR) methods: differential line broadening (DLB), saturation transfer difference (STD), and Carr–Purcell–Meiboom–Gill-transverse relaxation rate (CPMG-R_2_). First, compared to the NMR spectrum of free E6R, significant line broadening and chemical shift perturbations were observed in the presence of Sin3 PAH2, indicating a bound state for E6R (**Figure S4A**). Second, on-resonance saturation of Sin3 PAH2 decreased the intensity of E6R signals in the STD experiment, indicating transmission of magnetization from Sin3 PAH2 to interacting E6R ligands (**Figure S4B**). Finally, the presence of Sin3 PAH2 enhanced the relaxation rate of the E6R signal in the CPMG-R_2_ experiments, also indicative of E6R binding to the receptor protein (**Figures S4C and S4D**).

To confirm the specificity of this binding and identify amino acids interacting with E6R, we studied chemical shift perturbations in residues of the Sin3 PAH2 domain following addition of the compound. After assigning the majority of the peaks (85%) to the different amino acids using a ^13^C- and ^15^N-labeled yeast Sin3 PAH2 construct by NMR (**Figures S4E and S4F**), we confirmed that the secondary structures in solution were consistent with the tertiary one determined previously (**Figure S2H**). As indicated by NMR chemical shift perturbations, the binding of E6R to Sin3 PAH2 was localized to only a few amino acids lining the conserved Ume6 SID binding pocket, such as Val415, Phe467, Phe470 and Leu471 (**Figures 4H, S4G, S4H and S5A**). To model the binding of E6R, we virtually docked it against the region of the yeast Sin3 PAH2 domain that binds to the Ume6 SID peptide (**Figure 4I**). Similar to the binding of Ume6 SID to the Sin3 PAH2 domain and in agreement with the NMR results (**Figure 4H**), E6R is predominantly engaged in hydrophobic interactions with the Sin3 PAH2 domain (**Figure 4J**), with predicted distances between interacting moieties below 4 Å. Altogether, these results establish E6R as a Sin3 PAH2 binder capable of competing with the Ume6-SID peptide.

Finally, using ligand-detected NMR, we confirmed that E6R could bind to the conserved PAH2 domains (**Figure S5A**) of human SIN3A (**Figure S5B-E**) and SIN3B (**Figure S5F-I**). We then modeled the binding of E6R to the mammalian SIN3A and SIN3B PAH2 domains by docking it against all NMR conformations published for those proteins^45,46^. This revealed similar binding poses compared to the yeast model since, in both cases, E6R was predicted to bind to the same hydrophobic pocket, in the same orientation and by involving conserved amino acid positions (**Figures 4J-L and S5A**). These highly conserved binding poses in different species increase confidence in our binding model of E6R and indicate that it could potentially disrupt mammalian Sin3 HDAC complexes and associated gene regulatory functions (**Figure 1G**) by binding to the PAH2 domains of SIN3A and SIN3B and inhibiting interactions in which those domains are engaged.

### E6R reduces neuroblastoma cell invasion and tumor growth with restricted transcriptomic changes

In order to determine if E6R could produce biological responses comparable to a classical HDAC enzymatic inhibitor such as TSA in human cells, we tested its effect on neuroblastoma SK-N-BE(2)-C cell invasion, a phenotype highly sensitive to HDAC inhibition^38^. In addition, this phenotype was also reported to be sensitive to SIN3A inhibition^23^, making it a relevant model to study the effect of E6R. We observed that, similarly to TSA, E6R significantly reduced the invasion of neuroblastoma cells compared to the DMSO control (**Figure 5A**).

**Figure 5.**
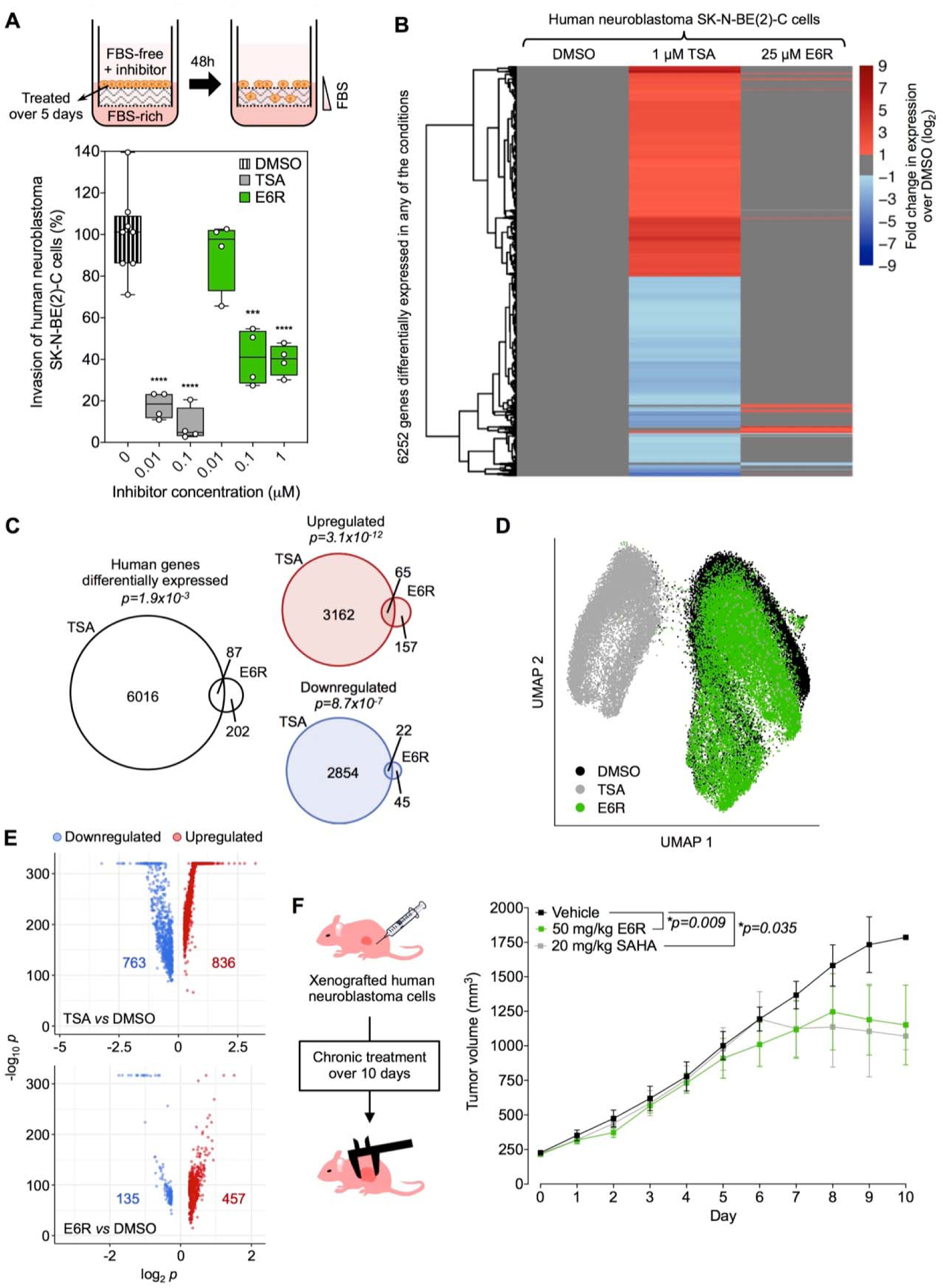
E6R reduces neuroblastoma cell invasion and tumor growth with restricted transcriptomic changes. **(A)** Box and whiskers plot showing invasion of human neuroblastoma SK-N-BE(2)-C cells following treatment with DMSO control, TSA, or E6R (n ≥ 4 biologically independent samples per group). The box extends from the 25^th^ to 75^th^ percentiles. Symbols represent independent repeats, whiskers minimum to maximum values, and bars within boxes medians. **(B)** Genes differentially expressed (RNA-seq) in human neuroblastoma SK-N-BE(2)-C cells following treatment with TSA or E6R (n = 3 biologically independent samples per group). For each condition, a row represents a gene log_2_(fold change). **(C)** Venn diagrams of the differentially expressed genes presented in (B). **(D)** Uniform manifold approximation and projection (UMAP) plot of scRNA-seq data for human neuroblastoma SK-N-BE(2)-C cells treated with TSA, E6R or DMSO control (46,751 cells total) (n = 2 biologically independent samples per group). Symbols represent individual cells. **(E)** Volcano plots showing differentially expressed genes (scRNA-seq) between cells treated with DMSO or E6R, and cells treated with DMSO or TSA from (D). **(F)** Effects of E6R and SAHA on tumor growth in mice xenografted with human neuroblastoma SK-N-BE(2)-C cells (n ≥ 5 biologically independent samples per group). Symbols represent means of replicates, error bars SEM, and lines connecting curves between data points. Statistical analyses, (A) unpaired t-test with Welch’s post-test correction relative to DMSO, and (F) paired t-test between curves of different treatments compared to the vehicle curve. Raw data are provided in **Source Data**.

To then benchmark the transcriptional effects of E6R to those of TSA, we measured associated transcriptomic changes in human neuroblastoma SK-N-BE(2)-C cells. Before conducting genome-wide analyses by RNA-seq, we established TSA and E6R working concentrations based on RT-qPCR titration experiments with three Sin3 HDAC complex-regulated genes (**Figure S6A**). The replicates of each individual condition tested by RNA-seq clustered together, showing high reproducibility of the treatments (**Figure S6B**). Remarkably, E6R modified expression of approximately 20-fold fewer genes compared to TSA (**Figures 5B and S6C**), with significant overlaps between the two conditions (**Figure 5C**).

To further assess the transcriptomic states of human neuroblastoma SK-N-BE(2)-C cells treated with DMSO, TSA or E6R, we conducted single-cell (sc) RNA-seq analyses. By performing dimensionality reduction and unbiased clustering on the resulting data, we confirmed that E6R yielded more subtle transcriptomic alterations than TSA since TSA-treated cells clustered distinctly and E6R-treated cells admixed with the DMSO control (**Figure 5D**) while perturbing expression of fewer genes (**Figure 5E**). Of the 57 genes detected in the scRNA-seq analysis from among the 65 genes upregulated by both TSA and E6R in the bulk RNA-seq approach (**Figure 5C**), 43 and 22 were upregulated by TSA and E6R, respectively. Meanwhile, of the 18 genes detected in the scRNA-seq analysis from among the 22 downregulated genes in the bulk RNA-seq study (**Figure 5C**), 13 and 5 were also downregulated by TSA and E6R, respectively (**Table S4**).

Finally, to examine the effect of E6R *in vivo*, we performed a three-arm efficacy study on immunocompromised mice xenografted with human neuroblastoma SK-N-BE(2)-C cells. We compared results to FDA-approved Vorinostat, also called suberoylanilide hydroxamic acid (SAHA), an established HDAC enzymatic inhibitor^8^ and structural analog of TSA reported to decrease neuroblastoma growth *in vivo*^48^, and tested in clinical trials (www.ClinicalTrials.gov identifiers: NCT01208454, NCT03332667). In this assay, both compounds were well tolerated by the animals and did not affect their weight over time (**Figure S6D**). E6R and SAHA both significantly reduced the growth of neuroblastoma tumors compared to the vehicle in the xenografted mice (**Figure 5F**), which illustrates the potential for medical applications of E6R. Together, these results indicate that E6R can yield biological effects similar to classical HDAC enzymatic inhibitors in human cells and mice, while inducing only a few specific transcriptomic changes.

### E6R reduces head-twitch response with restricted transcriptomic changes in the frontal cortex

Previous findings reported that repeated administration of the FDA-approved HDAC enzymatic inhibitor SAHA affects transcription of numerous plasticity-related genes in the frontal cortex (FC) and reduces head-twitch response (HTR) behavior^49,50^, a mouse behavioral proxy of human hallucinogen effects used, for example, to model sensory processing alterations observed in psychiatric conditions such as schizophrenia. These SAHA-dependent transcriptomic and behavioral responses were not observed in mice with conditional knockout of *Hdac2* (*Hdac2* cKO) in the pyramidal neurons^50^, suggesting that the effect of SAHA is dependent on functional Sin3 HDAC complexes. Based on these findings, we sought to test whether an E6R-mediated perturbation of Sin3 HDAC complexes would revert the psychedelic-induced HTR behavior (**Figure 6A**). To achieve this, we examined HTR in mice exposed to the psychedelic drug 1-(2,5-dimethoxy-4-iodophenyl)-2-aminopropane (DOI)^51^, following chronic treatment with E6R. We observed that E6R significantly reduced DOI-induced HTR in WT mice (**Figure 6B**), to the same extent as in untreated *Hdac2* cKO animals (**Figure 6C**) and without any apparent sign of toxicity (**Figure S6E**). Moreover, when the same group of *Hdac2* cKO mice was chronically treated with E6R, no additive effect was observed (**Figure 6C**), indicating that E6R requires functional HDAC complexes in the FC to reduce the HTR behavior.

**Figure 6.**
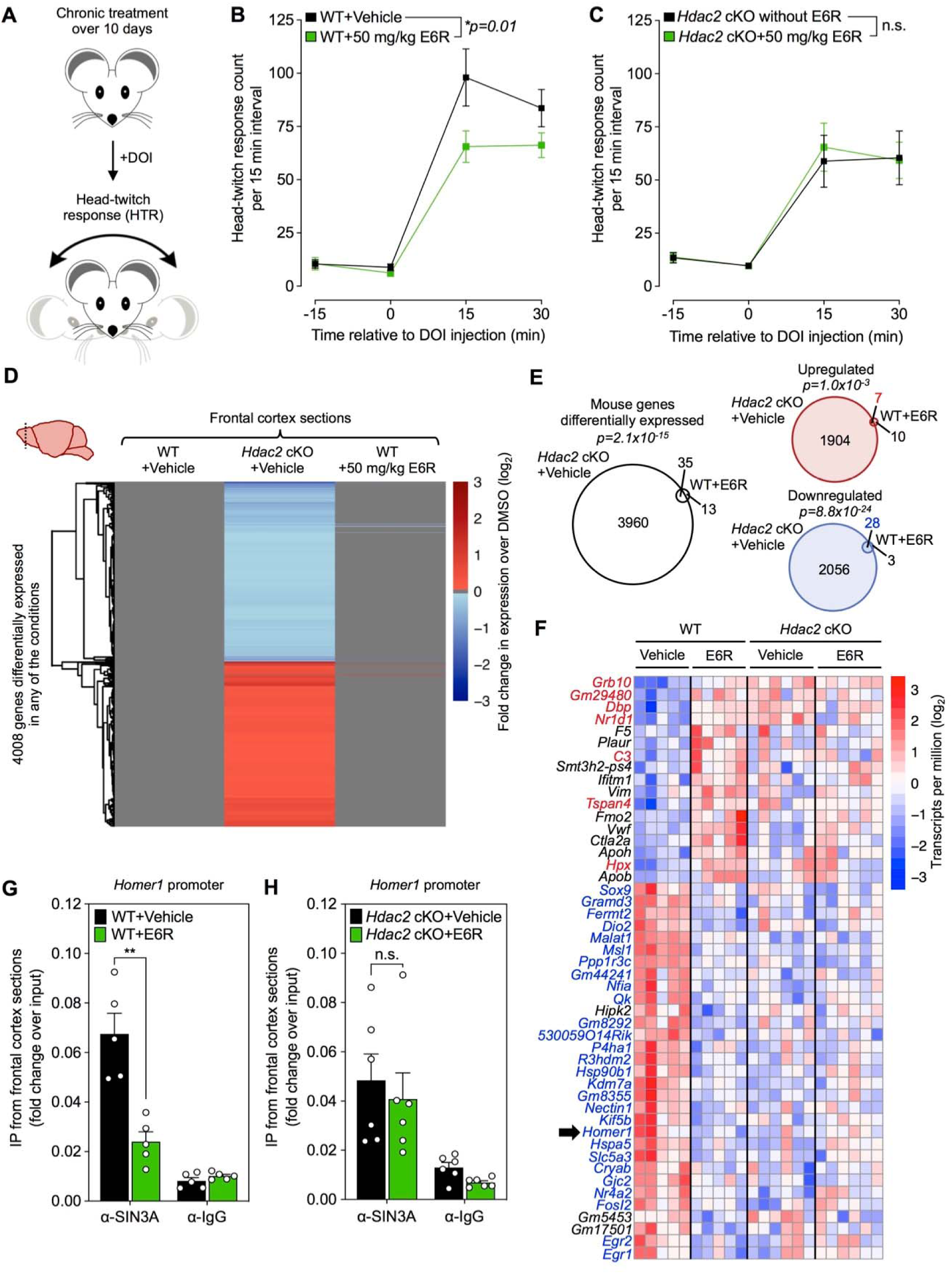
E6R reduces head-twitch response with restricted transcriptomic changes in the frontal cortex. (**A**) Principle of the psychedelic-induced HTR phenotype in mouse. **(B)** HTR phenotypes for WT mice treated chronically with E6R or vehicle (n ≥ 5 biologically independent samples per group). Symbols represent means of replicates, error bars SEM, and lines connecting curves between data points. **(C)** HTR phenotypes of *Hdac2* cKO mice before and after chronic E6R treatment (n = 5 biologically independent samples per group). Symbols represent means of replicates, error bars SEM, and lines connecting curves between data points. **(D)** Genes differentially expressed (RNA-seq) in FC sections of WT or *Hdac2* cKO mice chronically treated with E6R or vehicle, respectively (n ≥ 5 biologically independent samples per group). For each condition, a row represents a gene log_2_(fold change). **(E)** Venn diagrams for the differentially expressed genes presented in (D). The numbers of overlapping upregulated and downregulated genes are indicated in red and blue, respectively. **(F)** RNA-seq replicates for the 48 differentially expressed genes (FDR < 0.3) in the WT+E6R group compared to *Hdac2* cKO+Vehicle and *Hdac2* cKO+E6R groups using WT+Vehicle mice as references (n ≥ 5 biologically independent samples per group). Row-centered log_2_ transcripts per million (TPM) are shown and the blue/red gradients correspond to low/high expression, respectively. Upregulated and downregulated genes in both WT+E6R and *Hdac2* cKO+Vehicle groups are in red and blue fonts, respectively. Each column represents RNA-seq data for a different mouse. The black arrow indicates results for the *Homer1* gene. (**G and H**) Recruitment of SIN3A (ChIP-qPCR) at the *Homer1* locus in WT (G) or *Hdac2* cKO (H) mice treated chronically with E6R or vehicle, and compared to α-IgG negative controls (n ≥ 5 biologically independent samples per group). Symbols represent independent repeats, bars represent means, and error bars SEM. Statistical analyses, (B and C) two-way ordinary ANOVA with Bonferroni’s post-test correction, and (G and H) multiple t-tests compared to the vehicle conditions. Experiments and analyses were conducted with the same mice (D-H). Raw data are provided in **Source Data**.

To measure the impact of E6R on HDAC2-dependent transcriptional regulation in the FC, we conducted RNA-seq on FC tissue samples from WT or *Hdac2* cKO mice chronically treated with vehicle or E6R. Compared to conditional *Hdac2* deletion, treatment with E6R led to approximately 80-fold fewer transcriptomic changes in the FC. Indeed, comparing gene expression profiles of E6R-treated to vehicle-treated WT mice (five per cohort), we identified only 48 significant differentially expressed genes (**Figure 6D**). Importantly, the majority of the differentially expressed genes following E6R treatment were also altered in *Hdac2* cKO mice, suggesting that this compound only affects a subset of the HDAC2-regulated genes in the FC (**Figure 6E**). In addition, gene expression changes induced by E6R treatment were generally abolished in the *Hdac2* cKO background as compared to the WT background, suggesting that the E6R-induced transcriptomic changes rely on functional Sin3 HDAC complexes (**Figure 6F**). Together, these results are in agreement with those obtained in human neuroblastoma cells as they indicate that E6R can achieve HDAC inhibitor-like effects *in vivo* while inducing only limited and specific transcriptional changes.

Finally, to confirm that the E6R-mediated perturbations of gene regulatory functions *in vivo* result from inhibition of the Sin3 HDAC complex recruitment, we analyzed SIN3A occupancy at the *Homer1* locus in FC tissues by ChIP-qPCR. *Homer1* is a gene known to be directly regulated by a Sin3 HDAC complex in brain tissues^52^ and its expression is altered in many neuropsychiatric disorders such as schizophrenia where it is upregulated in rat models and deceased patients^53,54^. Thus, *Homer1* downregulation following treatment with E6R might potentially have beneficial therapeutic effects in this context (**Figure 6F**). ChIP-qPCR analyses confirmed that E6R significantly reduces the presence of SIN3A at the promoter of *Homer1* (**Figure 6G**). This disruption was not observed in *Hdac2* cKO mice, indicating one more time that the effect of E6R is dependent on functional HDAC complexes in the FC (**Figure 6H**). Together, those results strongly suggest that the biological activity and the selective transcriptomic effects of E6R originate from its ability to bind to the PAH2 domains of SIN3 proteins, which, in turn, only perturbs recruitment of a subset of the Sin3 HDAC sub-complexes traditionally affected through HDAC enzymatic inhibition.

## DISCUSSION

Targeting the transcriptional repression functions of the conserved Sin3 HDAC complex allowed us to identify E6R, a non-enzymatic and highly selective PPI inhibitor. Compared to classical HDAC enzymatic inhibitors, this compound induces beneficial biological effects while affecting expression of up to two orders of magnitude fewer genes by binding to the PAH2 domain of the SIN3 protein. E6R is an experimental compound which, in addition to bind to SIN3 PAH2 domains also binds to the conserved PAH1 and PAH3 domains of yeast and human SIN3 proteins according to NMR STD, DLB, and CPMG-R_2_ assays (data not shown). It will thus require further SAR-based optimizations to improve its potency and precise on-target effects for medical applications. In addition, the identification of SIN3 not only in this study, but also in a few other reports^22,23,25,26^ as an alternative target to disrupt HDAC complexes is quite remarkable and should motivate the development of alternative HDAC inhibitory strategies based on the identification of additional ligandable subunits. Further experimental tools might also be developed in cellular and animal models to characterize the effects of novel compounds, such as knockouts of *SIN3A*/*B* or other subunits of the Sin3 HDAC complex. In this context and in agreement with published studies^10^, our results also suggest that HDAC complexes can be directly involved in transcriptional activation processes, as evidenced by the decrease in basal expression levels of some direct HDAC target genes upon deletion of particular subunits or treatment with HDAC inhibitors.

Our approach combines recent advances in the characterization of PPI interfaces, development of PPI assays^39^, and systematic mapping of binary protein interactions^2^ and cellular complexes^55^ to prioritize new biological targets and assess them against diverse small-molecule libraries. Other biological complexes or pathways with a functional readout such as the SAGA transcriptional activator complex, the CCR4-NOT complex, the SARS-CoV-2 RdRp complex, or the TOR and MAP kinase pathways could also be selectively targeted following a similar strategy. These efforts to expand the druggable proteome by targeting functional, individual multimeric complexes will be supported by ultra-large virtual drug screening platforms^56^, systematic predictions of protein co-complex structures^57^ and binding partners^58^, as well as by the development of powerful structural biology technologies such as cryo-electron microscopy^59^. Notably, these technologies could also help characterizing the mechanisms of action of other compounds identified in this study.

Finally, while classic HDAC inhibitors lead to “multigenic” effects, derepressing some hundreds to thousands of genes while repressing others, HDAC complex inhibitors like E6R allow tailored, “oligogenic” effects on transcriptomes in mammalian cells and in mice. We hypothesize that this might potentially result in reduced toxicity and fewer adverse effects during treatments of disorders in which HDAC complexes play a role. Although this appears as an important advantage, it remains to be formally demonstrated that this higher selectivity can be linked to lower toxicity in animal models and patients. Overall, our results represent a major step forward in HDAC biology and epigenetics, providing a long awaited response to the poor selectivity of current epigenetic modulators.

## METHODS

### Culture of yeast cells

All yeast cells used in this study are from the *Saccharomyces cerevisiae* species and their genotypes are presented in **Table S5**. For each experiment, yeast cells from the glycerol stock were streaked onto non-selective (extract-peptone-dextrose, YPD) or desired selective solid medium, and grown for 3-5 days at 30°C to obtain fresh isolated single colonies. Except otherwise indicated, cells were cultured at 30°C under shaking conditions (200-220 rpm) in liquid non-selective (YPD) or desired selective medium until reaching the desired cell density for treatments, as described in the method details. Specific yeast growth conditions in each experiment are indicated in **Table S6**.

### Culture of human neuroblastoma SK-N-BE(2)-C cells

Cells were obtained from Kimberly Steigmaier’s lab (Dana-Farber Cancer Institute and Broad Institute of Harvard and MIT). They were previously authenticated by short tandem repeat (STR) profiling and were not contaminated by mycoplasma. Cells were maintained in cell culture flasks supplemented with 10% fetal bovine serum (FBS) and penicillin/streptomycin (PS) in Dulbecco’s Modified Eagle’s Medium (DMEM). They were maintained at a log phase in a humidified 5% CO_2_ incubator at 37°C until treatment with the different compounds as described in method details.

### Culture of human HEK293T cells

HEK293T cells were not contaminated by mycoplasma and were cultured as previously reported^39^.

For the PPI mapping experiments, cells were seeded at 6×10^4^ cells per well in 96-well flat-bottom, cell culture microplates (Greiner Bio-One, #655083), and cultured in DMEM supplemented with 10% fetal calf serum at 37°C/5% CO_2_.

For the retest of compound E6 by GPCA, HEK293T cells were seeded at a density of 3×10^5^ cells/mL/well in six-well plates (i.e. 2 mL per well) and cultured in DMEM medium supplemented with 10% FBS at 37°C/5% CO_2_.

### Media and conditions for selections of yeast cells

The media used in this study were previously described^60^. Cells were cultured as described above. For culture conditions requiring specific selection(s), synthetic complete (SC) media lacking one or several components (e.g. no uracil = SC lacking uracil or SC-URA; SC-LEU = SC lacking leucine; SC-TRP = SC lacking tryptophan) were used. Cycloheximide (CHX) or 5-fluoroorotic acid (5FOA) were added to the different media at the indicated final concentrations. When the *NatMX*, *KanMX* or *HphMX* cassette was used to delete a specific gene in the yeast genome, nourseothricin (clonnat), geneticin (G418), or hygromycin B was added to YPD or SC media to select for transformants. Conditions used in experiments involving yeast cells are detailed in **Table S6**.

### Yeast transformations

Yeast cells were transformed following a high-efficiency LiAc/salmon sperm (SS) carrier DNA/polyethylene glycol (PEG) protocol^60^. For each transformation reaction, 5 mL of log phase yeast cell culture (OD_600 nm_ ∼0.6-1) and ∼0.5 µg of high-purity plasmid, or 5-10 µg PCR-amplified marker cassette, was used. The resulting transformants were plated onto solid selective media or YPD supplemented with the proper antibiotics, depending on the plasmid or PCR-amplified marker cassette used. When antibiotics were used in the solid media, transformed cells were pre-cultured in liquid YPD (30°C) for 3 h before being plated. Transformants were picked after 3-5 days of incubation at 30°C for validations (e.g. via specific PCR amplification of the insert followed by product size confirmation with agarose gel electrophoresis and Sanger sequencing) or follow up experiments.

### Construction of yeast strains

The different gene deletion strains were generated by double homologous recombination, by transforming yeast cells with gene replacement cassettes as previously described^61–63^. Briefly, the marker cassettes were PCR-amplified with primers containing 5’-extensions (45-50 bases) directly adjacent to them, and homologous to the promoters or terminators of the targeted genomic loci. The resulting mutants were selected as described above, and single colonies picked and purified before lysing cells and validating the gene deletion/marker cassette by PCR amplification followed by agarose gel electrophoresis using specific primer pairs. MaV208 was generated from MaV108^33^ by disruption of two drug exporter genes using the *KanMX* gene cassette^64^, and the *HIS3* marker. First, MaV108 was transformed with a DNA fragment that was amplified by PCR reaction using a plasmid pLexA (Clontech) as a template for *HIS3* marker. Both ends of the PCR product included a short region homologous to *PDR5.* Transformed yeast cells were plated on SC plates lacking histidine (SC-HIS) and used to select a new strain named MaV118. This new yeast strain was further transformed with a DNA fragment that was amplified by PCR reaction using the plasmid pUG6 as a template for the *KanMX* gene cassette^64^. Both ends of the PCR product included a short region homologous to *KanMX.* Transformed cells were plated on YPD plates supplemented with 200 mg/L G418, and a new strain MaV208 was selected from the resistant colonies. For other strains, *NatMX, KanMX,* and *HphMX* cassettes were PCR-amplified from plasmids p4339 (gift from Charles Boone, University of Toronto), pRS400, and bRA89 (gift from James E. Haber, Brandeis University), respectively, with primers specific to the TEF promoter and TEF terminator. For the *WT+GAL4* control condition, WT MaV208 yeast cells were transformed with a pDEST-DB-scGal4 plasmid^39^ to induce expression of the Gal4 transcription factor and activation of *SPAL10::URA3.* For JOY134, WT MaV208 yeast cells were transformed with the pAR128 plasmid carrying *UME6* (pPL5920-*UME6*: *CEN Amp^R^ LEU2 UME6).* For JOY128 *((spal10::ura3)Δ::HphMX@ura3),* the *SPAL10::URA3* region corresponding to the Gal4 binding sites, the *URS* motif from *SPO13*^33^, the *URA3* reporter gene, and 53 bases of the *URA3* terminator, was replaced by the hph marker. The HoY013 strain was generated by inserting the PCR-amplified *SPAL10::URA3* reporter system from MaV103 genomic DNA, at the *ura3* locus of the Y8800 yeast strain. Y8800 cells were first transformed with a pDEST-DB-scGal4 plasmid before being transformed with the *SPAL10::URA3* cassette. Transformants were selected on SC-LEU-URA solid media. After purifying single colonies, the genomic insertion of *SPAL10::URA3* was confirmed by PCR with specific primers, and the pDEST-DB-scGal4 was then shuffled out by growing HoY013 cells into complete YPD medium. JOY200 was constructed by deleting *PDR5* and *SNQ2* efflux pumps in HoY013 as explained above.

### Spot tests and lawns of yeast cells

When cells reached OD_600 nm_ ∼0.6-1 in ∼1mL of the proper liquid medium, population densities were evaluated using counting grid microscope slides. They were then centrifuged, supernatants discarded and pellets washed twice with double-distilled water. The resulting pellets were then diluted in different volumes of water to obtain similar population densities throughout the samples. From these starting suspensions, serial dilutions (1:10) were made for every strain, and ∼8-10 µL of culture were spotted for each dilution, on the indicated solid agar media. Plates were then incubated according to the indicated conditions. For plating cells, the washed pellets were resuspended in water at the desired density, and spread out on the agar surface using sterile glass beads.

### Compound treatment and RNA extraction from yeast cells

After yeast cells reached OD_600 nm_ ∼0.5 in 0.8-1 mL of YPD, they were grown for two more hours at room temperature before being centrifuged to remove supernatant and freezing pellets on dry ice. For compound treatments, yeast cells at OD_600 nm_ ∼0.5 in YPD were incubated at room temperature for two hours with DMSO (0.7-1.6%) or the desired compound (from 25 mM stock in 100% DMSO) in an 2 mL-tube for a final volume of 0.8-1 mL in YPD, before being centrifuged, washed and frozen as described above. Yeast cells were then harvested for RNA extraction using a RiboPure RNA Purification Kit (Invitrogen, cat#AM1926), which involves mechanical cell wall disruption, phenol extraction of the lysate, and RNA purification using glass fiber, and filter-based RNA purification columns. RNA concentration was measured for individually purified RNA samples using a NanoDrop spectrophotometer. A total of 1 µg RNA from each sample was used for reverse transcription following a published procedure^65^.

### RT-qPCR in yeast cells and human neuroblastoma SK-N-BE(2)-C cells

The RT-qPCR protocol used in this study was described elsewhere^65^. Primers used in this study are presented in **Table S7**. Approximately 1 µg RNA per sample was used for reverse transcription. Using a thermocycler, RNA was denatured in the presence of 0.2 µg of 18 mer oligo-dT by heating the plate at 65°C for 10 min. The plate was then immediately cooled down on ice to anneal oligo-dT on the poly-A tail of mRNA. A master mix of AffinityScript Multiple Temperature Reverse Transcriptase (Agilent, cat#600107) was prepared following the manufacturer’s protocol, and mixed with RNA samples previously annealed with the oligo-dT. The plate was incubated at 42°C for 2 h to generate cDNAs from mRNA templates. The reverse transcriptase was then denatured by heating the plate at 70°C for 15 min. The resulting cDNA was diluted (1:20) in sterile, double-distilled water, and stored at -20°C until used for qPCR. To measure gene expression, the diluted cDNA was mixed with 5 µL PowerUp SYBR Green master mix (Applied Biosystems, cat#A25743), and the mix of primers (forward+reverse: 0.3 µM final concentration) for a total volume of 10 µL. Two to three technical replicates were generated per cDNA sample in the 384-well PCR plate (Applied Biosystems, cat#4343370). The plate was tightly sealed with an optical adhesive film (Applied Biosystems, cat#4311971). Quantitative PCR was conducted with a QuantStudio real-time PCR system or an ABI Prism system (both from Applied Biosystems), with the following experimental settings: 50°C/2 min, 95°C/10 min, and 40 cycles of 95°C/15 s and 60°C/30 s. Dissociation curves were checked for each PCR product to assess the specificity of PCR amplicons and products were sequenced to confirm target genes.

### Calculating relative expression from RT-qPCR measurements

Gene expression measured by RT-qPCR was calculated using the following equation:

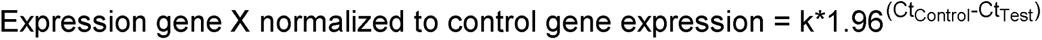

Where k is an arbitrary multiplier (k = 1000 in this study), 1.96 is the PCR amplification efficiency constant, Ct is the cycle threshold, Ct_Control_ is the Ct value for the housekeeping gene used as control for normalizations *(UBC6* in yeast^66^; *RPS11* in human cells^67^), and Ct_Test_ is the Ct value for the tested gene. Replicates corresponding to DMSO treatments or WT yeast cells were averaged and other data points normalized using those DMSO or WT values as references (i.e. expression relative to DMSO or WT). *TAF10* in yeast or *SNRPD3* in human cells were used as genes not known to be regulated by an HDAC complex (HDAC-unrelated genes). Two to three technical replicates were collected and averaged for each biologically independent sample per group. In the rare cases where the value of a technical replicate was undetermined or gave the same result as a control empty well in the plate, it was excluded from the calculation of the average.

### Agar diffusion assay for small molecule selection

Yeast cells were seeded on the appropriate solid media (dried for at least three days at room temperature with lid on) as described above, at the cell density per plate indicated in **Table S6**. After ∼30 min, the compounds to be tested (dissolved in 100% DMSO) were spotted (1 µL for the retests of 59 representative small molecules and 4 µL for the retests of prioritized compounds) at the indicated concentration(s) onto the seeded cells, to allow diffusion and formation of circular concentration gradients around the original spots (see **Note S1**).

### Constructing plasmids expressing Sin3/Rpd3L HDAC complex subunits

Plasmid pAR124 was generated by PCR amplifying the *SIN3* locus (chrXV: 316440-321724) from *S. cerevisiae* S288c genomic DNA (Novagen, cat #69240) with tailed primers containing Gateway attB sites and cloning this amplicon into pAR107. Plasmid pAR107 was generated by digesting pRS415 with the restriction enzymes SmaI and SalI (New England Biolabs), PCR amplifying the Gateway cassette from pQZ213 and assembling both products by gap repair in the yeast strain BY4733. Plasmid pAR128 was generated by digesting pPL5920 with MluI-HF and NsiI-HF (New England Biolabs), PCR amplifying the *UME6* locus (chrIV: 864920-868020) from *S. cerevisiae* S288c genomic DNA (Novagen, cat #69240), and ligating both products with the T4 DNA Ligase (New England Biolabs). Plasmid YEplac181-Sin3-(HA)_3_ *(2µ Amp^R^ LEU2 SIN3- (HA)_3_)* used for ChIP experiments was a gift from David J. Stillman (University of Utah) originally obtained from Kevin Struhl’s lab (Harvard Medical School).

### High-throughput chemical screening with the *URS-URA3* reporter assay

Using the agar diffusion assay described above, the yeast Sin3/Rpd3L HDAC complex was interrogated against 52,234 candidate small molecules (**Table S1**). JOY134 yeast cells were seeded at a density of ∼1-5×10^9^ cells per plate on solid SC-LEU-URA medium, poured into Nunc^TM^ OmniTray™ single-well plates. The media were dried for at least three days at room temperature (with lid on) before seeding yeast cells, and plates with seeded cells were dried approximately 30 min in a sterile airflow hood before starting compound spotting. Using a custom designed compound transfer workstation at the ICCB-Longwood Screening Facility at Harvard Medical School, 200 nL of compounds dissolved in DMSO were pinned from each well of a 384-well library plate onto duplicate assay plates. Concentrations of the used stock libraries are indicated in **Table S1**. In addition, 200 nL of DMSO and TSA (at a concentration of 75 mM in 100% DMSO) were manually spotted onto each assay plate. After five days of incubation at 30°C, each duplicated plate was manually scored by comparing the rings of growth from candidate compounds to the DMSO negative controls. For reconfirmation, potential hits from the primary screen were cherry-picked and 2 µL/compound were spotted onto JOY134 yeast cells seeded on large SC-LEU-URA square (24 cm) plates (∼5×10^9^ seeded cells per plate). Primary hits that conferred Ura_+_ phenotypes (rings of growth) in this retest experiment (275 distinct in total) were then clustered based on their structural similarities.

### Clustering and selecting retested compounds from screening

Among the 275 distinct compounds that conferred reproducible Ura_+_ phenotypes in JOY134 yeast cells (i.e. rings of growth on SC-LEU-URA), nine pan assay interference compounds (PAINS) were flagged using a pattern-matching algorithm with a published list of 481 chemical groups as input^68^. Similarities between structures were then automatically calculated based on functional class fingerprints (FCFP4) with distance to closest using the Tanimoto coefficient. An average cluster size of five compounds was selected, which outputted 56 different groups. The chemical structures within those clusters were then manually inspected and 59 representative small molecules were selected and purchased (dry powders) for retest in a second yeast strain, as described in the main text (in some cases, two to three representative compounds per individual cluster were selected). The seven prioritized compounds from screening were obtained from ChemDiv (www.chemdiv.com), under the following references (ChemDiv 6 library): G857-1026 (D2), C200-3465 (E2), 2969-0361 (E5), 5594-3013 (D4), C720-0661 (E6), E910-0075 (G6), and G515-0309 (B8). Their purities were confirmed to be >90-95% by liquid chromatography-mass-spectrometry (LC-MS).

### Treatment of human neuroblastoma SK-N-BE(2)-C cells with compounds

Cells were cultured as described above. For testing the effect of the compounds on gene expression, approximately 0.1 million cells were seeded in each well of 24-well cell culture plates, with 1 mL 10% FBS+PS+DMEM. Twenty-four hours after seeding, cells were exposed to the compounds or DMSO controls (0.4% for testing the seven confirmed small molecules or 0.8% for the titration and RNA-seq experiments) for 16 h in a humidified 5% CO_2_ incubator at 37°C, before harvesting them.

### RNA extraction from human neuroblastoma SK-N-BE(2)-C cells

Cells were harvested for RNA extraction using Nucleospin 96 RNA (Macherey-Nagel, cat#740709.4) following the manufacturer’s protocol. Briefly, supernatant was removed, and the cells were gently washed with 1x phosphate-buffered saline (PBS). 300 µL lysis buffer were added to each well of 24-well cell culture plates. Roughly 0.1 million cells were used for harvesting each RNA sample. RNA concentration was measured for all purified RNA samples using a nanodrop spectrophotometer and RT-qPCR was conducted as described above.

### Assessment of HDAC enzymatic activity

The HDAC-Glo I/II^TM^ screening system (Promega, cat#G6430)^69^ was adapted to measure the effect of compounds on the enzymatic activity of human and yeast HDACs (see **Note S2**). Purification of recombinant, human HDAC1 was performed from HEK293T cells stably over-expressing an N-terminally FLAG-tagged HDAC1 enzyme, as previously described^70^. FLAG-HDAC1 was eluted from anti-FLAG beads before running the assay, using a FLAG peptide. Nuclear extracts from human HeLa cells were provided by the supplier in the assay kit. HDAC enzymatic activity was measured following the manufacturer’s protocol. Briefly, linear ranges of the assay were determined by measuring luminescence signals, in a serial dilution manner, for semi-purified human HDAC1, or live yeast cells, diluted in the HDAC-Glo I/II buffer. For HeLa nuclear extracts, the manufacturer provided the working concentration in the assay kit protocol. For the different HDAC sources, dilution factors corresponding to signals in the linear ranges were used to work under optimal conditions of sensitivity (∼5-10x dilution for FLAG-HDAC1; ∼10^6^-10^7^ yeast cells/reaction). To test the effect of compounds on HDAC enzymatic activity, equal volumes of HDAC sources (5 µL), and inhibitors (5 µL) were added to the different wells of a white, flat bottom 384-well plate. The plate was then briefly centrifuged with a tabletop centrifuge, and gently shaken for 1 min at 700 rpm. The plate was tightly sealed to prevent evaporation, and incubated at room temperature for 1 h. Following this incubation, 10 µL of luciferase substrate were added to each well. The 384-well plate was then briefly centrifuged, and shaken for 1 min at 700 rpm before being tightly sealed, and incubated for 15 min at room temperature. Luminescence signals were measured using a TriStar luminometer from Berthold Technologies, with 1 s integration time per sample.

### Mass-spectrometry of FLAG-HDAC1 purified from HEK293T cells

The FLAG-HDAC1 enzymes were immunoprecipitated as described above, using anti-FLAG beads. Beads were treated with DMSO for 1 h, in triplicate, before being washed, and FLAG-HDAC1 enzymes eluted using the FLAG peptide. Proteins from three different replicates were precipitated by methanol/chloroform and digested with trypsin overnight at 37°C, in 50 mM NH_4_HCO_3_ pH 8.0. Peptides were quantified using a colorimetric peptide assay (Thermo Fisher Scientifc). Peptides (1 µg) were dissolved in solvent A (0.1% TFA in 2% ACN), directly loaded onto reversed-phase pre-column (Acclaim PepMap 100, Thermo Scientific) and eluted in backflush mode. Peptide separation was performed using a reversed-phase analytical column (Acclaim PepMap RSLC, 0.075 x 250 mm, Thermo Scientific) with a linear gradient of 4%-27.5% solvent B (0.1% FA in 98% ACN) for 35 min, 27.5%-50% solvent B for 10 min, 50%-95% solvent B for 10 min and holding at 95% for the last 5 min at a constant flow rate of 300 nL/min on an Ultimate 3000 RSLC system. The peptides were analyzed by an Orbitrap Fusion Lumos Tribrid mass spectrometer (ThermoFisher Scientific). The peptides were subjected to NSI source followed by tandem mass spectrometry (MS/MS) in Fusion Lumos coupled online to the nano-LC. Intact peptides were detected in the Orbitrap at a resolution of 120,000. Peptides were selected for MS/MS using HCD setting at 30, and ion fragments were detected in the Ion Trap. A data-dependent procedure that alternated between one MS scan followed by MS/MS scans was applied for 3 s for ions above a threshold ion count of 5.0×10^3^ in the MS survey scan with 40.0 s dynamic exclusion. The electrospray voltage applied was 2.1 kV. MS1 spectra were obtained with an AGC target of 4×10^5^ ions and a maximum injection time of 50 ms, and MS2 spectra were acquired with an AGC target of 5×10^4^ ions and a maximum injection time of 50 ms. For MS scans, the m/z scan range was 375 to 1800. The resulting MS/MS data was processed using Sequest HT search engine within Proteome Discoverer 2.5 against a human protein database obtained from Uniprot (87,489 entries). Trypsin was specified as cleavage enzyme allowing up to 2 missed cleavages, 4 modifications per peptide and up to five charges. Mass error was set to 10 ppm for precursor ions and 0.6 Da for fragment ions. Oxidation on Met and N-terminal acetylation were considered as variable modifications. False discovery rate (FDR) was assessed using Percolator and thresholds for protein, peptide and modification site were specified at 1%. Abundance ratios were calculated by Label Free Quantification (LFQ) of the precursor intensities within Proteome Discoverer 2.5. Proteins were considered only when they were confidently identified in at least two out of three replicates. Known HDAC1, HDAC2, SIN3A and SIN3B interactors according to mentha^71^ (downloaded on Aug 28, 2017) were checked and information on yeast *S. cerevisiae* orthologs was added according to the PANTHER database (version 16.0; queried on Jan 28, 2022) to construct **Table S3**. Orthologs marked as “LDO” (least diverged ortholog) or “O” (more diverged ortholog) in PANTHER were considered in **Table S3**.

### Cloning open reading frames (ORFs) into expression plasmids

ORFs corresponding to full-length subunits of the Sin3/Rpd3L HDAC complex were PCR-amplified from yeast genomic DNA, and cloned into Gateway entry vectors, pDONR223. Following bacteria transformations, single colonies were picked, and the quality of cloning was checked for every ORF by bi-directional Sanger DNA sequencing. They were then introduced into the different assay-specific expression vectors (GPCA, KISS) by LR clonase-mediated Gateway reactions (Life Technologies). LR reaction products were subsequently transformed into *E. coli* DH5α competent cells and grown for 24 h on ampicillin-containing TFB medium. Plasmid DNA was extracted using a NucleoSpin 96 Plasmid kit from Macherey-Nagel. After PCR-amplification of the cloned ORFs from purified plasmid DNAs with plasmid-specific primers, the size of each DNA amplicon was examined by agarose gel electrophoresis. For hsPRS-v2 and hsRRS-v2 pairs, the published N_1_N_2_ GPCA constructions were used^39^. For GPCA, ORFs coding for the Sin3/Rpd3L HDAC complex subunits were cloned into the GPCA-N1 and GPCA-N2 destination plasmids, which allowed expression of the tested proteins, X and Y, as N-terminal fusions of the two luciferase fragments, Luc1 (N_1_) and Luc2 (N_2_). For KISS, ORFs coding for proteins found in PPIs identified by GPCA were cloned into the KISS-C1 and N2-KISS destination plasmids to allow expression of the tested proteins, X and Y, as C- and N-terminal fusions, i.e. X-C_1_/N_2_-Y versus Y-C_1_/N_2_-X.

### Culture and transfection of HEK293T cells for GPCA

Cells were cultured as described above. Twenty-four hours following seeding, cells were transfected with 100 ng of each GPCA plasmid (GPCA-N1 and GPCA-N2) using linear polyethylenimine (PEI) to co-express the protein pairs fused with complementary luciferase fragments, Luc1 (N1) and Luc2 (N2). The DNA/PEI ratio used for transfection was 1:3 (mass:mass). GPCA vectors carry the human cytomegalovirus (CMV) promoter and are maintained as high copy numbers with the human virus SV40 replication origin in mammalian cells. Twenty-four hours after DNA transfection, the cell culture medium was removed, cells were gently washed with 150 µL of pre-warmed 1x PBS, 40 µL of lysis buffer were then added per well, and cell lysis was performed under vigorous shaking of the plate for 20 min at 900 rpm. Luminescence was measured for each N_1_-X/N_2_-Y pairwise combination by auto-injecting 50 µL Renilla luciferase substrate (Renilla Luciferase Assay system, catalog No. E2820, Promega) per well and integrating light output for 4 s using a TriStar luminometer (Berthold). The stock solution of PEI HCl (PEI MAX 40000; Polysciences Inc; cat#24765) was prepared according to the manufacturer’s instructions. Briefly, 200 mg of PEI powder were added to 170 mL of water, stirred until complete dissolution, and pH was adjusted to 7 with 1 M NaOH. Water was added to obtain a final concentration of 1 mg/mL, and the stock solution was filtered through a 0.22 µm membrane.

### Mapping binary PPIs with GPCA

GPCA (N_1_N_2_ version) was implemented as previously reported^39^. To work under conditions where GPCA maximizes detection of true positive PPIs while minimizing recovery of random protein pairs, a benchmarking experiment using the 60 hsPRS-v2 and 78 hsRRS-v2 pairs previously published^39^ was run in parallel to the all-by-all screening of the Sin3/Rpd3L HDAC complex PPIs. A luminescence threshold corresponding to an FDR of 1.0% hsRRS-v2 pairs was calculated from raw data and used to score interactions. This 1.0% hsRRS-v2 detection cutoff was determined using R quantile function after log_2_ transformations of the raw luminescence signals of hsRRS-v2 pairs, which resulted in a calculated value of 38240.50 units. All protein pairs with luminescence signals above that threshold were scored positive. A p value of 4.01×10^-10^ was calculated for the set of positive hsPRS-v2 pairs over the hsRRS-v2 random controls using R functions “Wilcox. test”. The mapping of binary PPIs of the Sin3/Rpd3L HDAC complex was conducted by pairwise testing all 12 subunits against each other’s, ultimately assessing every possible N_1_-X/N_2_-Y and N_1_-Y/N_2_-X pairwise combination. Positions of the tested pairs were randomized in the culture plates. Luminescence was measured for all 144 possible pairwise combinations (i.e. search space of 12 x 12 proteins), along with the 138 protein pairs from hsPRS-v2/hsRRS-v2. In this all-by-all screen, every protein pair scoring above the calculated 1.0% hsRRS-v2 detection cutoff was defined as positive, and associated to an empirical p value = 0.01.

### Processing published hsPRS-v2 and hsRRS-v2 data

In order to compare published data to results from this study obtained under the same conditions, raw luminescence values for N_1_N_2_ GPCA were first extracted from the literature^39^. The hsPRS-v2 recovery rate for these experiments was then recalculated using a theoretical hsRRS-v2 detection threshold of exactly 1.0% as described above, which resulted in a calculated value of 62745.18 units for the published data. Every hsPRS-v2 pair scoring above this signal was considered positive. Published results were then compared to those obtained here, for each individual protein pair, to evaluate the reproducibility of GPCA between different studies. KISS data were treated as described below.

### Retesting PPIs from GPCA with KISS orthogonal assay

KISS (C_1_N_2_ version) was implemented as previously reported^39,72^. HEK293T cells were cultured as described above. Cells were transfected with bait, prey and reporter plasmids (KISS-C1 and N2-KISS) corresponding to empty controls (unfused gp130 tag or unfused TYK2 C-terminal fragment tag) or Sin3/Rpd3L HDAC complex interacting proteins (both orientations were tested, i.e. X-Y and Y-X) initially identified by GPCA, applying a standard calcium phosphate transfection method. Luciferase activity was measured 48 h after transfection using the Luciferase Assay System kit (Promega) on a Enspire luminometer (Perkin-Elmer). The average of six culture wells was used. A normalized luminescence ratio (NLR) cutoff corresponding to exactly 1.0% hsRRS-v2 pairs scored positive was applied to the published data^39^. This calculated cutoff of 3.81 was used as reference to validate Sin3/Rpd3L HDAC complex interactions from GPCA. A p value of 4.51×10^-7^ was calculated, using the recalibrated published data^39^, for the set of positive hsPRS-v2 pairs over the hsRRS-v2 random controls using R functions “Wilcox.test”. The luciferase ratio obtained for each Sin3/Rpd3L HDAC bait-prey protein pair versus that obtained for the combination of the same bait with a negative control prey (unfused gp130), and versus that obtained for the combination of the same prey with a negative control bait (unfused TYK2 C-terminal fragment) were evaluated against the reference NLR corresponding to 1.0% hsRRS-v2 pair detection. NLR was calculated as follows, for each protein pair, X-Y:

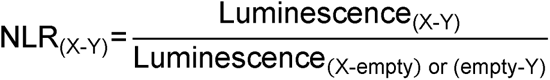

An interaction pair was scored positive when both NLR, using X-empty or empty-Y, exceeded the reference cutoff for either of the two configurations tested, corresponding to empirical p values = 0.01. Between the two NLR values of each positive pair and tested orientation, the smallest was used to construct **Figure S2D**.

### Testing and confirming binary PPI inhibition by GPCA

The ability of compounds to inhibit binary PPIs in the Sin3/Rpd3L HDAC complex was tested by GPCA in a high-throughput screening format. Interactions identified in the mapping at 1.0% hsRRS-v2 detection cutoff were reconstituted by expressing protein pairs in the exact same orientations (a single orientation per PPI was used to test compounds). Briefly, the ten binary PPIs of the Sin3/Rpd3L HDAC complex were tested against DMSO controls (n ≥ 7) (1% of the final volume), TSA, TRX, and the seven confirmed small molecules. HEK293T cells were transfected with 100 ng of each GPCA plasmid expressing either protein X, or protein Y, as described above. Twenty-four hours after DNA transfection, cells were treated with 10 µM of compounds, or equivalent DMSO volumes, and incubated for 1 h at 37°C. The luminescence signal for each PPI was measured by averaging the different DMSO replicates, and calculating the standard deviation of DMSO values for each interaction. For each PPI, an individual data point for every compound tested was obtained and this value was compared to the averaged DMSO value. For each interaction, a particular compound was scored as a PPI inhibitor when it decreased its luminescence signal by at least three standard deviations compared to the corresponding averaged DMSO value.

The independent retest of compound E6 against the N_1_-Sin3/N_2_-Ume6 interaction and the full-length *Gaussia princeps* luciferase was done in the linear range of signal. Twenty-four hours following seeding as described above, cells were transfected with 2 µg of each subunit-expressing plasmid (i.e. N_1_-Sin3, N_2_-Ume6), or 1 µg of the full-length *Gaussia princeps* luciferase-expressing plasmid using linear PEI. The mass/mass DNA/PEI ratio used for transfection was 1:3. Twenty-four hours post-transfection, cells were trypsinized and resuspended into the culture medium at a density of 3×10^5^ cells/mL before being seeded at a density of 3.5×10^4^ cells/well (i.e. 100 µL per well) in 96-well, flat-bottom microplates (Greiner Bio-One, cat#655083). After 6 h, cells were treated with the indicated concentrations of the compound (stock concentration of 25 mM in 100% DMSO), for a total volume of 1% DMSO per well (three replicates for each concentration and 11 replicates for DMSO alone). Plates were then incubated for 16 h at 37°C. The medium was removed and the lysis was implemented as described above. 50 µL/well of the Renilla luciferase substrate (Renilla Luciferase Assay system, cat#E2820, Promega) were then added and luminescence was read approximately two minutes after injection using a Centro XS^3^ LB960 plate reader (3 s integration). For each plate, signal was measured twice and the first measurement was used for calculations. Results were normalized by averaging raw luminescence values from DMSO wells and dividing each data point by this averaged value.

### ChIP-qPCR and ChIP-seq experiments in yeast cells

ChIP procedures in yeast were adapted from previously established protocols, with minor modifications^73–75^. Primers used in this study are presented in **Table S7**. Briefly, JOY116 *(sin3Δ)* yeast cells transformed with the YEplac181-Sin3-(HA)_3_ plasmid were cultured in liquid SC-LEU, until OD_600 nm_ ∼0.7. Then, DMSO (0.6% of total volume), 400 µM E6, or 400 µM TSA was added to 2×10^9^ yeast cells, and the resulting suspensions were incubated for 2 h at 30°C (200 rpm), and subsequently treated with 1% formaldehyde for 20 min at 4°C for cross-linking. Then, 125 mM glycine was applied for 5 min at room temperature to quench cross-linking. Yeast cells were centrifuged (10 min at 1,000 g), and the resulting pellets were washed twice with 50 mL ice-cold PBS. Yeast pellets were resuspended in 500 µL of ChIP lysis buffer (50 mM HEPES-KOH pH 7.5, 140 mM NaCl, 1 mM EDTA, 1% Triton X-100, 0.1% sodium deoxycholate, 0.1% SDS, and yeast protease inhibitors). Next, 500 µL of the yeast suspensions in lysis buffer were mixed with 500 µL of zirconia beads in a screw-top microcentrifuge tube. For cell lysis, the tubes were vortexed at maximum speed for 15 min at 4°C. Cell lysates were collected by punching a hole on the bottom of the screw-cap tube, with a sterile needle, and centrifuging the lysates into new Eppendorf tubes. Then, 300 µL of lysates were added into 1.5 mL Bioruptor Pico microtubes with caps (Diagenode, cat#C30010016), and subsequent sonication was conducted with a Bioruptor Pico sonication device, in the following conditions: three cycles of 30 s on and 30 s off at 4°C (sizes checked using a 1.5% agarose gel after reverse cross-linking). The sonicated lysates were cleared by 30 min centrifugation at maximum speed at 4°C. The resulting cleared lysates with sheared chromatin were saved, snap freezed, and stored at -80°C until use. Next, 10 µL cleared lysates were reverse-cross linked overnight at 65°C in the presence of proteinase K. Phenol-chloroform extractions and EtOH precipitations were then applied to purify the sheared DNA. DNA agarose gel running and DNA concentration measurements with a NanoDrop spectrophotometer were finally conducted to assess the size and yield of the sheared DNA, respectively. For the pull-downs, 25 µg of sheared chromatin from each sample were diluted (1:10) in ice-cold dilution buffer (1% Triton X-100, 2 mM EDTA, 20 mM Tris-HCl pH 8, 150 mM NaCl, and yeast protease inhibitors), and mixed with 10 µg ChIP-grade anti-HA (Abcam, Cambridge, MA, USA, cat#ab9110)^76^, or isotype control anti-IgG (Abcam, Cambridge, MA, USA, cat#ab171870) antibodies. Then, 50 µL of pre-washed Dynabeads Protein G for Immunoprecipitation (ThermoFisher, cat#10003D) were added to each individual sample. Chromatin immunoprecipitation was performed by incubating samples at 4°C overnight, under gentle shaking (300 rpm). Then, the Dynabeads were washed three times with 1 mL ice-cold wash buffer (0.1% SDS, 1% Triton X-100, 2 mM EDTA, 20 mM Tris-HCl pH 8, 150 mM NaCl), and once with 1 mL ice-cold final wash buffer (0.1% SDS, 1% Triton X-100, 2 mM EDTA pH 8, 20 mM Tris-HCl pH 8, 500 mM NaCl), using a magnetic rack. DNA was eluted with 120 µL elution buffer (1% SDS, 100 mM NaHCO_3_), by incubating samples at 30°C for 15 min. For input, 10 µL of sheared cell lysates were added to 110 µL of elution buffer. Then, 4 µL proteinase K were added to the eluted DNA or inputs, and reverse-cross linking was achieved by incubating samples overnight at 65°C. After overnight incubation, phenol-chloroform extractions and EtOH precipitations were conducted to purify ChIP-DNA. The ChIP-DNA was eluted in a total volume of 200 µL double-distilled water, and then used for ChIP-qPCR or ChIP-seq analyses. Briefly, for ChIP-qPCR, 4 µL of eluted ChIP-DNA were mixed with 5 µL of PowerUp SYBR Green master mix (Applied Biosystems, cat#A25743), and 1 µL of the primer mix (forward+reverse: 3 µM final concentration), for a total volume of 10 µL. Two technical replicates were generated for each DNA sample from two or three ChIP replicates (independent pull-downs). ChIP-DNA samples were then transferred into the 384-well skirted PCR plate (Applied Biosystems, cat#4343370). The plate was tightly sealed with an optical adhesive film (Applied Biosystems, cat#4311971). qPCR was conducted with the experimental settings described above. Dissociation curves were checked for each PCR product to assess the specificity of PCR amplicons.

### ChIP-seq library preparation

ChIP-seq libraries were prepared using Swift S2 Acel reagents on a Beckman Coulter Biomek i7 liquid handling platform from approximately 1 ng of DNA according to manufacturer’s protocol and 14 cycles of PCR amplification. Finished sequencing libraries were quantified by Qubit fluorometer and Agilent TapeStation 2200. Library pooling and indexing was evaluated with shallow sequencing on an Illumina MiSeq. Subsequently, libraries were sequenced on a NovaSeq targeting 40 million 100 bp read pairs by the Molecular Biology Core Facility at Dana-Farber Cancer Institute.

### ChIP-seq analyses

ChIP-seq tracks were visualized using the Integrative Genomics Viewer (IGV) v.2.8.10 from the Broad Institute^77^. ChIP-seq reads were mapped to the yeast *S. cerevisiae* genome (R64-1-1) with bwa v.0.7.8-r455 (http://bio-bwa.sourceforge.net/) using the mem algorithm. Peaks were called with MACS2 v.2.2.7.1^78^ using input sequencing data as a control (n = 2 per condition). Genomic locations (yeast strain S288C) of all ORFs were downloaded from the *Saccharomyces* Genome Database (SGD) (http://www.yeastgenome.org) in 2019. Genomic locations of the summits were obtained from “abs_summit” column in the MACS output files. We consider a gene is occupied by Sin3 in the ChIP-seq experiments, if the start of the gene’s ORF (i.e. ATG) is within 600 bp downstream of a summit of the consensus peaks from the two repeats, and if fold enrichment over input ≥1.5. Noticeably, *IME2* is not present in this list due to peaks positioned above 600 bp upstream from the ATG. We consider that Sin3 occupancy is reduced when (fold enrichment DMSO)/(fold enrichment treatment) ≥1.5. The p-values (two-sided Fisher’s exact test) of the overlaps (Venn diagram) and bar graphs were calculated using python 2.7.10 function “scipy.stats.fisher_exact”. A list of *URS*-containing genes was obtained by searching the MotifMap database (http://motifmap.ics.uci.edu/, Nov. 2019)^42^ using *UME6* motifs Harbison_61 (“TAGCCGCCS”), and M01503 (“NNNNWNGGCGGCWAHHNNNN“) at FDR cutoff 0.1. If the start of an ORF is within 600 bp downstream of the motif, the gene corresponding to the ORF is considered as an *URS*-containing gene. *URA3* was manually added to this list since the yeast strains used in this study contain the synthetic *SPAL10::URA3* reporter gene. To get a list of Ume6-bound genes, we downloaded GSE110681_Ume6_ChIP-exo5.0_peaks.gff.gz, and GSE110681_Ume6_ChIP-seq_peaks.gff.gz from the NCBI GEO website under accession GSE110681^43^. Using the genomic locations of the peaks extracted from the gff files, we identified all the genes with ORFs whose start (i.e. ATG) is located within 600 bp downstream of the peaks. To make the list more stringent, only the genes identified using both ChIP-exo5.0 and ChIP-seq gff files were considered as Ume6-bound genes.

### Cloning yeast Sin3 and Ume6 and human SIN3A and SIN3B protein domains

A DNA sequence coding for residues 500-543 of *S. cerevisiae* Ume6 (SID domain) was inserted into the pET28PP (N-terminal, cleavable polyHIS fusion) vector for crystallography. The *S. cerevisiae* Ume6 fragment (residues 516-531) untagged or N-terminally tagged with a fluorescein isothiocyanate (FITC) or a 5-carboxytetramethylrhodamine (TAMRA) probe (an aminohexanoic acid (AHA) linker was inserted between FITC or TAMRA and the Ume6 fragment), were synthesized by the Analytical Core Facility of the Department of Physiology at Tufts University or by Genscript, respectively. Peptides were received as dried powders and dissolved in deuterated DMSO. A gene fragment coding for residues 402-473 of *S. cerevisiae* Sin3 (PAH2 domain) was inserted into pET28PP, pMALX(E), and pGEX6P2 (N-terminal, cleavable GST fusion) vectors for different experiments (see below). DNA sequences coding for residues 301-390 of human SIN3A or residues 155-241 of human SIN3B (PAH2 domains) were inserted into the pMALX(E) vector (received from Lars C. Pedersen^79^) for NMR experiments.

### Expression and purification of yeast and human proteins

Bicistronic yeast Sin3 PAH2-Ume6 SID: a bicistronic expression construct of yeast Sin3 (residues 402-473) and Ume6 (residues 500-543) was overexpressed in *E. coli* BL21 (DE3) in TB medium in the presence of 50 µg/mL of kanamycin. Cells were grown at 37°C to an OD of 0.8, cooled to 17°C, induced with 500 µM isopropyl-1-thio-D-galactopyranoside (IPTG), incubated overnight at 17°C, collected by centrifugation, and stored at -80°C. Cell pellets were lysed in buffer A (50 mM HEPES, pH 7.5, 300 mM NaCl, 10% glycerol, 20 mM Imidazole, and 7 mM β-mercaptoethanol) and the resulting lysate was centrifuged at 30,000 g for 40 min. Ni-NTA beads (Qiagen) were mixed with lysate supernatant for 30 min and washed with buffer A. Beads were transferred to an FPLC-compatible column and the bound protein was washed further with buffer A for 10 column volumes, and eluted with buffer B (50 mM HEPES pH 7.5, 300 mM NaCl, 10% glycerol, 300 mM Imidazole, and 7 mM BME). Human rhinovirus 3C protease was added to the eluted protein and incubated at 4°C overnight. The sample was concentrated and passed through a Superdex 200 10/300 column (GE Healthcare) in a buffer containing 20 mM HEPES, pH 7.5, 200 mM NaCl, 5% glycerol, and 1 mM TCEP. Fractions were pooled, concentrated to approximately 19 mg/mL and frozen at -80°C.

Free yeast Sin3 PAH2 and human SIN3A/B PAH2 proteins (pMALX(E)): a construct of yeast MBP-Sin3 (residues 402-473) was overexpressed in *E. coli* BL21 (DE3) in TB medium in the presence of 100 µg/mL of Ampicillin. Cells were grown at 37°C to an OD of 0.8, cooled to 17°C, induced with 500 µM IPTG, incubated overnight at 17°C, collected by centrifugation, and stored at -80°C. Cell pellets were lysed in buffer A (50 mM HEPES, pH 7.5, 500 mM NaCl, 10% glycerol, and 7 mM β-mercaptoethanol) and the resulting lysate was centrifuged at 30,000 g for 40 min. Amylose beads (NEB) were mixed with lysate supernatant for 1.5 h and washed with buffer A. Beads were transferred to an FPLC-compatible column and the bound protein was washed further with buffer A containing additional 1 M NaCl for 10 column volumes, and eluted with buffer B (25 mM HEPES pH 7.5, 200 mM NaCl, 5% glycerol, 20 mM maltose, and 7 mM BME). The eluted sample was concentrated and passed through a Superdex 200 16/600 column (GE Healthcare) in a buffer containing 20 mM HEPES, pH 7.5, 200 mM NaCl, 5% glycerol, 0.5 mM TCEP, and 1 mM DTT. Fractions were pooled, concentrated to approximately 20 mg/mL and frozen at -80°C. Other MBP-tagged proteins were expressed and purified according to the same protocol.

GST-tagged yeast Sin3 PAH2 protein: a construct of yeast GST-Sin3 (residues 402-473) was overexpressed in *E. coli* BL21 (DE3) in TB medium in the presence of 100 µg/mL of Ampicillin. Cells were grown at 37°C to an OD of 0.8, cooled to 17°C, induced with 500 µM IPTG, incubated overnight at 17°C, collected by centrifugation, and stored at -80°C. Cell pellets were lysed in buffer A (25 mM HEPES, pH 7.5, 200 mM NaCl, 5% glycerol, and 7 mM β-mercaptoethanol) and the resulting lysate was centrifuged at 30,000 g for 40 min. Glutathione beads (GE Healthcare) were mixed with lysate supernatant for 1.5 h and washed with buffer A. Beads were transferred to an FPLC-compatible column and the bound protein was washed further with buffer A containing additional 1 M NaCl for 10 column volumes, and eluted with buffer B (25 mM HEPES pH 7.5, 200 mM NaCl, 5% glycerol, 20 mM glutathione, and 7 mM BME). The eluted sample was concentrated and passed through a Superdex 200 16/600 column (GE Healthcare) in a buffer containing 20 mM HEPES, pH 7.5, 200 mM NaCl, 5% glycerol, 0.5 mM TCEP, and 1 mM DTT. All fractions were pooled.

Expression of yeast ^13^C/^15^N Sin3 PAH2: a construct of yeast GST-Sin3 PAH2 was overexpressed in *E. coli* BL21 (DE3) in 1 L of labeling medium; minimum medium supplemented with 1 g ^15^N-ammonium chloride (Cambridge isotope) and 2 g ^13^C-glucose (Cambridge isotope). Briefly, 10 mL overnight culture in LB medium were spun down gently at 500 g for 5 min to pellet the cells, resuspended in labeling medium, and transferred to 1 L labeling medium. Cells were grown at 37°C to an OD of 0.8, cooled to 17°C, induced with 500 µM IPTG, incubated overnight at 17°C, collected by centrifugation, and stored at -80°C. ^13^C/^15^N-double labeled protein was purified as described above. The GST-tag was cleaved by incubating with GST-HRV3C protease overnight in cold room. Then, cleaved GST and GST-3C were removed by passing through GSH beads, and cleaved proteins were concentrated to approximately 17 mg/mL and frozen at -80°C.

### Crystallization of the yeast Sin3 PAH2-Ume6 SID co-complex

With the aid of Formulatrix NT8, RockImager and ArtRobbins Phoenix liquid handlers, a sample of 400 µM Sin3 PAH2-Ume6 SID protein and 5 µM trypsin was mixed and co-crystallized in an equivalent volume of 1.2 M NaCitrate and 0.1 M TrisHCl pH 8.0 by sitting-drop vapor diffusion at 20°C after three days.

### Crystallization of the free yeast MBP-Sin3 PAH2 domain

A sample of 400 µM protein and 5 mM maltose was co-crystallized in (NH_4_)_2_SO_4_ and 0.1 M BisTris pH 6.0 by hanging-drop vapor diffusion at 20°C using a combination of Formulatrix NT8 and ArtRobbins Phoenix liquid handlers and visualized using a Formulatrix RockImager. Large, single crystals were transferred into crystallization buffer containing 25% glycerol prior to flash-freezing in liquid nitrogen and shipped to the synchrotron for data collection.

### Data collections and structure determinations

Bicistronic yeast Sin3 PAH2-Ume6 SID (6XAW): diffraction data from Sin3 PAH2-Ume6 SID complex crystals were collected at beamline 24ID-C of the NE-CAT at the Advanced Photon Source (Argonne National Laboratory). Data sets were integrated and scaled using XDS^80^. Structures were solved by Br-SAD using the programme Autosol in Phenix 1.13_2998 package^81^. Iterative manual model building and refinement using Phenix 1.13_2998 and Coot^82^ led to a model with excellent statistics.

Free yeast Sin3 PAH2 (6XDJ): diffraction data from MBP-Sin3 PAH2 crystals were collected at beamline 24ID-E of the NE-CAT at the Advanced Photon Source (Argonne National Laboratory). Data sets were integrated and scaled using XDS^80^. Structures were solved by molecular replacement using the programme Phaser 2.8.3^83^ and the search model of MBP from PDB entry 4JBZ. Iterative manual model building and refinement using Phenix 1.16_3549^81^ and Coot^82^ led to a model with excellent statistics.

### Fluorescence polarization (FP) assays

Dissociation constant (K_D_) of the Sin3 PAH2-Ume6 SID interaction: a solution of 30 nM FITC-Ume6 peptide in FP buffer (10.6 mM Na_2_HPO_4_, 1.93 mM NaH_2_PO_4_, 0.5 mM EDTA, 0.01% NaN_3_, pH 7.6) was prepared in Corning 3575 384-well plates to establish the K_D_ of the GST-Sin3 PAH2-FITC-Ume6 SID interaction. GST-Sin3 PAH2 was titrated to the FITC-Ume6 SID peptide starting at a concentration of 300 µM, and followed by two-fold dilutions for a total of 24 points. The same procedure was used to determine the K_D_ of the interaction with the TAMRA-Ume6 SID peptide (fixed concentration of 30 nM) but starting at a concentration of 200 µM.

IC_50_ determinations for unlabeled Ume6 SID, E6, E6R and E6S: the concentrations of FITC-Ume6 SID (30 nM) and GST-Sin3 PAH2 (3 µM) were held constant and the compounds were titrated (in triplicate) with an HP D300 (Hewlett-Packard, CA) to the assay plate. The assayed concentrations are given here in µM: 5, 6.6, 8.6, 11.2, 14.7, 19.3, 25.3, 33.1, 43.4, 56.9, 74.5, 97.7, 128.0, 167.7, 219.7, 287.9, 377.2, 494.2, 647.5 and 848.3. DMSO alone was also diluted following the same procedure and the corresponding values (defined as 100%) were used to normalize FP results at each concentration. Experiments with the TAMRA fluorescent probe were conducted in triplicate with 30 nM TAMRA-Ume6 SID and 6 µM GST-Sin3 PAH2, for the following concentrations in µM: 0, 1.4, 1.9, 2.5, 3.4, 4.7, 6.4, 8.7, 11.8, 16.0, 21.8, 29.7, 40.4, 55.0, 74.9, 101.9, 138.7, 188.8, 256.9 and 349.7. Here, the DMSO data point (i.e. 0 µM) was defined as 100% and used to normalize all FP results. Since molecules did not completely displace the tagged Ume6 SID peptide from GST-Sin3 PAH2, the lowest values were set as the asymptote during curve fitting and IC_50_ calculations, as reported previously^38^. Curve fitting was done using the Prism/Graphpad function: “inhibitor vs. response – three parameter fit”.

### Synthesis of enantiomers E6R and E6S

Synthesis of Ethyl 4-hydroxy-2-oxo-3,4-dihydro-2H-pyrido[1,2-a]pyrimidine-3-carboxylate (1): 2-aminopyridine (1038.2 mg, 11.0 mmol) and triethylmethanetricarboxylate (5000 mg, 21.5 mmol) were combined and dissolved in xylene (12 mL). The reaction was then heated to reflux for 90 min. Hexanes (60 mL) was then added and the mixture was stirred at room temperature for 2 h. The reaction was then filtered, and the precipitate was collected. The precipitate was then treated with boiling water, filtered and the resulting filtrate was collected and concentrated to afford the desired product (1067.5 mg, 41%). LC-MS (ESI) m/z 235.18 [M^+^H]^+^; calcd for C_11_H_11_N_2_O_4_^+^: 235.07.

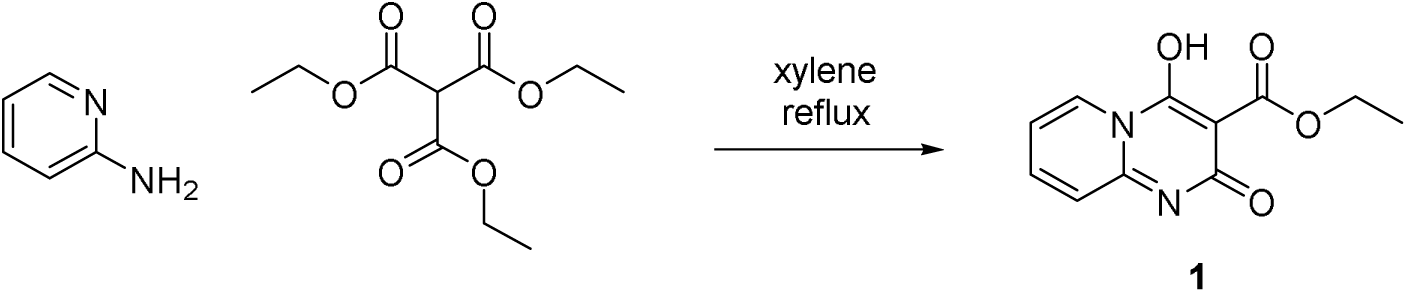

Synthesis of 4-hydroxy-2-oxo-N-((*R*)-1-phenylethyl)-3,4-dihydro-2H-pyrido[1,2-a]pyrimidine-3-carboxamide (E6R): (*R*)-(+)-1-phenylethylamine (1.7 mL, 13.5 mmol) was added to a solution of 1 (1056.9 mg, 4.5 mmol) in EtOH (8 mL). The reaction was heated to reflux for 30 h and then removed from heat and cooled to room temperature. The reaction was then filtered, and the resulting precipitate was recrystallized in EtOH to afford the desired product (966.3 mg, 69%). LC-MS (ESI) m/z 309.67 [M_+_H]_+_; calcd for C_17_H_17_N_3_O_3_^+^: 310.12. ^1^H NMR (500 MHz, DMSO-d6) δ 9.97 (d, *J* = 7.8 Hz, 1H), 9.01-8.87 (m, 1H), 8.09 (ddd, *J* = 8.7, 6.8, 1.6 Hz, 1H), 7.54 (dt, *J* = 8.9, 1.1 Hz, 1H), 7.44-7.34 (m, 5H), 7.31-7.25 (m, 1H), 5.19 (p, *J* = 7.1 Hz, 1H), 1.54 (d, *J* = 6.9 Hz, 3H).

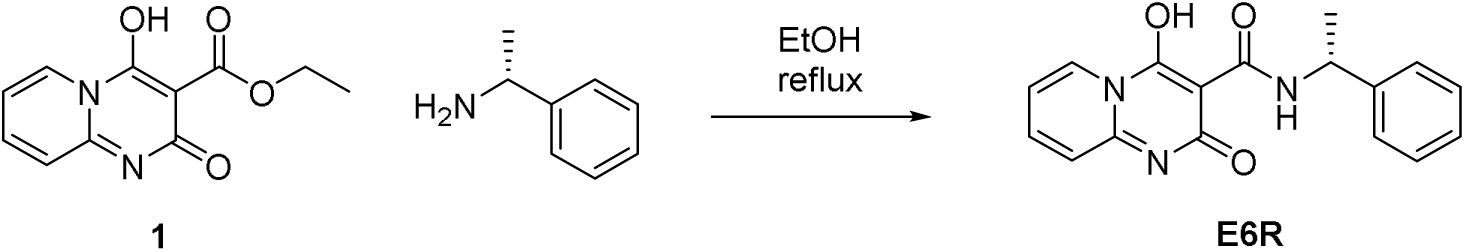

Synthesis of 4-hydroxy-2-oxo-N-((*S*)-1-phenylethyl)-3,4-dihydro-2H-pyrido[1,2-a]pyrimidine-3-carboxamide (E6S): (*S*)-(-)-1-phenylethylamine (380 µL, 3.0 mmol) was added to a solution of 1 (470.0 mg, 2 mmol) in EtOH (4 mL). The reaction was heated to reflux for 30 h and then removed from heat and cooled to room temperature. The reaction was then filtered, and the resulting precipitate was recrystallized in EtOH to afford the desired product (400.0 mg, 65%). LC-MS (ESI) m/z 309.67 [M_+_H]_+_; calcd for C_17_H_17_N_3_O_3_^+^: 310.12.

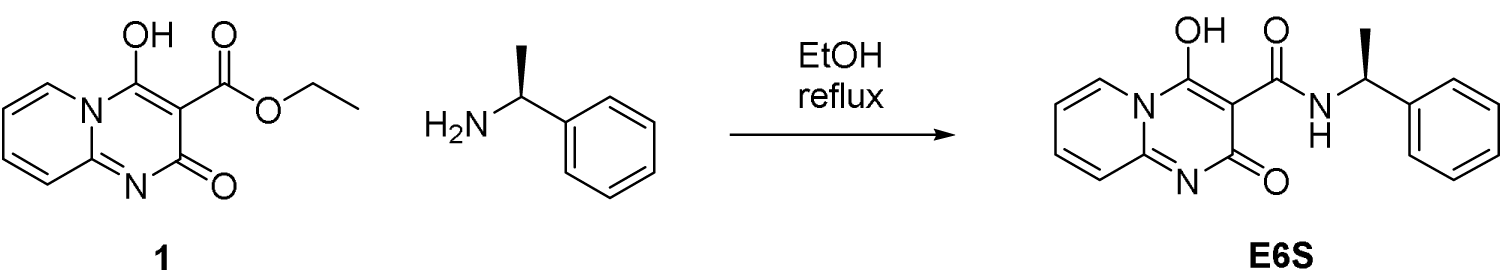

In house E6R+S racemic mixture was made by mixing equal amounts of the synthesized E6R and E6S enantiomers.

### RNA-seq in yeast and human neuroblastoma SK-N-BE(2)-C cells

Following identical treatments as for RT-qPCR measurements, RNA was extracted as described above for yeast and human neuroblastoma cells. Quality of the samples was checked by RT-qPCR of established Sin3 HDAC complex-regulated genes before doing RNA-seq. RNA-seq libraries for yeast (n = 3 per tested condition) and neuroblastoma cells (n = 3 per tested condition) were prepared according to the TruSeq stranded mRNA protocol (Illumina) and 101-bp single-end reads produced on an Illumina NovaSeq 6000 instrument. Reads were mapped to the human GRCh38 transcriptome (Ensembl cDNA release 97) or yeast *S. cerevisiae* R64-1-1 (Ensembl cDNA release 97) and quantified using Salmon v0.8.2^84^. Read counts were summed to the gene level using tximport v.1.2.0^85^ and differential expression was assessed using DESeq2 v.1.14.1^86^. Only genes whose adjusted p values < 0.05 were considered for analyses, and a |log_2_ fold-change in expression| cutoff ≥ 1 was selected. Hierarchical clustering was performed and heatmaps were generated in R using the pheatmap v.1.0.12 package (hclust complete clustering on Euclidean distance). The p-values of the overlaps were determined using two-sided Fisher’s exact test and calculated using python 2.7.10 function “scipy.stats.fisher_exact”.

### Nuclear magnetic resonance (NMR) experiments

Ligand-detected NMR experiments: all results were acquired using a 3 mm tube at 25°C on a Bruker 600 MHz Avance II system equipped with a CPPTCI cryoprobe (Bruker BioSpin Corp.). In all experiments, the concentration of E6R was fixed at 300 µM (from a stock of 25 mM in deuterated DMSO) and the concentration of the protein at 10 µM. The volume of deuterated DMSO was kept constant (1.5%) in all tested samples. Right before the experiments, the compound and protein were diluted in 1x PBS containing 0.1% deuterated dithiothreitol (DTT) to a total volume of 200 µL. Samples were diluted by 5% with D_2_O for magnetic field lock. Spectra were acquired for ligand only, protein only, and ligand+protein. Samples also contained residual protein buffers: 10.6 mM Na_2_HPO_4_, 1.93 mM N_a_H_2_PO_4_, 0.5 mM EDTA, 0.01% NaN_3_ (pH 7.6) for yeast GST-Sin3 PAH2; 20 mM HEPES-pH 7.5, 200 mM NaCl, 5% glycerol, 40 mM maltose, 0.5 mM TCEP, 1 mM DTT for human MBP-SIN3A PAH2; and 20 mM HEPES-pH 7.5, 200 mM NaCl, 5% glycerol, 0.5 mM TCEP, 1 mM DTT for human MBP-SIN3B PAH2. STD experiments^87^ were acquired with 160 scans, with protein irradiation applied for 3 s at 0 ppm and -20 ppm for the on and off resonance experiments, respectively. On and off resonance spectra were collected and stored separately. One-dimensional CPMG-R_2_ edited experiments^88^ were collected for samples with and without proteins using seven CPMG delays (0, 25, 50, 100, 300, 500, and 800 ms) with four scans per delay. Proton spectra used to measure DLB and chemical shift perturbations in the absence and presence of the proteins were recorded with 16 scans using a recycle delay of 3 s. TopSpin3.2 software (Bruker BioSpin Corp.) was used to acquire and process the data. Data analysis was performed using Matlab (MathWorks, Inc.).

Ligand titration experiments: ^15^N-SOFAST HMQC spectra were acquired for 25 µM ^15^N-labeled yeast Sin3 PAH2 protein sample in the absence and presence of 75 µM or 250 µM E6R (diluted from 25 mM stock but with total deuterated DMSO concentration of each NMR sample kept constant at 2%). The samples also contained 20 mM HEPES-pH 7.5, 200 mM NaCl, 3 mM DTT, 0.5 mM TCEP, and 10% D_2_O. Experiments were performed at 20°C by non-uniform sampling on a Bruker Avance II 600 MHz spectrometer equipped with a CPPTCI cryoprobe (Bruker BioSpin Corp.). The relaxation delay was 0.2 s and the number of scans was 64. Indirect ^15^N-dimension had a maximum of 128 complex points and 50% non-uniform sampling rate. Combined chemical shift changes in **Figure S4H** correspond to “sqrt((DeltaHcs)^2^+(DeltaNcs/5)^2^)”. A TopSpin3.2 sofware (Bruker BioSpin Corp.) was used to acquire and process data.

NMR backbone assignments: non-uniformly sampled HNCA, HNCOCA, HNCO, HNCACO, HNCACB, CBCACONH and ^15^N-NOESY experiments were performed at 20°C on a Bruker NEO 600 MHz spectrometer equipped with a cryogenic probe using the TopSpin4.1 software (Bruker BioSpin Corp.). Two 300 µM ^15^N-^13^C doubly labeled yeast Sin3 PAH2 domain samples were used to minimize the effect from sample degradation. The sample conditions were the same as those used for ligand titration experiments. The data were processed using the NMRPipe^89^ and hmsIST^90^ software, and analyzed with the CARA 1.9 programme^91^. The secondary structure of the Sin3 PAH2 domain in solution was analyzed with the talosN software^92^.

### E6R docking in yeast Sin3 PAH2 and mouse SIN3A/SIN3B PAH2 domains

AutoDock Vina with rigid receptor structures was used to dock E6R in the free PAH2 domain of *S. cerevisiae* Sin3 (structure experimentally determined in this study), restricted to the Ume6 peptide-binding region^93^. AutoDockTools was used to prepare the dockings^94,95^. Exhaustiveness was set to ten, and ten replicates for each ligand/receptor structure pair were acquired. For the mouse *M. musculus* SIN3A PAH2 model, the 30 free NMR structures/conformations available on PDB (2F05) were interrogated in the docking (ensemble docking) using QuickVina W^96^. For the mouse SIN3B PAH2 model, the 20 NMR structures/conformations available on PDB (2L9S) were interrogated using QuickVina W^96^. In both cases, blind docking against the entire mouse domain was conducted, and the best docking poses of the entire ensembles were examined. The same docking pose was obtained in multiple conformations of the NMR ensembles. Exhaustiveness was set to 100, and ten replicates for each ligand/receptor structure pair were acquired. Multiple sequence alignments of human, mouse and yeast *S. cerevisiae* SIN3 PAH2 domains were generated with Jalview Version 2^97^. Residues conserved between two, but not three, species were annotated as semi-conserved.

### Human neuroblastoma SK-N-BE(2)-C cell invasion assay

The cell invasion assay was implemented as previously reported^19^. Cells were seeded at 5×10^4^ cells per well density in a 24-well plate. Cells were treated for five days with compounds or with DMSO control (0.4%) in DMEM containing 10% FBS+PS, in quadruplicate. After incubation for 120 h, cells were changed to serum-free medium with the same compound concentrations as during the 5-day treatment, and 50,000 live cells were seeded on top of a Corning BioCoat Matrigel Invasion Chamber (cat#354480). Medium with 20% FBS was added as chemoattractant to the bottom of the well. Invasion assay was performed per manufacturer instructions. After 48 h, non-invading cells were removed by scrubbing with a cotton tip swab. Cells that remained inside the matrix were stained with 1% crystal violet dye and counted by eye. Percentage of invasive cells was calculated relative to average value of DMSO controls.

### Single cell isolation for human neuroblastoma SK-N-BE(2)-C cells

Cells were cultured as described above. They were exposed to test compounds for 16 h at 25 µM E6R, 1 µM TSA, or 1% DMSO control. Cells were harvested by incubating them in TryplE for 10 min. Cells were spun down for 5 min at 300 g, and the supernatant was removed. Cells were counted via hemocytometer to 20,000 cells and resuspended in 200 µL of media.

### Single-cell RNA-seq for human neuroblastoma SK-N-BE(2)-C cells

High-throughput scRNA-seq was performed using the Seq-Well S^3^ platform as previously described^98,99^. Briefly, 20,000 cells were applied to the surface of each Seq-Well array which had been preloaded with uniquely barcoded poly(dT) mRNA capture beads (Chemgenes). Following cell loading, arrays were washed with RPMI and then sealed with a polycarbonate membrane and placed in a 37°C incubator for 30 min. After membrane sealing, the arrays were submerged in a lysis buffer (5 M guanidine thiocyanate, 10 mM EDTA, 0.1% β-mercaptoethanol, 0.1% Sarkosyl) for 20 min, and subsequently incubated in a 2 M NaCl hybridization buffer for 40 min to promote hybridization of the mRNA to the bead-bound capture oligos. Next, the beads were removed from the arrays, and they were resuspended in a master mix for reverse transcription containing Maxima H Minus Reverse Transcriptase and buffer, dNTPs, RNase inhibitor, a template switch oligonucleotide, and PEG for 30 min at room temperature, and overnight at 52°C with end-over-end rotation. This was followed by exonuclease digestion and second strand synthesis as described previously^99^. PCR amplification was then performed to generate whole transcriptome amplification (WTA) products, and two SPRI cleanups were carried out using AMPure XP beads at 0.6x and 0.8x volumetric ratios. Then, the eluted products were quantified using a Qubit. cDNA libraries were then created using Illumina Nextera XT DNA Library Preparation Kit. Library cleanup was performed using SPRI purification as described above and library concentration and size distribution was determined using a Qubit fluorometer and Agilent TapeStation. Libraries for each sample were sequenced on a NextSeq 500 and a Nextseq 2000 with the following read structures: read 1: 21 bp; read 2: 50 bp; and index 1: 8 bp, and read 1: 26 bp; read 2: 50 bp; and index 1: 8 bp respectively. For each library 21 bases were sequenced in read 1, containing the cell barcode (12 bp) information and a unique molecular identifier (UMI, 8 bp), whereas 50 bases were obtained for the read 2 sequence. After sequencing, reads were demultiplexed using Illumina bcl2fastq (v2.20.0.422) and the resulting FASTQs were concatenated and aligned to the the human “GRCh38-2020-A’’ reference genome using StarSolo alignment pipeline (v2.7.9a) on Terra.

### Seurat QC/UMAP generation for scRNA-seq data

A total of 90,306 cells and 36,601 genes were obtained across 6 samples (three conditions: DMSO, TSA, E6R; each tested in duplicate). Subsequently, low quality cells were filtered out, including cells with fewer than 200 unique genes, greater than 9,000 unique genes, and greater than 40% mitochondrial reads, resulting in 46,751 cells and 25,468 genes. The data were then normalized, scaled by a factor of 10,000, and logarithmized. Following filtering, an initial dimensionality reduction was performed by performing PCA over the 2,000 most variable genes. We visualized cells from each array using a UMAP plot across 25 principal components over the highly variable genes and default parameters grouped by the treatment conditions. To identify differences between drug treatment conditions, we first obtained the differentially expressed genes between the conditions. Differentially expressed genes were determined using the FindMarkers function on Seurat 4.1.0 with “test.use=MAST” and default parameters. For these comparisons, we looked at the genes expressed by cells treated with either TSA or E6R with respect to DMSO. The genes were then filtered for adjusted p value <0.05, and plotted as a volcano plot using EnhancedVolcano 1.8.0. Log fold changes of the 65 genes upregulated by both TSA and E6R and 22 genes downregulated by both TSA and E6R in the bulk RNA-seq approach were determined using the FindMarkers function on Seurat 4.1.0 with “test.use=MAST”, “logfc.threshold = 0”, and “min.pct = 0”. Python 3.8.5 was used for generating the plots.

### Mice xenografted with human neuroblastoma SK-N-BE(2)-C cells

Xenograft studies were performed at the TRACE PDX platform of the Leuven Cancer Institute (LKI) at KU Leuven - UZ Leuven (https://gbiomed.kuleuven.be/english/research/50488876/54502087/Trace), according to the ARRIVE guidelines. The department has the obligatory accreditation of the authorized Belgian Ministry and is registered under license number LA1210604. The animals are housed (according to the described Belgian and European laws, guidelines and policies for animal experiments, housing and care) in the Central Animal Facilities of the university. These facilities have the obligatory accreditation of the authorized Belgian Ministry and are registered under license number LA2210393. Personnel of the Central Animal Facilities and laboratory staff have to be trained in handling animals and must have the appropriate certificate in Laboratory Animal Science. These training measures are in agreement with the Belgian law of September 13, 2004, concerning the training of people involved in animal experimentation.

Treatment experiments included 24 female nude NMRI mice (Taconic; strain BomTac:NMRI-*Foxn1^nu^*) implanted (10 weeks old at implantation) with human neuroblastoma cell line SK-N-BE(2)-C (2 million cells/implant/mouse). Sample size has been calculated using the software G power version 3.1.9.2. A two-way repetitive measure ANOVA was applied to evaluate differences between groups and between days, and the interaction between groups and days for tumor growth. By assuming an effect size of 0.25, the amount required for a power of 80% and an alpha-error probability of 0.05% is estimated to be six mice for each treatment group.

Human neuroblastoma SK-N-BE(2)-C cells were cultured as described above, with slight modifications: cells were maintained in cell culture flasks supplemented with 10% FBS and 2 mM of L-glutamine, 100 units of penicillin and 100 µg/mL of streptomycin. Cells, tested for a full panel of murine pathogens (Impact profile I, IDEXX) prior to engraftment, were resuspended in 50% matrigel solution in PBS for injection in the mouse right flank. When tumors reached a volume of approximately 200 mm^3^, mice were randomly assigned to vehicle (34% DMSO, 55% saline 0.9%, 5% ethanol, 5% kolliphor, 1% NaOH 1 M), 50 mg/kg E6R, or 20 mg/kg SAHA treatment groups (fresh powders dissolved into the vehicle solution). Treatments were administered daily intraperitoneally for 10 days or until reaching human endpoint (i.e. tumor volume > 1,500 mm^3^). Tumors were measured daily with digital calipers, and volume was estimated as V = L x W x H x (π/6) (L: length; W: width; H: height). Drug treatments did not affect mice weight. All data were analyzed using GraphPad Prism 8. Comparisons between the vehicle and treated groups were tested for statistical significance using paired t-tests.

### Mice used for the psychosis mouse model

Animal care and experimental procedures were performed according to an Institutional Animal Care and Use Committee (IACUC) approved protocol at Virginia Commonwealth University, Richmond, VA, USA.

Adult male C57BL/6N mice (10-12 weeks old at start of experiments) purchased from Taconic laboratories (Bar Arbor, ME), and C57BL/6N *Hdac2* cKO inhouse bred male mice (described previously^100^) were used for the experimental groups (vehicle and E6R). For assays including *Hdac2* cKO mice, we bred homozygous *Hdac2^loxP/loxP^* mice to the *CaMKIIα-Cre* transgenic line and used the *Hdac2^loxP/loxP^:CaMKIIα-Cre (Hdac2* cKO) mice and control littermates. All animals were housed in groups of 5-6 in 500 cm^2^ Plexiglas cages (Allentown, NJ) in the animal care facilities at 12 h light/dark cycles (lights on, 8:00 to 20:00) at 23°C with food and water *ad libitum.* Cages contained bedding (Teklad ¼” Corncob bedding, Envigo) and one Nestlests™ (Ancare, Bellmore, NY, USA). Frontal cortex tissue samples were collected as described in the method details. Mouse group sizes were based on prior experience with these procedures.

### Head-twitch response (HTR) assay in mouse

For automated recording of HTR, WT and *Hdac2* cKO male mice were ear-tagged as previously described^101^. For the *Hdac2* cKO group (n = 5), an intraperitoneal injection (2 mg/kg; 10 µL/g) of the psychedelic drug 1-(2,5-dimethoxy-4-iodophenyl)-2-aminopropane (DOI) (Sigma Aldrich) was delivered and HTR recorded and quantified. All mice were allowed to recover for 7 days before starting the chronic treatments. After this washout period of 7 days, chronic treatments over 10 days (daily intraperitoneal injections around the same time: 5 µL/g) was administered to *Hdac2* cKO and a group of WT mice. In parallel, another group of WT mice were chronically administered with vehicle (34% DMSO, 55% saline 0.9%, 5% ethanol, 5% kolliphor, 1% NaOH 1 M), or 50 mg/kg E6R (fresh powder dissolved into the vehicle solution). Twenty-four hours after the last injection, DOI (2 mg/kg; 10 µL/g) was injected and HTR subsequently recorded and quantified. For each session, the animals were habituated to the testing chamber prior to DOI administration. Mice were then returned to the chamber right after DOI administration. HTR were recorded during both habituation and post-DOI administration for 30 min and quantified in 15 min intervals. Data were processed as previously described^51,101^. Potential false positives and false negatives were visually inspected during the signal processing. Results presented in **Figures 6B and 6C**: for WT mice, one group (n = 5) was chronically treated with vehicle, and the other group (n = 6) was chronically treated with 50 mg/kg E6R. Twenty-four hours after the last injection, both groups received 2 mg/kg DOI, and HTR was recorded for each group as described above. For the *Hdac2* cKO group (n = 5), drug naïve mice were treated with 2 mg/kg DOI. Following a washout period of 7 days, and chronic treatment with 50 mg/kg E6R (over 10 days), 2 mg/kg DOI was injected again to the same mice (24 h after the last injection of E6R), and HTR recorded as described above. Therefore, for *Hdac2* cKO animals, the effect of DOI was compared before and after chronic E6R treatment within the same group.

### Drug treatment and dissection of mouse tissues

E6R was dissolved in a vehicle solution containing saline 55% v/v, DMSO 34% v/v, ethanol 5% v/v, kolliphor EL (Sigma-Aldrich) 5% v/v, and NaOH 1M 1% v/v. The vehicle group received the same solution without E6R. The number of mice per group was: WT+Vehicle (n = 5), WT+50 mg/kg E6R (n = 5), *Hdac2* cKO+Vehicle (n = 6) and *Hdac2* cKO+50 mg/kg E6R (n = 6). Animals were injected daily by intraperitoneal injection with 5 µL/g of the solution (50 mg/kg of E6R, or vehicle) at 11 AM for 10 days. Animal wellness was verified by daily visual inspection and weighing. No abnormal behavior or significant body mass decrease (>5%) was noted in any of the groups. On day 11, animals were sacrificed by bleeding via heart puncture under carbon dioxide anesthesia. The brains were quickly removed and the bilateral frontal cortex (bregma 1.90 to 1.40 mm) was dissected followed by total RNA extraction or ChIP-qPCR analyses.

### RNA extraction from mouse tissues

Qiagen™ RNeasy mini kits (Valencia, CA) were used following the manufacturer’s protocol. Extracted RNA was further purified using Qiagen™ RNeasy Mini columns to eliminate traces of DNA, and dissolved in nuclease-free water. Purity of the RNA preparation was determined as the 260/280 nm ratio with expected values between 2 and 2.3. Samples were stored at -80°C before being shipped to the GENEWIZ company (www.genewiz.com) on dry ice for library construction and RNA sequencing with Illumina HiSeq 2×150 bp configuration (single index, per lane).

### Processing RNA-seq data from mouse tissues

RNA integrity (RIN) and yield were assessed on an Agilent 2100 Bioanalyzer, and samples with RIN larger than 7 were selected. Prior to the RNA-seq, ribosomal RNA was depleted from the samples using the RiboMinus™ Human/Mouse Transcriptome Isolation Kit (Thermo Fisher Scientific, Waltham, MA). All samples were sequenced on an Illumina Hi-Seq sequencer to produce 150 bp paired-end reads. Sequencing adapters were removed using Trimmomatic v.0.33^102^. Quality control at each processing step was performed using the FastQC tool v.0.11.2 (quality base calls, GC content distribution, duplicate levels, complexity level) (https://www.bioinformatics.babraham.ac.uk/projects/fastqc/)^103^. The Mouse GRCm38/mm10 reference genome was obtained from UCSC Genome Browser Gateway (http://hgdownload.soe.ucsc.edu/goldenPath/mm10/bigZips/chromFa.tar.gz), and the corresponding gene annotation file was obtained from Ensembl (ftp://ftp.ensembl.org/pub/release-83/gtf/mus_musculus/Mus_musculus.GRCm38.83.gtf.gz) on February 20, 2016. Only autosomes, mitochondrial and sex chromosomes were used. Reads were aligned using the subread v.1.6.2 aligner^104^. Gene counts were obtained for each sample using the featureCounts v.1.2.6 software^105^. RNA-seq counts were preprocessed and analyzed for differential expression using the edgeR v.3.24.3^106^ R package. P-values for differentially expressed genes were corrected using a FDR multiple testing correction method^107^ with FDR <0.3 considered significant. In **Figure 6D**, hierarchical clustering was performed and heatmaps were generated in R using the pheatmap v.1.0.12 package (hclust complete clustering on Euclidean distance). The p values of the overlaps in **Figure 6E** were determined using the hypergeometric test. For **Figure 6F**, row-median centered log_2_(TPM+1) expression profiles for the indicated selected genes were visualized using the pheatmap package v.1.0.12. All statistical calculations were performed within R/Bioconductor environment v.3.5.3.

### ChIP-qPCR assays from frontal cortex sections of mouse brains

Tissues from frontal cortex sections of the same mice used for RNA-seq were used for ChIP-qPCR experiments. Primers used in this study are presented in **Table S7**. ChIP assays were performed using the MAGnify ChIP system (ThermoFisher, MA) according to the manufacturer’s protocol with a few modifications. Small sections (50 mg) of the frontal cortex were collected. Chromatin was sheared with the Covaris^®^ S2 system (Covaris, Woburn, MA) using the following programme: duty cycle 5%, intensity 2, 200 cycles per burst, 60 s cycle per time, 20 cycles, temperature 4°C. A volume of 10 µL chromatin was used per sample. A total of 1 µg of the anti-SIN3A (Abcam, Cambridge, MA, USA, cat#ab3479)^108^ or anti-IgG (Abcam, Cambridge, MA, USA, cat#ab37355) antibody was used per sample. Quantitative real-time PCR analysis was performed following magnetic beads-based DNA purification. The DNA samples were amplified in quadruplicate reactions as previously described^100^ using the QuantStudio 6 Flex Real-Time PCR system (Applied Biosystems) using the PowerUP SYBR Green Master Mix (Applied Biosystems).

### Data and code availability

All raw data are provided with this paper (**Source Data**) and are available upon request. The mass spectrometry proteomics data have been deposited to the ProteomeXchange Consortium via the PRIDE partner repository with the dataset identifiers PXD031200 and 10.6019/PXD031200 (username; password: gPbEXy77). Co-complex Sin3 PAH2-Ume6 SID and free Sin3 PAH2 structural data have been deposited in the Protein Data Bank (PDB) under the accession codes 6XAW and 6XDJ, respectively. NMR data have been deposited in the Biological Magnetic Resonance Data Bank (BMRB) with the identifier 51585 (https://bmrb.io/author_view/51585_hy_aywfbqba.str). ChIP-seq data have been deposited in the Gene Expression Omnibus (GEO) database under the accession code GSE211772 (https://www.ncbi.nlm.nih.gov/geo/query/acc.cgi?acc=GSE211772; token: wjqryoogtjszfqd). RNA-seq data for yeast, human neuroblastoma cells, and mouse FC tissue samples have been deposited in the GEO database under the accession codes GSE211775 (https://www.ncbi.nlm.nih.gov/geo/query/acc.cgi?acc=GSE211775; token: srsrqkuaxnmhxeh), GSE211773 (https://www.ncbi.nlm.nih.gov/geo/query/acc.cgi?acc=GSE211773; token: mrcpcimmznwlnyr), and GSE211774 (https://www.ncbi.nlm.nih.gov/geo/query/acc.cgi?acc=GSE211774; token: knevcqamddwlfoh), respectively. Single-cell RNA-seq data have been deposited in the GEO database under the accession code GSE211593 (https://www.ncbi.nlm.nih.gov/geo/query/acc.cgi?acc=GSE211593). This paper does not report original codes, and software used in the study is publicly available and referenced in the text.

### Statistical analyses

Except otherwise indicated, statistical analyses were performed on Prism (GraphPad) with each statistical test, definitions of center (mean or median) and dispersion (standard error of the mean, standard deviation, box and whiskers), and the number of samples per group (n, referring to number of biologically independent replicates) indicated in the corresponding figure legends and/or method details. Except otherwise indicated, calculations and normalizations of data, and definitions of standard error of the proportion were performed in Microsoft Excel 2011. Statistical significance was determined by p ≤ 0.05. Statistically significant comparisons in each figure are indicated with asterisks, *p ≤ 0.05; **p ≤ 0.01; ***p ≤ 0.001; ****p ≤ 0.0001.

## ACKNOWLEDGMENTS

We thank all members of the Vidal, Twizere, Dequiedt, González-Maeso, Dhe-Paganon, Buhrlage, Arthanari, and Shalek laboratories for helpful discussions during this project, as well as Kimberly Stegmaier and Clare Malone (Dana-Farber Cancer Institute and Broad Institute of Harvard and M.I.T.) for providing the neuroblastoma cell line, and Alessio Ciulli (University of Dundee) for providing critical feedback. We also thank Sam Kunes (Harvard University), Cédric Diot and Marian Walhout (University of Massachusetts Medical School) for helpful discussions throughout this work. Xavier Morelli, Yves Collette, and Philippe Roche from the Centre de Recherche en Cancérologie de Marseille (Aix-Marseille University) are thanked for providing the 2P2I chemical library, as well as John Beutler from the National Cancer Institute, U.S.A., for providing the libraries of natural products. David Stillman (University of Utah) and Kevin Struhl (Harvard Medical School) are thanked for providing, among other resources, the YEplac181-Sin3-(HA)_3_ plasmid. The authors also thank Jennifer Smith, Caroline Shamu, David Wrobel and the entire staff at the Harvard Medical School’s Institute of Chemistry and Cell Biology-Longwood Screening Facility for their assistance in the chemical screening, as well as Lauren Brown and John Porco Jr. from the Boston University Center for Molecular Discovery. Gregory Heffron and Charles Sheahan (Harvard Medical School’s NMR Core Facility), as well as Hideki Endoh, Monica Roberson and Benjamin Silva are thanked for the technical assistance. The authors also thank Adeline Deward for designing the graphical abstract, Fabian Stinkens and Felicia Vervloesem from the TRACE PDX platform of KU Leuven, as well as staff members from the University of Liège-GIGA Genomics platform and the Dana-Farber Cancer Institute Molecular Biology Core facility. We thank Lars Pedersen (NIEHS, National Institutes of Health (N.I.H.), U.S.A.) for providing the pMALX(E) plasmid. This work was based upon research conducted at the Northeastern Collaborative Access Team beamlines, which were funded by the National Institute of General Medical Sciences from the U.S.A. N.I.H. (P30 GM124165). The Eiger 16M detector on the 24-ID-E beam line was funded by a U.S.A. N.I.H.-O.R.I.P. H.E.I. grant (S10OD021527). This research used resources of the Advanced Photon Source, a U.S.A. Department of Energy (D.O.E.) Office of Science User Facility operated for the D.O.E. Office of Science by Argonne National Laboratory under Contract No. DE-AC02-06CH11357. This work was primarily supported by the U.S.A. N.I.H. grants R01GM130885 (M.V.), R01GM133185 (M.V., M.A.C.), R01CA266194 (M.V.), R01MH084894 (J.G.M.), R01MH111940 (J.G.M.), P30DA033934 (J.G.M.), T32MH020030 (M.d.l.F.R.), with additional support from the Bridge Project, a partnership between the Koch Institute for Integrative Cancer Research at Massachusetts Institute of Technology and the Dana-Farber/Harvard Cancer Center (A.K.S., M.V.), Belgian Fonds de la Recherche Scientifique F.R.S.-FNRS Télévie grant FC27371 (Crédit no 7454518F) (J.O., J.C.T.), Wallonia-Brussels International (WBI)-World Excellence doctoral fellowships (J.O., F.L.), a University of Liège mobility grant (J.O.), the Léon Fredericq Foundation (J.O., F.D., J.C.T.), a Claudia Adams Barr award (S.G.C.), a Belgian Fonds de la Recherche Scientifique F.R.S.-FNRS Télévie grant FC31747 (Crédit no 7459421F) (F.L., J.C.T.), a Belgian American Educational Foundation doctoral research fellowship (F.L.), a Belgian Fonds de la Recherche Scientifique F.R.S.-FNRS Télévie grant FC38907 (Crédit no 7462320F) (J.B., J.C.T.), the Belgian Federation for Cancer grant 2018-035 to the KU Leuven TRACE platform (M.F.B., E.L.), the Consorzio per lo Sviluppo dei Sistemi a Grande Interfase (A.H.), an Austrian Science Fund’s Schrödinger fellowship J3872-B21 (A.B.), an American Heart Association’s fellowship 19POST34380800 (A.B.), the Linde Family Foundation sponsored research agreement grants NIBR, 3DC, DDCF (S.D.P.), and the University of Liège Special Funds (F.D., J.C.T.). F.D. and M.V. are Chercheurs Qualifiés Honoraires, J.C.T. is a Maître de Recherche, and T.M.O. is a Chargée de Recherche of the Belgian Fonds de la Recherche Scientifique F.R.S.-FNRS.

## AUTHOR CONTRIBUTIONS

The project was conceived and supervised by J.O., S.G.C., J.C.T., and M.V. J.O. and M.V. constructed the yeast strains. J.O. developed the *URS-URA3* reporter assay, conducted high-throughput chemical screens, retested the selected compounds, and performed RT-qPCR experiments in yeast cells. J.O. and S.G.C. conducted RT-qPCR experiments with neuroblastoma cells, and ChIP-qPCR experiments in yeast. J.O. and A.B. performed FP experiments. S.G.C. performed HDAC activity assays, binary PPI mapping and original inhibition tests by GPCA with help from J.O. S.S. designed, performed, and analyzed experiments for the psychosis mouse model with significant contributions from M.d.l.F.R., and feedback from J.O., S.G.C., and J.G.M. T.M.O. analyzed RNA-seq and ChIP-seq data with help from J.O., S.G.C. and T.H., and feedback from Y.W., J.C.T., F.D. and M.V. F.L. provided significant technical assistance with RT-qPCR and ChIP-qPCR experiments in yeast cells. V.V.B.Jr conducted the cell invasion assay with help from J.O. C.G. performed the *in silico* dockings with feedback from J.O. and A.B. P.W.C. helped analyzing and visualizing NMR data. J.B. conducted the independent retest of compound E6 by GPCA with feedback from J.O. and J.C.T. E.A.G., J.L., K.S., and Z.C.Y. purified Ume6 and SIN3 proteins, and conducted crystal screens under the supervision of H.S.S. and S.D.P., and feedback from J.O., A.B. and Z.Y.J.S. B.H. and A.C.V. synthesized E6 enantiomers with feedback from S.J.B. J. Bruyr purified HDAC1 with help from J.C.T. S.I., T.J., J.D.B., and S.K.N. performed scRNA-seq experiments and analyzed data with help from A.K.S. and feedback from J.O., D.E.H. and F.D. A.R. constructed plasmids expressing Ume6 and Sin3 with help from F.L. and K. Spirohn. H.Y. constructed the HoY013 yeast strain with feedback from J.O. and A.R. N.C., A.S., and A.H. helped conducting the HDAC activity assays. D.V. conducted mass-spectrometry experiments with help from J.C.T., and analyzed data with feedback from J.O., T.H., F.D., and J.C.T. M.F.B. and E.L. supervised the mouse xenograft experiments, with feedback from J.O. and J.C.T. I.L. supervised orthogonal KISS experiments with feedback from J.O., S.G.C. and J.T. H.S.S. and S.D.P. determined crystal structures. M.G.D. analyzed RNA-seq data for the mouse psychosis model with feedback from S.S., M.d.l.F.R., and J.G.M. L.W., K.D., M.A.C., and D.E.H. provided critical insights. Z.Y.J.S. performed NMR experiments with help from J.O. and A.B., and feedback from P.W.C., and H.A. A.B. helped analyzing and visualizing structures with critical feedback from J.O. and K.D. S.J.B. prioritized compounds from the primary screen with help from J.O. J.O. wrote the manuscript with help from T.M.O., D.E.H., J.C.T., and F.D., and critical feedback from M.V. All authors discussed and reviewed the results.

## REFERENCES

1 Holdgate, G. A., Meek, T. D. & Grimley, R. L. Mechanistic enzymology in drug discovery: a fresh perspective. Nat Rev Drug Discov 17, 115–132 https://doi.org:10.1038/nrd.2017.219 (2018).

2 Luck, K. et al. A reference map of the human binary protein interactome. Nature 580, 402–408 https://doi.org:10.1038/s41586-020-2188-x (2020).

3 Drew, K., Wallingford, J. B. & Marcotte, E. M. hu.MAP 2.0: integration of over 15,000 proteomic experiments builds a global compendium of human multiprotein assemblies. Mol Syst Biol 17, e10016 https://doi.org:10.15252/msb.202010016 (2021).

4 Li, Y. et al. Zinc-dependent Deacetylase (HDAC) Inhibitors with Different Zinc Binding Groups. Curr Top Med Chem 19, 223–241 https://doi.org:10.2174/1568026619666190122144949 (2019).

5 Eckschlager, T., Plch, J., Stiborova, M. & Hrabeta, J. Histone Deacetylase Inhibitors as Anticancer Drugs. Int J Mol Sci 18, 1414 https://doi.org:10.3390/ijms18071414 (2017).

6 Ho, T. C. S., Chan, A. H. Y. & Ganesan, A. Thirty Years of HDAC Inhibitors: 2020 Insight and Hindsight. J Med Chem 63, 12460–12484 https://doi.org:10.1021/acs.jmedchem.0c00830 (2020).

7 Bondarev, A. D. et al. Recent developments of HDAC inhibitors: Emerging indications and novel molecules. Br J Clin Pharmacol 87, 4577–4597 https://doi.org:10.1111/bcp.14889 (2021).

8 Marks, P. A. & Breslow, R. Dimethyl sulfoxide to vorinostat: development of this histone deacetylase inhibitor as an anticancer drug. Nat Biotechnol 25, 84–90 https://doi.org:10.1038/nbt1272 (2007).

9 Glaser, K. B. et al. Gene Expression Profiling of Multiple Histone Deacetylase (HDAC) Inhibitors: Defining a Common Gene Set Produced by HDAC Inhibition in T24 and MDA Carcinoma Cell Lines. Mol Cancer Ther 2, 151–163 (2003).

10 Li, W. & Sun, Z. Mechanism of Action for HDAC Inhibitors-Insights from Omics Approaches. Int J Mol Sci 20, 1616 https://doi.org:10.3390/ijms20071616 (2019).

11 Shah, R. R. Safety and Tolerability of Histone Deacetylase (HDAC) Inhibitors in Oncology. Drug Saf 42, 235–245 https://doi.org:10.1007/s40264-018-0773-9 (2019).

12 Peng, X., Sun, Z., Kuang, P. & Chen, J. Recent progress on HDAC inhibitors with dual targeting capabilities for cancer treatment. Eur J Med Chem 208, 112831 https://doi.org:10.1016/j.ejmech.2020.112831 (2020).

13 Maolanon, A. R., Madsen, A. S. & Olsen, C. A. Innovative Strategies for Selective Inhibition of Histone Deacetylases. Cell Chem Biol 23, 759–768 https://doi.org:10.1016/j.chembiol.2016.06.011 (2016).

14 Millard, C. J., Watson, P. J., Fairall, L. & Schwabe, J. W. R. Targeting Class I Histone Deacetylases in a “Complex” Environment. Trends Pharmacol Sci 38, 363–377 https://doi.org:10.1016/j.tips.2016.12.006 (2017).

15 Bantscheff, M. et al. Chemoproteomics profiling of HDAC inhibitors reveals selective targeting of HDAC complexes. Nat Biotechnol 29, 255–265 https://doi.org:10.1038/nbt.1759 (2011).

16 Joshi, P. et al. The functional interactome landscape of the human histone deacetylase family. Mol Syst Biol 9, 672 https://doi.org:10.1038/msb.2013.26 (2013).

17 Thaler, F. & Mercurio, C. Towards Selective Inhibition of Histone Deacetylase Isoforms: What Has Been Achieved, Where We Are and What Will Be Next. ChemMedChem 9, 523–536 https://doi.org:10.1002/cmdc.201300413 (2014).

18 Adams, M. K. et al. Differential Complex Formation via Paralogs in the Human Sin3 Protein Interaction Network. Mol Cell Proteomics 19, 1468–1484 https://doi.org:10.1074/mcp.RA120.002078 (2020).

19 Adams, G. E., Chandru, A. & Cowley, S. M. Co-repressor, co-activator and general transcription factor: the many faces of the Sin3 histone deacetylase (HDAC) complex. Biochem J 475, 3921–3932 https://doi.org:10.1042/bcj20170314 (2018).

20 Le Guezennec, X., Vermeulen, M. & Stunnenberg, H. G. Molecular characterization of Sin3 PAH-domain interactor specificity and identification of PAH partners. Nucleic Acids Res 34, 3929–3937 https://doi.org:10.1093/nar/gkl537 (2006).

21 Farias, E. F. et al. Interference with Sin3 function induces epigenetic reprogramming and differentiation in breast cancer cells. Proc Natl Acad Sci USA 107, 11811–11816 https://doi.org:10.1073/pnas.1006737107 (2010).

22 Conforti, P. et al. Binding of the repressor complex REST-mSIN3b by small molecules restores neuronal gene transcription in Huntington’s disease models. J Neurochem 127, 22–35 https://doi.org:10.1111/jnc.12348 (2013).

23 Kwon, Y.-J. et al. Selective Inhibition of SIN3 Corepressor with Avermectins as a Novel Therapeutic Strategy in Triple-Negative Breast Cancer. Mol Cancer Ther 14, 1824–1836 https://doi.org:10.1158/1535-7163.Mct-14-0980-t (2015).

24 Kadamb, R. et al. Invasive phenotype in triple negative breast cancer is inhibited by blocking SIN3A–PF1 interaction through KLF9 mediated repression of ITGA6 and ITGB1. Transl Oncol 16, 101320 https://doi.org:10.1016/j.tranon.2021.101320 (2022).

25 Ueda, H. et al. A mimetic of the mSin3-binding helix of NRSF/REST ameliorates abnormal pain behavior in chronic pain models. Bioorg Med Chem Lett 27, 4705–4709 https://doi.org:10.1016/j.bmcl.2017.09.006 (2017).

26 Kurita, J. I., Hirao, Y., Nakano, H., Fukunishi, Y. & Nishimura, Y. Sertraline, chlorprothixene, and chlorpromazine characteristically interact with the REST-binding site of the corepressor mSin3, showing medulloblastoma cell growth inhibitory activities. Sci Rep 8, 13763 https://doi.org:10.1038/s41598-018-31852-1 (2018).

27 Taunton, J., Hassig, C. A. & Schreiber, S. L. A mammalian histone deacetylase related to the yeast transcriptional regulator Rpd3p. Science 272, 408–411 https://doi.org:10.1126/science.272.5260.408 (1996).

28 Vidal, M., Buckley, A. M., Hilger, F. & Gaber, R. F. Direct selection for mutants with increased K+ transport in Saccharomyces cerevisiae. Genetics 125, 313–320 https://doi.org:10.1093/genetics/125.2.313 (1990).

29 Vidal, M. & Gaber, R. F. RPD3 encodes a second factor required to achieve maximum positive and negative transcriptional states in Saccharomyces cerevisiae. Mol Cell Biol 11, 6317–6327 https://doi.org:doi:10.1128/mcb.11.12.6317-6327.1991 (1991).

30 Vidal, M., Strich, R., Esposito, R. E. & Gaber, R. F. RPD1 (SIN3/UME4) is required for maximal activation and repression of diverse yeast genes. Mol Cell Biol 11, 6306–6316 https://doi.org:10.1128/mcb.11.12.6306-6316.1991 (1991).

31 Vidal, M. Playing Hide-and-Seek with Yeast. Cell 166, 1069–1073 https://doi.org:10.1016/j.cell.2016.08.018 (2016).

32 Sardiu, M. E. et al. Determining protein complex connectivity using a probabilistic deletion network derived from quantitative proteomics. PLoS One 4, e7310 https://doi.org:10.1371/journal.pone.0007310 (2009).

33 Vidal, M., Brachmann, R. K., Fattaey, A., Harlow, E. & Boeke, J. D. Reverse two-hybrid and one-hybrid systems to detect dissociation of protein-protein and DNA-protein interactions. Proc Natl Acad Sci USA 93, 10315–10320 https://doi.org:10.1073/pnas.93.19.10315 (1996).

34 Strich, R. et al. UME6 is a key regulator of nitrogen repression and meiotic development. Genes Dev 8, 796–810 https://doi.org:10.1101/gad.8.7.796 (1994).

35 Yoshida, M., Horinouchi, S. & Beppu, T. Trichostatin A and trapoxin: novel chemical probes for the role of histone acetylation in chromatin structure and function. Bioessays 17, 423–430 https://doi.org:10.1002/bies.950170510 (1995).

36 Moye-Rowley, W. S. Transcriptional control of multidrug resistance in the yeast Saccharomyces. Prog Nucleic Acid Res Mol Biol 73, 251–279 https://doi.org:10.1016/s0079-6603(03)01008-0 (2003).

37 Frumm, Stacey M. et al. Selective HDAC1/HDAC2 Inhibitors Induce Neuroblastoma Differentiation. Chem Biol 20, 713–725 https://doi.org:10.1016/j.chembiol.2013.03.020 (2013).

38 Phimmachanh, M., Han, J. Z. R., O’Donnell, Y. E. I., Latham, S. L. & Croucher, D. R. Histone Deacetylases and Histone Deacetylase Inhibitors in Neuroblastoma. Front Cell Dev Biol 8, 578770 https://doi.org:10.3389/fcell.2020.578770 (2020).

39 Choi, S. G. et al. Maximizing binary interactome mapping with a minimal number of assays. Nat Commun 10, 3907 https://doi.org:10.1038/s41467-019-11809-2 (2019).

40 Kadosh, D. & Struhl, K. Repression by Ume6 Involves Recruitment of a Complex Containing Sin3 Corepressor and Rpd3 Histone Deacetylase to Target Promoters. Cell 89, 365–371 https://doi.org:10.1016/S0092-8674(00)80217-2 (1997).

41 Sardiu, M. E. et al. Suberoylanilide hydroxamic acid (SAHA)-induced dynamics of a human histone deacetylase protein interaction network. Mol Cell Proteomics 13, 3114–3125 https://doi.org:10.1074/mcp.M113.037127 (2014).

42 Daily, K., Patel, V. R., Rigor, P., Xie, X. & Baldi, P. MotifMap: integrative genome-wide maps of regulatory motif sites for model species. BMC Bioinform 12, 495 https://doi.org:10.1186/1471-2105-12-495 (2011).

43 Rossi, M. J., Lai, W. K. M. & Pugh, B. F. Simplified ChIP-exo assays. Nat Commun 9, 2842 https://doi.org:10.1038/s41467-018-05265-7 (2018).

44 Washburn, B. K. & Esposito, R. E. Identification of the Sin3-binding site in Ume6 defines a two-step process for conversion of Ume6 from a transcriptional repressor to an activator in yeast. Mol Cell Biol 21, 2057–2069 https://doi.org:10.1128/MCB.21.6.2057-2069.2001 (2001).

45 Kumar, G. S., Xie, T., Zhang, Y. & Radhakrishnan, I. Solution structure of the mSin3A PAH2-Pf1 SID1 complex: a Mad1/Mxd1-like interaction disrupted by MRG15 in the Rpd3S/Sin3S complex. J Mol Biol 408, 987–1000 https://doi.org:10.1016/i.imb.2011.03.043 (2011).

46 van Ingen, H., Baltussen, M. A. H., Aelen, J. & Vuister, G. W. Role of Structural and Dynamical Plasticity in Sin3: The Free PAH2 Domain is a Folded Module in mSin3B. J Mol Biol 358, 485–497 https://doi.org:10.1016/i.imb.2006.01.100 (2006).

47 Nishikawa, J. L. et al. Inhibiting fungal multidrug resistance by disrupting an activator-Mediator interaction. Nature 530, 485–489 https://doi.org:10.1038/nature16963 (2016).

48 Rettig, I. et al. Selective inhibition of HDAC8 decreases neuroblastoma growth in vitro and in vivo and enhances retinoic acid-mediated differentiation. Cell Death Dis 6, e1657 https://doi.org:10.1038/cddis.2015.24 (2015).

49 Kurita, M. et al. HDAC2 regulates atypical antipsychotic responses through the modulation of mGlu2 promoter activity. Nat Neurosci 15, 1245–1254 https://doi.org:10.1038/nn.3181 (2012).

50 de la Fuente Revenga, M., et al. HDAC2-dependent Antipsychotic-like Effects of Chronic Treatment with the HDAC Inhibitor SAHA in Mice. Neuroscience 388, 102–117 https://doi.org:10.1016/i.neuroscience.2018.07.010 (2018).

51 de la Fuente Revenga, M., et al. Fully automated head-twitch detection system for the study of 5-HT2A receptor pharmacology in vivo. Sci Rep 9, 14247 https://doi.org:10.1038/s41598-019-49913-4 (2019).

52 Bridi, M. et al. Transcriptional corepressor SIN3A regulates hippocampal synaptic plasticity via Homer1/mGluR5 signaling. JCI Insight 5, e92385 https://doi.org:10.1172/jci.insight.92385 (2020).

53 Matosin, N. et al. Molecular evidence of synaptic pathology in the CA1 region in schizophrenia. NPJ Schizophr 2, 16022 https://doi.org:10.1038/npjschz.2016.22 (2016).

54 Elfving, B. et al. Differential expression of synaptic markers regulated during neurodevelopment in a rat model of schizophrenia-like behavior. Prog Neuropsychopharmacol Biol Psychiatry 95, 109669 https://doi.org:10.1016/j.pnpbp.2019.109669 (2019).

55 Huttlin, E. L. et al. Dual proteome-scale networks reveal cell-specific remodeling of the human interactome. Cell 184, 3022–3040.e3028 https://doi.org:10.1016/i.cell.2021.04.011 (2021).

56 Gorgulla, C. et al. An open-source drug discovery platform enables ultra-large virtual screens. Nature 580, 663–668 https://doi.org:10.1038/s41586-020-2117-z (2020).

57 Humphreys, I. R. et al. Computed structures of core eukaryotic protein complexes. Science 374, eabm4805 https://doi.org:doi:10.1126/science.abm4805 (2021).

58 Cao, L. et al. Design of protein-binding proteins from the target structure alone. Nature 605, 551–560 https://doi.org:10.1038/s41586-022-04654-9 (2022).

59 Callaway, E. Revolutionary cryo-EM is taking over structural biology. Nature 578, 201 https://doi.org:10.1038/d41586-020-00341-9 (2020).

60 Yu, H. et al. High-quality binary protein interaction map of the yeast interactome network. Science 322, 104–110 https://doi.org:10.1126/science.1158684 (2008).

61 Baganz, F., Hayes, A., Marren, D., Gardner, D. C. & Oliver, S. G. Suitability of replacement markers for functional analysis studies in Saccharomyces cerevisiae. Yeast 13, 1563–1573 https://doi.org:10.1002/(SICI)1097-0061(199712)13:16<1563 (1997).

62 Wendland, J. PCR-based methods facilitate targeted gene manipulations and cloning procedures. Curr Genet 44, 115–123 https://doi.org:10.1007/s00294-003-0436-x (2003).

63 Gardner, J. M. & Jaspersen, S. L. Manipulating the yeast genome: deletion, mutation, and tagging by PCR. Methods Mol Biol 1205, 45–78 https://doi.org:10.1007/978-1-4939-1363-3_5 (2014).

64 Guldener, U., Heck, S., Fielder, T., Beinhauer, J. & Hegemann, J. H. A new efficient gene disruption cassette for repeated use in budding yeast. Nucleic Acids Res 24, 2519–2524 https://doi.org:10.1093/nar/24.13.2519 (1996).

65 Choi, S. G., Jia, J., Pfeffer, R. L. & Sealfon, S. C. G proteins and autocrine signaling differentially regulate gonadotropin subunit expression in pituitary gonadotrope. J Biol Chem 287, 21550–21560 https://doi.org:10.1074/jbc.M112.348607 (2012).

66 Teste, M. A., Duquenne, M., Francois, J. M. & Parrou, J. L. Validation of reference genes for quantitative expression analysis by real-time RT-PCR in Saccharomyces cerevisiae. BMC Mol Biol 10, 99 https://doi.org:10.1186/1471-2199-10-99 (2009).

67 Zhan, C. et al. Identification of reference genes for qRT-PCR in human lung squamous-cell carcinoma by RNA-Seq. Acta Biochim Biophys Sin (Shanghai) 46, 330–337 https://doi.org:10.1093/abbs/gmt153 (2014).

68 Baell, J. B. & Holloway, G. A. New substructure filters for removal of pan assay interference compounds (PAINS) from screening libraries and for their exclusion in bioassays. J Med Chem 53, 2719–2740 https://doi.org:10.1021/jm901137j (2010).

69 Hsu, C. W. et al. Identification of HDAC Inhibitors Using a Cell-Based HDAC I/II Assay. J Biomol Screen 21, 643–652 https://doi.org:10.1177/1087057116629381 (2016).

70 Fischle, W. et al. Enzymatic activity associated with class II HDACs is dependent on a multiprotein complex containing HDAC3 and SMRT/N-CoR. Mol Cell 9, 45–57 https://doi.org:10.1016/s1097-2765(01)00429-4 (2002).

71 Calderone, A., Castagnoli, L. & Cesareni, G. mentha: a resource for browsing integrated protein-interaction networks. Nat Methods 10, 690–691 https://doi.org:10.1038/nmeth.2561 (2013).

72 Lievens, S. et al. Kinase Substrate Sensor (KISS), a mammalian in situ protein interaction sensor. Mol Cell Proteomics 13, 3332–3342 https://doi.org:10.1074/mcp.M114.041087 (2014).

73 Barski, A. et al. High-resolution profiling of histone methylations in the human genome. Cell 129, 823–837 https://doi.org:10.1016/j.cell.2007.05.009 (2007).

74 Jezek, M., Jacques, A., Jaiswal, D. & Green, E. M. Chromatin Immunoprecipitation (ChIP) of Histone Modifications from Saccharomyces cerevisiae. J Vis Exp https://doi.org:doi.org:10.3791/57080 (2017).

75 Sertil, O., Vemula, A., Salmon, S. L., Morse, R. H. & Lowry, C. V. Direct role for the Rpd3 complex in transcriptional induction of the anaerobic DAN/TIR genes in yeast. Mol Cell Biol 27, 2037–2047 https://doi.org:10.1128/MCB.02297-06 (2007).

76 Pal, S., Postnikoff, S. D., Chavez, M. & Tyler, J. K. Impaired cohesion and homologous recombination during replicative aging in budding yeast. Sci Adv 4, eaaq0236 https://doi.org:10.1126/sciadv.aaq0236 (2018).

77 Robinson, J. T. et al. Integrative genomics viewer. Nat Biotechnol 29, 24–26 https://doi.org:10.1038/nbt.1754 (2011).

78 Zhang, Y. et al. Model-based analysis of ChIP-Seq (MACS). Genome Biol 9, R137 https://doi.org:10.1186/gb-2008-9-9-r137 (2008).

79 Moon, A. F., Mueller, G. A., Zhong, X. & Pedersen, L. C. A synergistic approach to protein crystallization: combination of a fixed-arm carrier with surface entropy reduction. Protein Sci 19, 901–913 https://doi.org:10.1002/pro.368 (2010).

80 Kabsch, W. Integration, scaling, space-group assignment and post-refinement. Acta Crystallogr D Biol Crystallogr 66, 133–144 https://doi.org:10.1107/S0907444909047374 (2010).

81 Adams, P. D. et al. PHENIX: a comprehensive Python-based system for macromolecular structure solution. Acta Crystallogr D Biol Crystallogr 66, 213–221 https://doi.org:10.1107/S0907444909052925 (2010).

82 Emsley, P. & Cowtan, K. Coot: model-building tools for molecular graphics. Acta Crystallogr D Biol Crystallogr 60, 2126–2132 https://doi.org:10.1107/S0907444904019158 (2004).

83 McCoy, A. J. et al. Phaser crystallographic software. J Appl Crystallogr 40, 658–674 https://doi.org:10.1107/S0021889807021206 (2007).

84 Patro, R., Duggal, G., Love, M. I., Irizarry, R. A. & Kingsford, C. Salmon provides fast and bias-aware quantification of transcript expression. Nat Methods 14, 417–419 https://doi.org:10.1038/nmeth.4197 (2017).

85 Soneson, C., Love, M. I. & Robinson, M. D. Differential analyses for RNA-seq: transcript-level estimates improve gene-level inferences. F1000Res 4, 1521 https://doi.org:10.12688/f1000research.7563.2 (2015).

86 Love, M. I., Huber, W. & Anders, S. Moderated estimation of fold change and dispersion for RNA-seq data with DESeq2. Genome Biol 15, 550 https://doi.org:10.1186/s13059-014-0550-8 (2014).

87 Mayer, M. & Meyer, B. Group epitope mapping by saturation transfer difference NMR to identify segments of a ligand in direct contact with a protein receptor. J Am Chem Soc 123, 6108–6117 https://doi.org:10.1021/ja0100120 (2001).

88 Hajduk, P. J., Olejniczak, E. T. & Fesik, S. W. One-Dimensional Relaxation- and Diffusion-Edited NMR Methods for Screening Compounds That Bind to Macromolecules. J Am Chem Soc 119, 12257–12261 https://doi.org:10.1021/ja9715962 (1997).

89 Delaglio, F. et al. NMRPipe: a multidimensional spectral processing system based on UNIX pipes. J Biomol NMR 6, 277–293 https://doi.org:10.1007/BF00197809 (1995).

90 Hyberts, S. G., Milbradt, A. G., Wagner, A. B., Arthanari, H. & Wagner, G. Application of iterative soft thresholding for fast reconstruction of NMR data non-uniformly sampled with multidimensional Poisson Gap scheduling. J Biomol NMR52, 315–327 https://doi.org:10.1007/s10858-012-9611-z (2012).

91 Keller, R. L. J. Optimizing the process of nuclear magnetic resonance spectrum analysis and computer aided resonance assignment, ETH Zurich, https://doi.org:10.3929/ethz-a-005068942 (2005).

92 Shen, Y. & Bax, A. Protein backbone and sidechain torsion angles predicted from NMR chemical shifts using artificial neural networks. J Biomol NMR 56, 227–241 https://doi.org:10.1007/s10858-013-9741-y (2013).

93 Trott, O. & Olson, A. J. AutoDock Vina: improving the speed and accuracy of docking with a new scoring function, efficient optimization, and multithreading. J Comput Chem 31, 455–461 https://doi.org:10.1002/jcc.21334 (2010).

94 Morris, G. M. et al. AutoDock4 and AutoDockTools4: Automated docking with selective receptor flexibility. J Comput Chem 30, 2785–2791 https://doi.org:10.1002/jcc.21256 (2009).

95 Sanner, M. F. Python: a programming language for software integration and development. J Mol Graph Model 17, 57–61 (1999).

96 Hassan, N. M., Alhossary, A. A., Mu, Y. & Kwoh, C. K. Protein-Ligand Blind Docking Using QuickVina-W With Inter-Process Spatio-Temporal Integration. Sci Rep 7, 15451 https://doi.org:10.1038/s41598-017-15571-7 (2017).

97 Waterhouse, A. M., Procter, J. B., Martin, D. M., Clamp, M. & Barton, G. J. Jalview Version 2--a multiple sequence alignment editor and analysis workbench. Bioinformatics 25, 1189–1191 https://doi.org:10.1093/bioinformatics/btp033 (2009).

98 Gierahn, T. M. et al. Seq-Well: portable, low-cost RNA sequencing of single cells at high throughput. Nat Methods 14, 395–398 https://doi.org:10.1038/nmeth.4179 (2017).

99 Hughes, T. K. et al. Second-Strand Synthesis-Based Massively Parallel scRNA-Seq Reveals Cellular States and Molecular Features of Human Inflammatory Skin Pathologies. Immunity 53, 878–894.e877 https://doi.org:10.1016/j.immuni.2020.09.015 (2020).

100 Ibi, D. et al. Antipsychotic-induced Hdac2 transcription via NF-kappaB leads to synaptic and cognitive side effects. Nat Neurosci 20, 1247–1259 https://doi.org:10.1038/nn.4616 (2017).

101 de la Fuente Revenga, M., Vohra, H. Z. & Gonzalez-Maeso, J. Automated quantification of head-twitch response in mice via ear tag reporter coupled with biphasic detection. J Neurosci Methods 334, 108595 https://doi.org:10.1016/j.jneumeth.2020.108595 (2020).

102 Bolger, A. M., Lohse, M. & Usadel, B. Trimmomatic: a flexible trimmer for Illumina sequence data. Bioinformatics 30, 2114–2120 https://doi.org:10.1093/bioinformatics/btu170 (2014).

103 Andrews, S. FastQC: a quality control tool for high throughput sequence data, https://www.bioinformatics.babraham.ac.uk/projects/fastqc/ (2010).

104 Liao, Y., Smyth, G. K. & Shi, W. The Subread aligner: fast, accurate and scalable read mapping by seed-and-vote. Nucleic Acids Res 41, e108 https://doi.org:10.1093/nar/gkt214 (2013).

105 Liao, Y., Smyth, G. K. & Shi, W. featureCounts: an efficient general purpose program for assigning sequence reads to genomic features. Bioinformatics 30, 923–930 https://doi.org:10.1093/bioinformatics/btt656 (2014).

106 Robinson, M. D., McCarthy, D. J. & Smyth, G. K. edgeR: a Bioconductor package for differential expression analysis of digital gene expression data. Bioinformatics 26, 139–140 https://doi.org:10.1093/bioinformatics/btp616 (2010).

107 Benjamini, Y. & Hochberg, Y. Controlling the False Discovery Rate: A Practical and Powerful Approach to Multiple Testing. J R Stat Soc Series B Stat Methodol 57, 289–300 https://doi.org:10.1111/j.2517-6161.1995.tb02031.x (1995).

108 Dai, Q. et al. Striking a balance: regulation of transposable elements by Zfp281 and Mll2 in mouse embryonic stem cells. Nucleic Acids Res 45, 12301–12310 https://doi.org:10.1093/nar/gkx841 (2017).

