## Supplementary Information for "Expanding the HDAC druggable landscape beyond enzymatic activity"

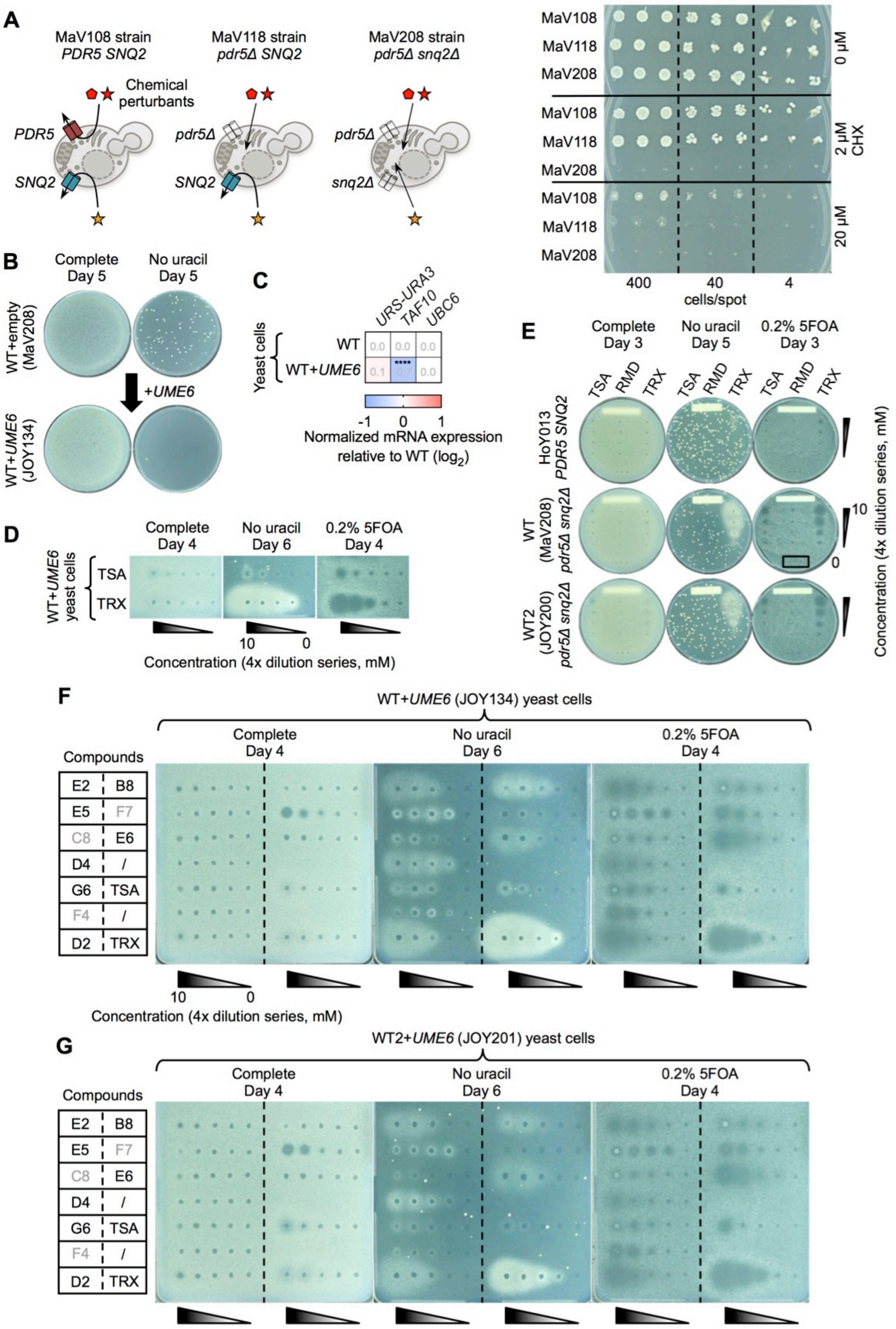


**Figure S1. Drug sensitive yeast strains used to identify Sin3 HDAC complex inhibitors with the *URS-URA3* reporter assay**

(**A**) Deletions of *PDR5* and *SNQ2* efflux pumps in *S. cerevisiae* MaV strains and associated phenotypes on media supplemented with increasing concentrations of cycloheximide (CHX) for different cell densities spotted (n = 3 biologically independent replicates).

(**B**) Reduction of spontaneous growth of $Ura^+$ mutant colonies in the *URS-URA3* reporter assay when the WT MaV208 yeast strain is transformed with a *UME6*-expressing plasmid (WT+*UME6* = JOY134 yeast strain).

(**C**) RT-qPCR analysis of the *URS-URA3* reporter gene in the WT+*UME6* (JOY134) yeast strain relative to the WT (MaV208) strain (n = 3 biologically independent samples per group). Values represent means of replicates. Values for *UBC6* were used to normalize data.

(**D**) Phenotypes of TSA and TRX in the *URS-URA3* reporter assay.

(**E**) Phenotypes of TSA, Romidepsin (RMD) and TRX in the *URS-URA3* reporter assay in the JOY200 yeast strain (WT2) compared to HoY013 and MaV208. The black frame indicates the three control DMSO spots (same position on each plate).

(**F and G**) Phenotypes of TSA, TRX, and the seven Sin3 HDAC complex inhibitors in the *URS-URA3* reporter assay in the WT+*UME6* (JOY134) (F), and WT2+*UME6* (JOY201) (G) yeast strains.

Statistical analysis, (C) two-way ordinary ANOVA with Šidák's post-test correction relative to WT. Raw data are provided in **Source Data**.

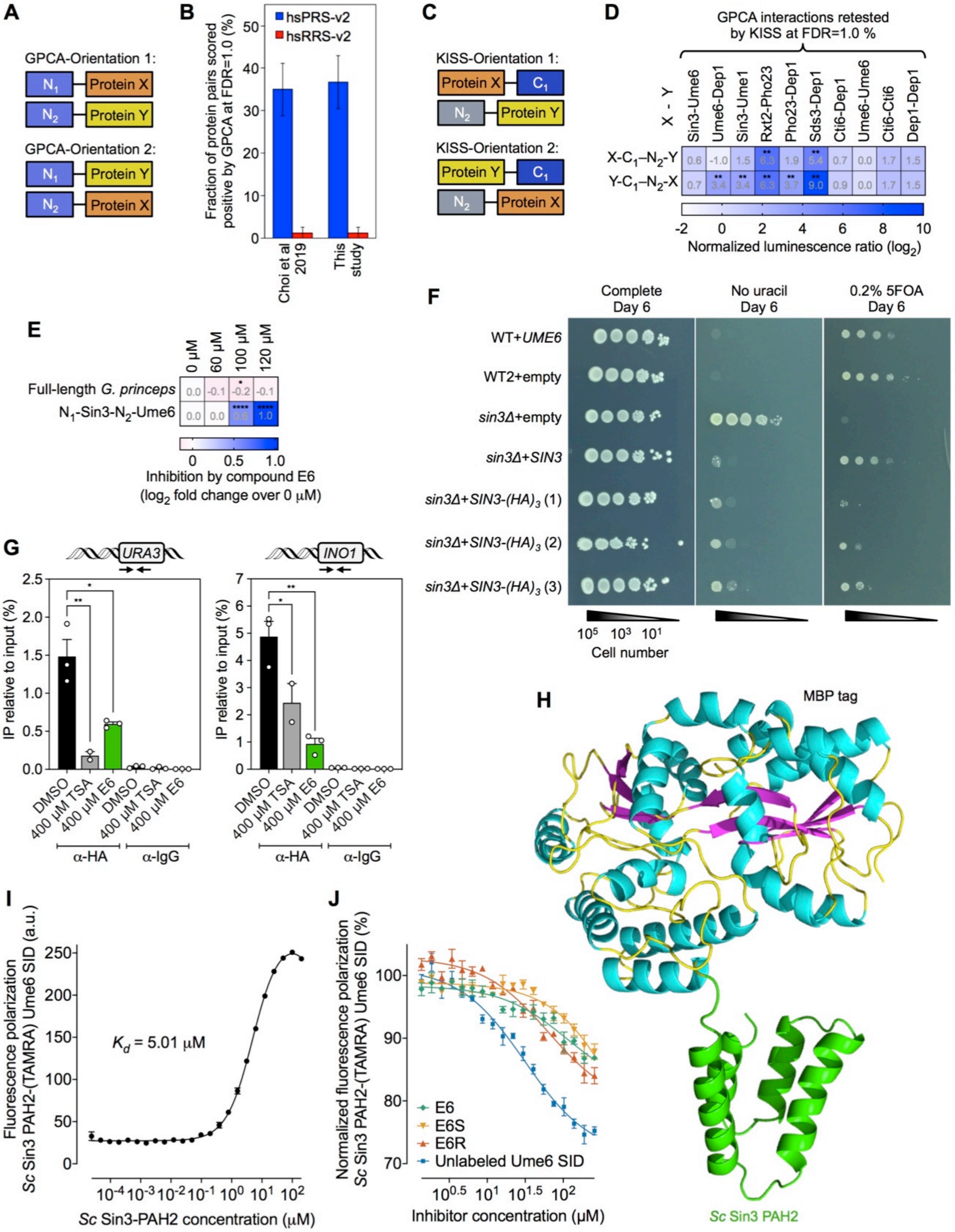


**Figure S2. E6 prevents yeast Sin3/Rpd3L HDAC complex recruitment to Sin3/Rpd3-regulated genes and inhibits the Sin3-Ume6 interaction**

(**A**) GPCA versions used to systematically map intra-complex subunit-subunit interactions.

(**B**) Detection of positive control PPIs (hsPRS-v2) by $N_1N_2$ GPCA at a selected cutoff of 1.0% recovery rate of negative control protein pairs, hsRRS-v2 (FDR = 1.0%) (n = 1 replicate). Assay performances from this study were compared to reported data[39] at the same cutoff. Bars represent recovery rates and error bars standard errors of the proportion.

(**C**) KISS assay versions used to retest intra-complex PPIs identified by $N_1N_2$ GPCA.

(**D**) Retest of intra-complex PPIs from **Figure 2B** by $C_1N_2$ KISS assay versions presented in (C) (n = 6 biologically independent samples per group). Published NLR value for FDR = 1.0%[39] was used as cutoff to score interactions. Values represent means of replicates.

(**E**) Inhibition of the full-length *G. princeps* luciferase and yeast Sin3-Ume6 binary PPI by compound E6 in GPCA (n ≥ 3 biologically independent samples per group).

(**F**) Phenotypes of the Sin3-$(HA)_3$-expressing yeast strain (n = 3 replicates) used for ChIP-qPCR and ChIP-seq analyses, compared to other control strains.

(**G**) ChIP-qPCR analyses for recruitment of the yeast Sin3/Rpd3L HDAC complex at the *URS-URA3* and *INO1* loci following treatment with DMSO control, TSA or E6. Data are compared to α-IgG negative controls (n ≥ 2 biologically independent samples per group). Arrows on the schematics indicate rough positions of the forward and reverse primers. Symbols represent independent repeats, bars means, and error bars SEM.

(**H**) Tertiary structure of the free PAH2 domain of *Sc* Sin3 with the MBP tag.

(**I and J**) FP titration curves showing (I) the interaction of *Sc* Sin3 PAH2 domain (GST-tagged) with Ume6 SID (TAMRA-labeled) (n = 3 biologically independent samples per group), and (J) inhibition of the *Sc* Sin3 PAH2-Ume6 SID (TAMRA-labeled) interaction by unlabeled Ume6 SID, E6, E6S, or E6R (n = 3 biologically independent samples per group) with fitted $IC_{50}$ values of 31.4 μM, 113.0 μM, 597.3 μM and 63.7 μM, respectively. Symbols represent means with SEM, and lines fitted curves.

Statistical analyses, (D) based on empirical p value, (E) two-way ordinary, and (G) one-way ANOVA with Dunnett's multiple comparisons post-test relative to 0 μM (E), or DMSO control in each pull-down (G). Raw data are provided in **Source Data**.

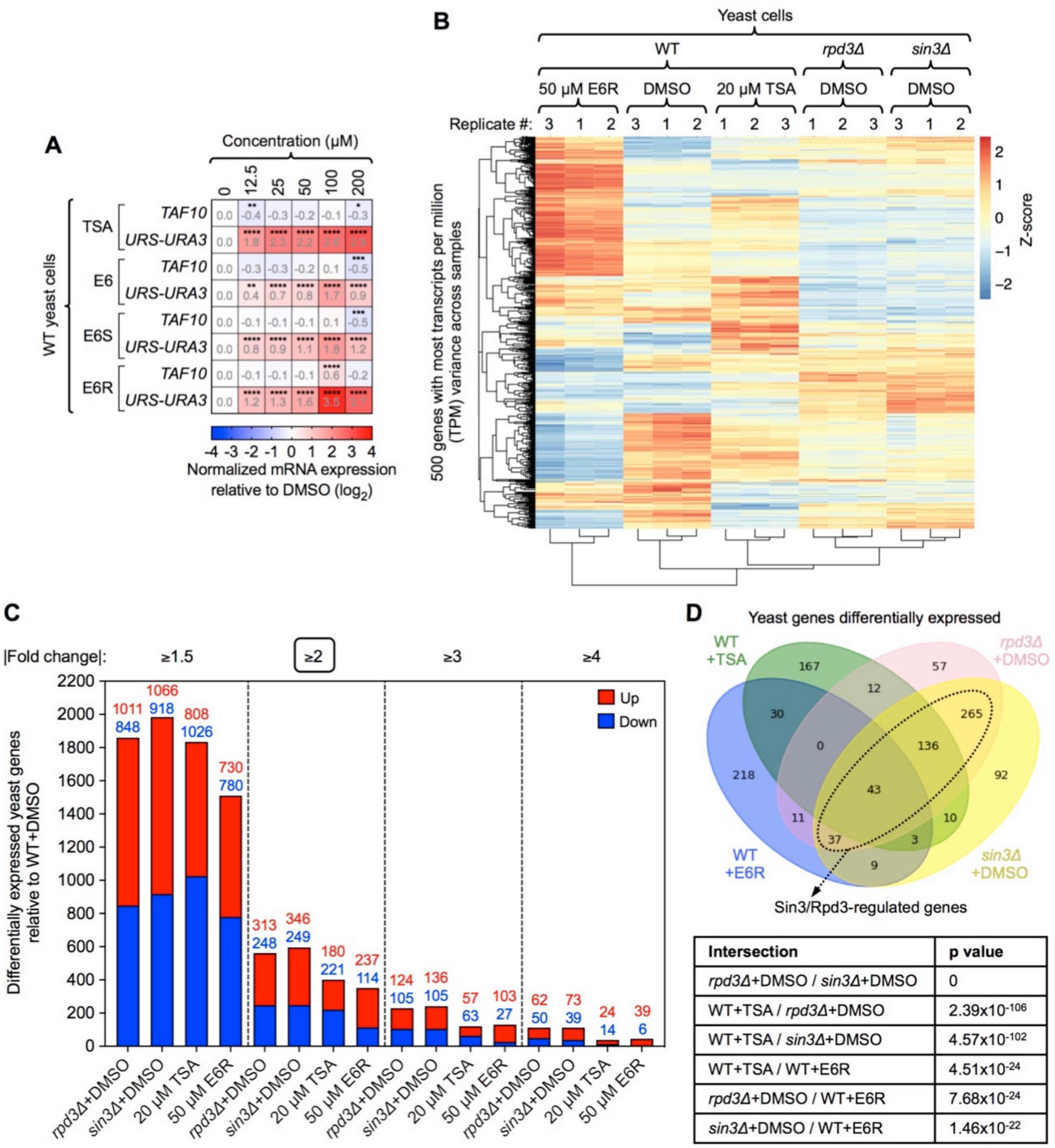


| Intersection | p value |
|---|---|
| *rpd3Δ*+DMSO / *sin3Δ*+DMSO | 0 |
| WT+TSA / *rpd3Δ*+DMSO | $2.39 \times 10^{-106}$ |
| WT+TSA / *sin3Δ*+DMSO | $4.57 \times 10^{-102}$ |
| WT+TSA / WT+E6R | $4.51 \times 10^{-24}$ |
| *rpd3Δ*+DMSO / WT+E6R | $7.68 \times 10^{-24}$ |
| *sin3Δ*+DMSO / WT+E6R | $1.46 \times 10^{-22}$ |

**Figure S3. E6R induces fewer transcriptomic perturbations than TSA among the Sin3/Rpd3-regulated genes in yeast**

(**A**) RT-qPCR analysis of the *URS-URA3* reporter gene and *TAF10* control gene for different doses of TSA, commercial E6, E6S, and E6R (n ≥ 3 biologically independent samples per group). Values represent means of replicates. Values for *UBC6* were used to normalize data.

(**B**) Heat map showing clustering of the different replicates from RNA-seq results in yeast cells (n = 3 biologically independent samples per group).

(**C**) Number of differentially expressed genes in *rpd3Δ*+DMSO and *sin3Δ*+DMSO mutant strains or in WT yeast cells treated with TSA or E6R, relative to WT+DMSO controls. The black frame indicates the fold change cutoff used for analyses.

(**D**) Venn diagram and corresponding p values for the differentially expressed yeast genes identified in (C). The p values of the different overlaps are indicated in the table and the subset of Sin3/Rpd3-regulated genes is circled with a dashed black line.
Statistical analyses, (A) two-way ordinary ANOVA with Dunnett's multiple comparisons post-test relative to DMSO. Raw data are provided in **Source Data**.

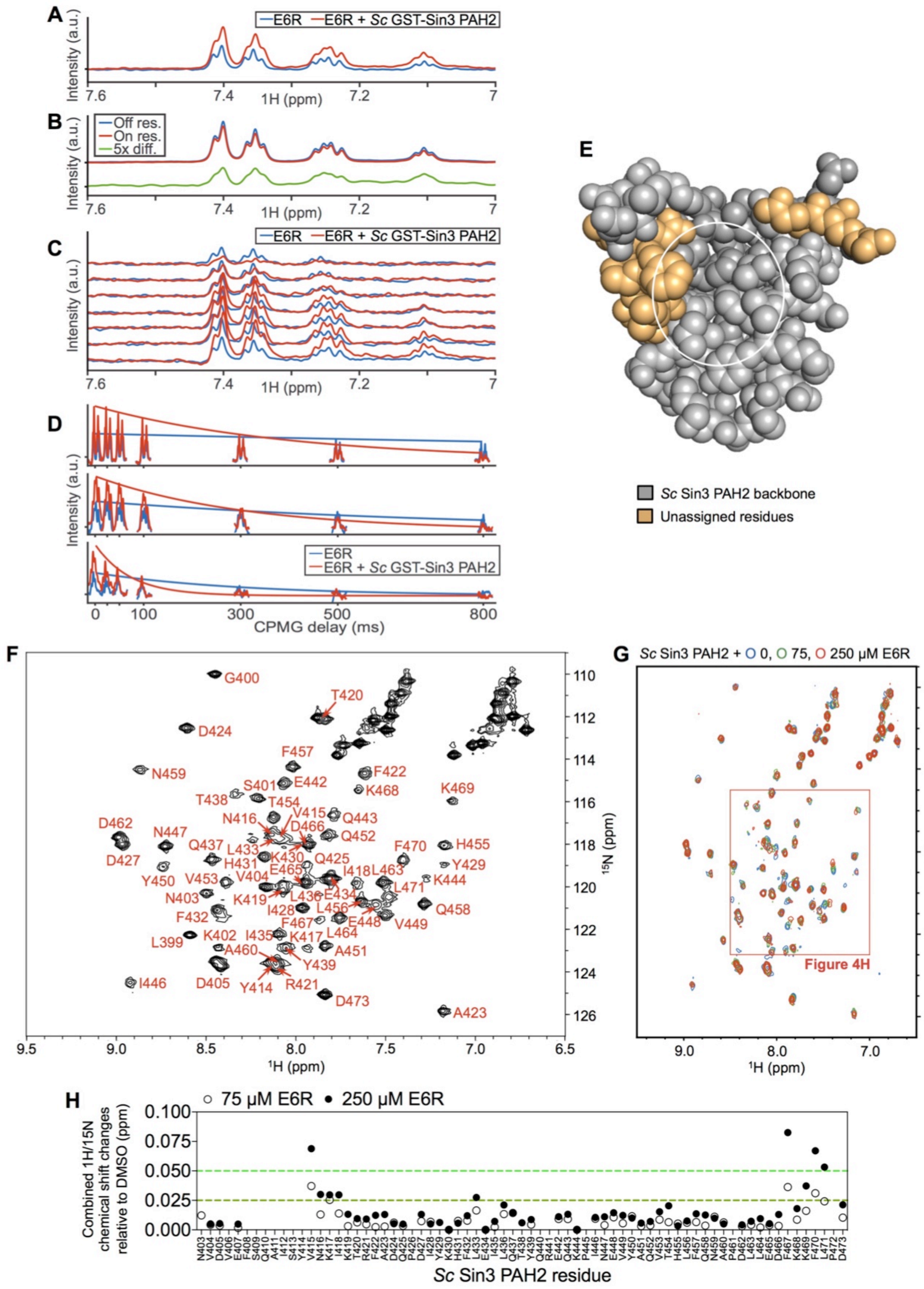


**Figure S4. E6R binds to the yeast Sin3 PAH2 domain**

(**A-D**) Ligand-detected $^{1}$H-NMR spectra for E6R (300 µM) in presence or absence of the *Sc* Sin3 PAH2 domain (10 µM). (A) $^{1}$H-NMR DLB spectra. (B) $^{1}$H-NMR STD spectra where the green curve indicates the difference (x5) between intensities. (C) $^{1}$H-NMR

CPMG-$R_2$ spectra. (D) Regions near 7.4 ppm, 7.25 ppm, and 7.1 ppm from (C) as a function of CPMG delay time.

(**E**) $^{15}N$-HSQC NMR spectrum showing assignment of the free *Sc* Sin3 PAH2 backbone (25 µM). Assigned residues are indicated in red.

(**F**) Space-filled representation of the *Sc* Sin3 PAH2 backbone assignment from (E). The white circle indicates relative position of the deep hydrophobic pocket formed by helices α1 and α2 and where E6R is predicted to bind (**Figure 4J**).

(**G**) $^{15}N$-SOFAST HMQC NMR spectra of untagged *Sc* Sin3 PAH2 (25 µM) with or without E6R. The red frame indicates the area on the graph used in **Figure 4H**.

(**H**) Scatter plot showing the *Sc* Sin3 PAH2 (25 µM) amino acid residues perturbed upon addition of 75 or 250 µM E6R (chemical shifts >0.025 ppm dark green line; >0.05 ppm light green line).

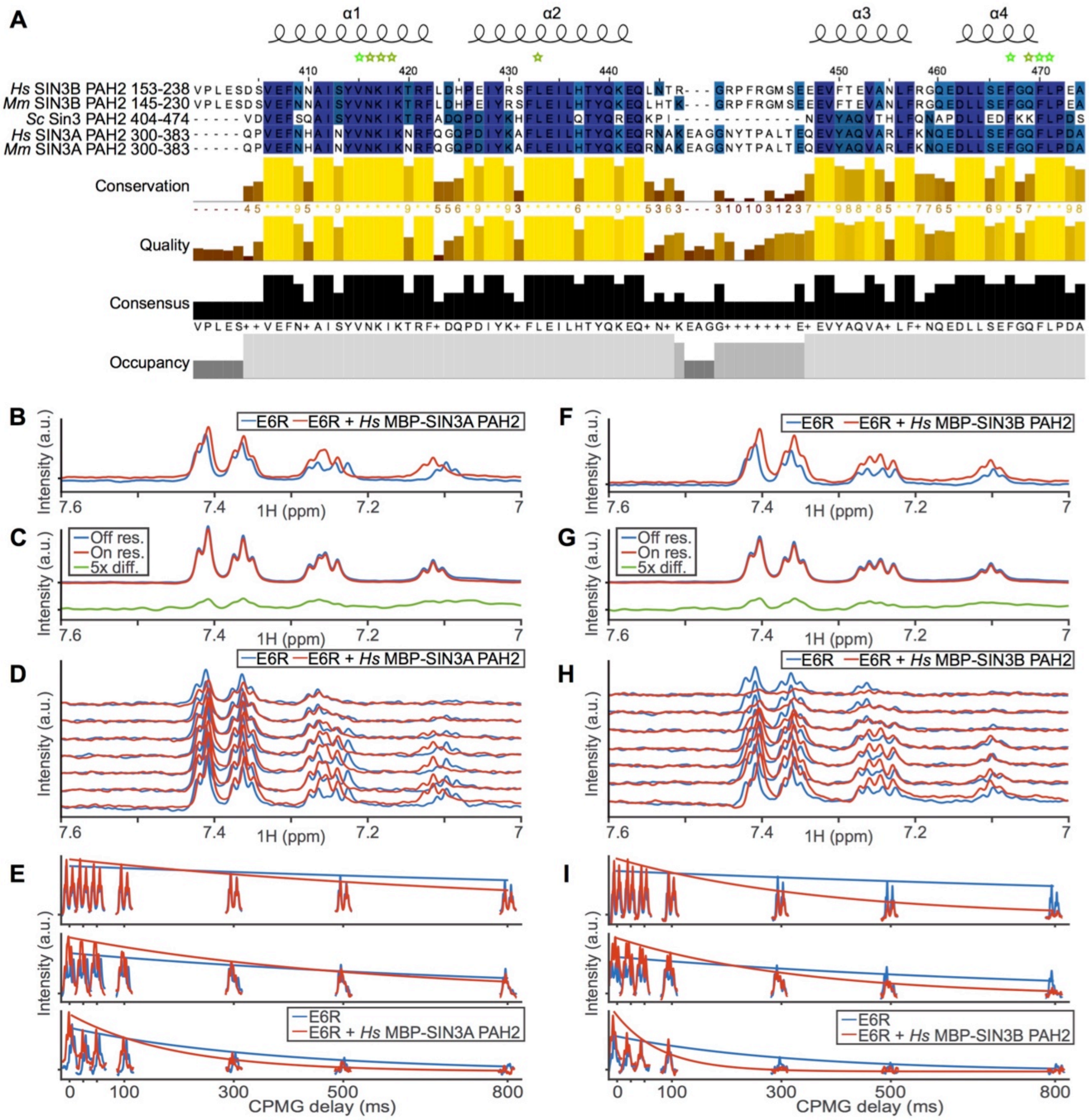


**Figure S5. E6R binds to the conserved human SIN3A and SIN3B PAH2 domains**

(**A**) Profile of amino acid conservation between yeast *Sc*, mouse *M. musculus* (*Mm*) and *H. sapiens* (*Hs*) SIN3 PAH2 domains. Residues mapped to the E6R-binding site in the *Sc* Sin3 PAH2 domain by NMR are highlighted with stars (chemical shifts >0.025 ppm in dark green, and >0.05 ppm in light green). Deep blue corresponds to residues conserved across all SIN3 PAH2 homologs and sky blue to semi-conserved residues. Residues from *Sc* Sin3 PAH2 used as reference for numbering.

(**B-I**) Ligand-detected $^{1}$H-NMR spectra for E6R (300 µM) in presence or absence of the human SIN3A PAH2 (B-E), or SIN3B PAH2 (F-I) domains (10 µM). (B and F) $^{1}$H-NMR DLB spectra. (C and G) $^{1}$H-NMR STD spectra where green curves indicate differences (x5) between intensities. (D and H) $^{1}$H-NMR CPMG-$R_2$ spectra. (E and I) Regions near 7.4 ppm, 7.25 ppm, and 7.1 ppm from (D and H) as a function of CPMG delay times.

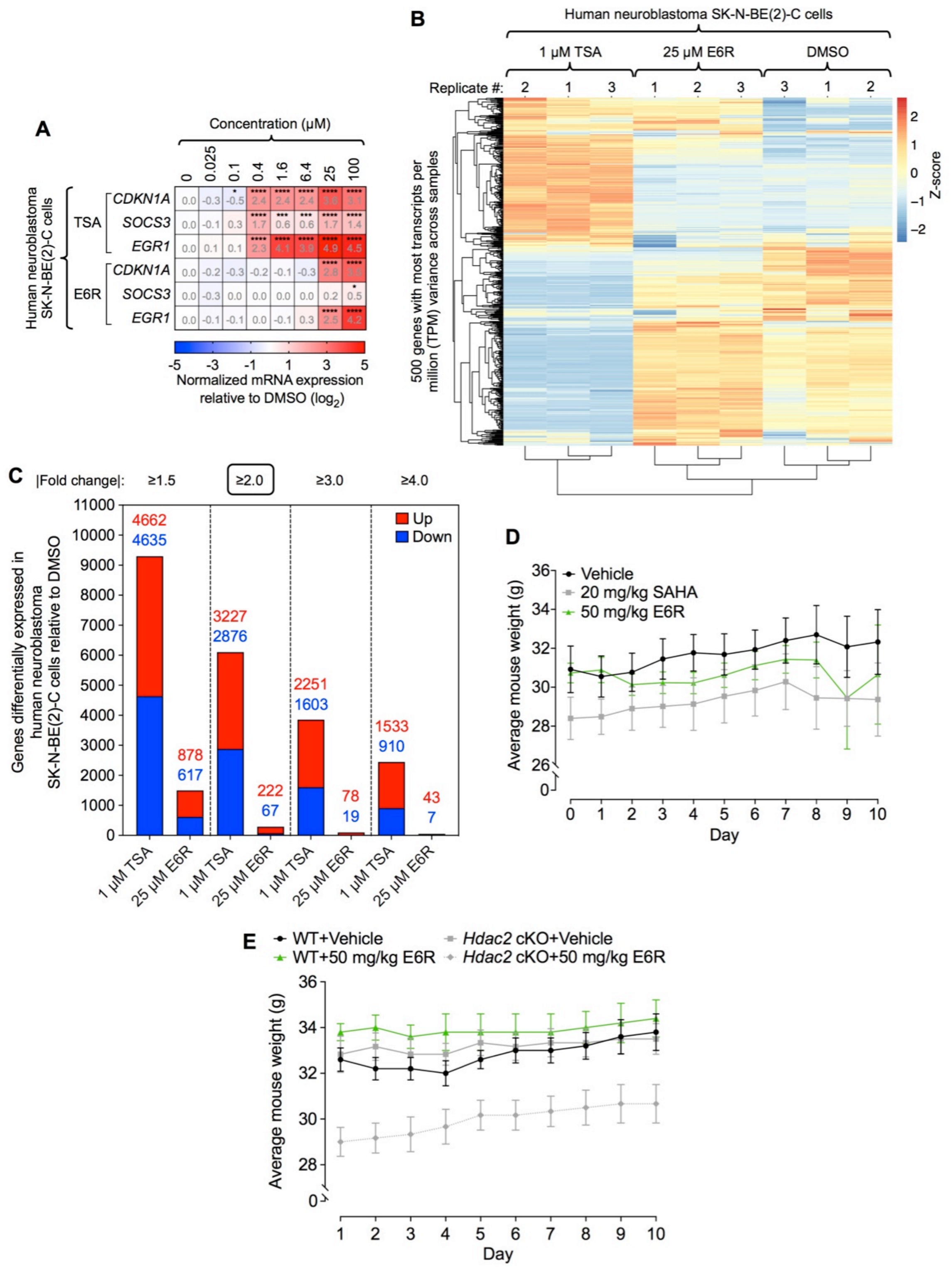


**Figure S6. E6R induces restricted transcriptomic changes compared to enzymatic inhibition and is not toxic *in vivo***

(**A**) RT-qPCR analysis of three human Sin3 HDAC complex-regulated genes in neuroblastoma SK-N-BE(2)-C cells for different doses of TSA and E6R (n ≥ 3 biologically independent samples per group). Values represent means of replicates. Values for *RPS11* were used to normalize data.

(**B**) Heat map showing clustering of the different replicates from RNA-seq results in human neuroblastoma SK-N-BE(2)-C cells (n = 3 biologically independent samples per group).

(**C**) Number of differentially expressed genes in human neuroblastoma SK-N-BE(2)-C cells treated with TSA or E6R, relative to the DMSO control. The black frame indicates the fold change cutoff used for analyses.

(**D and E**) Body weight curves of (D) nude female mice xenografted with human neuroblastoma SK-N-BE(2)-C cells and treated with vehicle, SAHA or E6R (n ≥ 5 biologically independent samples per group), and (E) adult WT and *Hdac2* cKO male mice treated with vehicle or E6R (n ≥ 5 biologically independent samples per group) from which extracted FC tissue samples were used for RNA-seq and ChIP-qPCR analyses in **Figure 6D-H**. Symbols represent means, error bars SEM, and lines connecting curves between data points.

Statistical analyses, (A) two-way ordinary ANOVA with Dunnett's multiple comparisons post-test relative to DMSO. Raw data are provided in **Source Data**.

**Table S1.** Chemical libraries used in this study and results of the chemical genetic screen

<table>
<tr><th rowspan="2">Library name</th><th rowspan="2">Library source</th><th rowspan="2">Library concentration</th><th rowspan="2">Class of molecules</th><th colspan="4">Number of small molecules</th></tr>
<tr><th>Within library</th><th>Tested positive</th><th>Selected for retests</th><th>Tested positive in two strains</th></tr>
<tr><td>BU-CMD 2017</td><td rowspan="7">Harvard Medical School's Institute of Chemistry and Cell Biology – Longwood (ICCB)</td><td>10 mM</td><td>Complex 3D chemotypes</td><td>2,457</td><td>14</td><td>7</td><td>0</td></tr>
<tr><td>ChemDiv6</td><td>10 mM</td><td>Diverse, drug-like properties</td><td>44,000</td><td>250</td><td>49</td><td>7</td></tr>
<tr><td>Prestwick 2</td><td>2 mg/mL</td><td rowspan="3">Known bioactives</td><td>1,120</td><td>1</td><td>1</td><td>0</td></tr>
<tr><td>Biomol 4 – FDA approved</td><td>2 mg/mL</td><td>640</td><td>0</td><td>0</td><td>0</td></tr>
<tr><td>Selleck</td><td>10 mM</td><td>1,858</td><td>8</td><td>2</td><td>0</td></tr>
<tr><td>Human microbiota extracts 1</td><td>15 mg/mL</td><td rowspan="3">Natural products</td><td>309</td><td>0</td><td>0</td><td>0</td></tr>
<tr><td>Human microbiota extracts 2</td><td>10 mg/mL</td><td>160</td><td>0</td><td>0</td><td>0</td></tr>
<tr><td>Natural products set IV</td><td>National Cancer Institute</td><td>10 mM</td><td>419</td><td>4</td><td>1</td><td>0</td></tr>
<tr><td>2P2I3D-v2</td><td>Aix-Marseille University's Integrative Structural and Chemical Biology group</td><td>1 mM</td><td>PPI-oriented</td><td>1,278</td><td>0</td><td>0</td><td>0</td></tr>
<tr><td colspan="4">Total</td><td>52,234</td><td>277 (275 unique)</td><td>60 (59 unique)</td><td>7</td></tr>
</table>

**Table S2.** List of proteins generated by mass spectrometry analyses of purified human FLAG-HDAC1 complexes and comparison to known interactors of HDAC1/2 and SIN3A/B subunits from the literature. Yeast *S. cerevisiae* orthologs were obtained from PANTHER. Proteins identified in at least two out of the three biologically independent replicates were used to generate data

| Gene name | UniProt code | Known interactor(s) | HDAC complex(es) | Yeast orthologs(s) |
|---|---|---|---|---|
| *HSPD1* | P10809 | HDAC1 | / | Hsp60, Tcm62 |
| *RERE* | Q9P2R6 | HDAC1/2 | ELM-SANT | / |
| *ARID4A* | P29374 | HDAC1, SIN3A | Sin3 | Swi1, Rsc9 |
| *ARID4B* | Q4LE39 | HDAC1/2 | | / |
| *BRMS1* | H0YCF7 | HDAC1/2, SIN3A | | Sds3, Dep1 |
| *BRMS1L* | H0YHD0 | HDAC1 | | |
| *CHD3* | Q12873 | HDAC1/2 | NuRD | / |
| *CHD4* | A0A2R8Y5J0 | | | |
| *CDK2AP1* | O14519 | | | |
| *DNTTIP1* | Q9H147 | | / | |
| *GSE1* | Q14687 | | CoREST | |
| *HMG20A* | H0YKL0 | | | |
| *HDAC1* | Q13547 | HDAC1/2, SIN3A/B | Sin3, CoREST, NuRD, MiDAC, SHIP | Rpd3 |
| *HDAC2* | Q92769 | | | |
| *SAP130* | Q9H0E3 | HDAC1/2 | Sin3 | / |
| *H4C1* | P62805 | | / | Hhf2 |
| *RBBP4* | Q09028 | HDAC1/2, SIN3A | Sin3 | Hat2, Msi1, Wtm1, Wtm2, Ume1 |
| *RBBP7* | Q16576 | | | |
| *ING1* | A0A087WXF7 | | | Pho23, Yng1, Yng2 |
| *KPNA4* | O00629 | HDAC1 | / | Srp1 |
| *ING2* | C9J4X5 | HDAC1, SIN3A | Sin3 | / |
| *KDM1A* | R4GMQ1 | HDAC1/2 | CoREST | / |
| *MIER1* | Q8N108 | | ELM-SANT | |
| *MIER2* | Q8N344 | | | |
| *MTA1* | E7ESY4 | | NuRD | |
| *MTA2* | O94776 | HDAC1/2, SIN3A | | |
| *MTA3* | E7EQY4 | HDAC2 | | |
| *MBD2* | Q9UBB5 | HDAC1/2, SIN3A | | |
| *MBD3* | O95983 | HDAC1/2 | | |
| *SIN3A* | Q96ST3 | | Sin3, CoREST | Sin3 |
| *SIN3B* | O75182 | | | |
| *PHF21A* | Q96BD5 | | CoREST | / |
| *PFDN5* | Q99471 | HDAC1 | / | Gim5 |
| *DDX17* | A0A0U1RQJ0 | HDAC1/2 | | Dbp2 |
| *RCOR1* | Q9UKL0 | | CoREST | / |
| *RCOR2* | Q8IZ40 | HDAC2 | | |
| *RCOR3* | Q9P2K3 | HDAC1/2 | | |
| *SUDS3* | Q9H7L9 | HDAC1/2, SIN3A/B | Sin3 | Sds3, Dep1 |
| *SINHCAF* | Q9NP50 | HDAC1 | | / |
| *HMG20B* | Q9P0W2 | HDAC1/2 | CoREST | |
| *TCP1* | P17987 | HDAC1 | / | Tcp1 |
| *CCT8* | P50990 | | | Cct8 |
| *GATAD2A* | Q86YP4 | HDAC1/2 | | / |
| *GATAD2B* | Q8WXI9 | | | |
| *TRERF1* | Q96PN7 | HDAC1 | MiDAC | |
| *OGT* | C9JZL3 | SIN3A | / | |
| *C16orf87* | Q6PH81 | HDAC1 | | |
| *ZMYM2* | Q9UBW7 | HDAC1/2 | CoREST | |
| *ZMYM3* | Q14202 | HDAC1 | | |
| *ZNF217* | O75362 | HDAC1/2 | | |

**Table S3.** Crystallographic data for the yeast *S. cerevisiae* Sin3 PAH2 domain alone or in complex with the Ume6 SID peptide

| Structure name | SIN3.0402-0473.DGL8HY | SIN3.UME6.CF1098 |
|---|---|---|
| Ligand | / | Ume6 peptide |
| RCSB accession code | 6XDJ | 6XAW |
| Data collection [a] | | |
| Space group | P 1 $2_1$ 1 | P $4_3$ 2 2 |
| Cell dimensions<br>a, b, c (Å)<br>a, b, g (°) | <br>79.73, 144.86, 97.09<br>90, 107.97, 90 | <br>56.77, 56.77, 63.90<br>90, 90, 90 |
| Resolution (Å) [b] | 75.84 – 2.2 (2.279 – 2.2) | 33.99 – 1.84 (1.906 – 1.84) |
| $R_{pim}$ | 0.06203 (0.695) | 0.04904 (0.6838) |
| I / sI | 7.85 (1.07) | 16.40 (1.10) |
| Completeness (%) | 99.09 (98.08) | 99.54 (99.13) |
| Redundancy | 3.5 (3.3) | 12.3 (12.6) |
| Refinement | | |
| No. reflections, unique | 105108 (10363) | 9504 (907) |
| $R_{work}$ / $R_{free}$ | 0.2057 (0.3189) / 0.2399 (0.3566) | 0.1920 (0.4023) / 0.2122 (0.4140) |
| No. non-hydrogen atoms<br>Protein<br>Ligand/ion<br>Water | <br>12508<br>125<br>702 | <br>701<br>4<br>93 |
| B-factors<br>Protein<br>Ligand/ion<br>Water | <br>51.73<br>43.38<br>52.26 | <br>29.60<br>41.96<br>42.86 |
| R.m.s. deviations<br>Bond lengths (Å)<br>Bond angles (°) | <br>0.006<br>1.02 | <br>0.007<br>1.05 |
| Ramachandran plot | | |
| Favored (%) | 98.15 | 100.00 |
| Allowed (%) | 1.85 | 0.00 |
| Not allowed (%) | 0.00 | 0.00 |

[a] A single crystal was used to collect data for each of the structures reported here
[b] Values in parentheses are for the highest resolution shell

**Table S4.** Human genes differentially expressed by TSA and/or E6R (relative to DMSO) in the bulk RNA-seq and scRNA-seq studies in neuroblastoma SK-N-BE(2)-C cells

| Differentially expressed genes by both TSA and E6R in the bulk RNA-seq and scRNA-seq studies | Up- or downregulated by TSA and/or E6R |
|---|---|
| *ADM2* | Upregulated by TSA |
| *ARC* | |
| *CDKN1A* | |
| *DOCK6* | |
| *DUSP5* | |
| *EGR1* | |
| *ERN1* | |
| *KLF2* | |
| *MAPKAPK2* | |
| *MTRNR2L4* | |
| *NCKAP5L* | |
| *NECTIN2* | |
| *PGPEP1* | |
| *PKD1* | |
| *PLCD3* | |
| *PRLHR* | |
| *RHBDD2* | |
| *RPS6KA2* | |
| *SCN4A* | |
| *SESN2* | |
| *YPEL5* | |
| *ZNF487* | |
| *ANKMY2* | Upregulated by TSA and E6R |
| *ASL* | |
| *BBC3* | |
| *CALCB* | |
| *CARS* | |
| *CASZ1* | |
| *CPEB4* | |
| *CTH* | |
| *DGKI* | |
| *DUSP10* | |
| *FOS* | |
| *IER2* | |
| *IER3* | |
| *LAT2* | |
| *MSX2* | |
| *NR4A1* | |
| *PPP1R15A* | |
| *SYT5* | |
| *TOX2* | |
| *ULBP1* | |
| *VGF* | |
| *NR4A3* | Upregulated by E6R |
| *ABCA12* | Downregulated by TSA |
| *AQP1* | |
| *ATXN1* | |
| *DCLK3* | |
| *MCHR1* | |
| *NXPH2* | |
| *RHOU* | |
| *RPH3A* | |
| *SLC12A3* | |
| *SLITRK6* | |
| *APLN* | Downregulated by TSA and E6R |
| *SH3TC2* | |
| *TRIM29* | |
| *CABP7* | Downregulated by E6R |
| *SLITRK1* | |

**Table S5.** Yeast strains used in this study

| Yeast strain | Source | Alias in this study |
|---|---|---|
| *S. cerevisiae*: Strain background: MaV108: *MATa leu2-3,112 trp1-901 his3-200 ura3-52 ade2-101 gal4Δ gal80Δ SPAL10::URA3@ura3 GAL1::LacZ can1$^R$ cyh2$^R$* | Vidal et al.[31] | N/A |
| *S. cerevisiae*: Strain background: MaV118: *MaV108*, *pdr5::HIS3* | This paper | N/A |
| *S. cerevisiae*: Strain background: MaV208 (defined as WT in this paper): *MaV118*, *snq2::KanMX* | | WT |
| *S. cerevisiae*: Strain background: JOY111: *MaV208*, *rpd3Δ::NatMX* | | *rpd3Δ* |
| *S. cerevisiae*: Strain background: JOY112: *MaV208*, *ume6Δ::NatMX* | | *ume6Δ* |
| *S. cerevisiae*: Strain background: JOY113: *MaV208*, *sds3Δ::NatMX* | | *sds3Δ* |
| *S. cerevisiae*: Strain background: JOY114: *MaV208*, *sap30Δ::NatMX* | | *sap30Δ* |
| *S. cerevisiae*: Strain background: JOY115: *MaV208*, *pho23Δ::NatMX* | | *pho23Δ* |
| *S. cerevisiae*: Strain background: JOY116: *MaV208*, *sin3Δ::NatMX* | | *sin3Δ* |
| *S. cerevisiae*: Strain background: JOY117: *MaV208*, *rxt2Δ::NatMX* | | *rxt2Δ* |
| *S. cerevisiae*: Strain background: JOY118: *MaV208*, *rxt3Δ::NatMX* | | *rxt3Δ* |
| *S. cerevisiae*: Strain background: JOY119: *MaV208*, *dep1Δ::NatMX* | | *dep1Δ* |
| *S. cerevisiae*: Strain background: JOY120: *MaV208*, *cti6Δ::NatMX* | | *cti6Δ* |
| *S. cerevisiae*: Strain background: JOY121: *MaV208*, *ume1Δ::NatMX* | | *ume1Δ* |
| *S. cerevisiae*: Strain background: JOY122: *MaV208*, *ash1Δ::NatMX* | | *ash1Δ* |
| *S. cerevisiae*: Strain background: JOY128: *MaV208*, *(spal10::ura3)Δ::HpHMX@ura3* | | *ura3Δ* |
| *S. cerevisiae*: Strain background: JOY134: *MaV208*, *CEN LEU2 UME6* (pAR128 plasmid) | | WT+*UME6* |
| *S. cerevisiae*: Strain background: HoY013: *Y8800*, *SPAL10::URA3@ura3* | | N/A |
| *S. cerevisiae*: Strain background: JOY200: *HoY013*, *snq2Δ::KanMX pdr5Δ::NatMX* | | WT2 |
| *S. cerevisiae*: Strain background: JOY201: *JOY200*, *CEN LEU2 UME6* (pAR128 plasmid) | | WT2+*UME6* |
| *S. cerevisiae*: Strain background: JOY137: *JOY112*, *sin3Δ::TRP1* | | *sin3Δ/ume6Δ* |

**Table S6.** Experimental conditions used for the different yeast cell selections in this study

| Figure(s) | Yeast strain(s) | Selection media | Growth conditions and cell densities |
|---|---|---|---|
| 1B | Indicated in Figure 1B | Complete = SC<br>No uracil = SC-URA<br>0.2% 5FOA = SC + 0.2% 5FOA | Indicated in Figure 1B |
| 1C | Indicated in Figure 1C | YPD | Growth in YPD up to $OD_{600\ nm}$ = 0.5 then incubate for 2 h |
| 1E | WT+*UME6* = MaV208+pAR128 = JOY134 | Complete = SC-LEU<br>No uracil = SC-LEU-URA<br>0.2% 5FOA = SC-LEU+0.2% 5FOA | Complete = $2x10^7$ cells/plate<br>No uracil = $5x10^9$ cells/plate<br>0.2% 5FOA = $2x10^7$ cells/plate |
| 1F | WT = MaV208 | YPD | Growth in YPD up to $OD_{600\ nm}$ = 0.5 then incubate for 2 h with compound |
| 3A-F | *sin3Δ+SIN3-(HA)$_3$* = JOY116+YEplac181-Sin3-$(HA)_3$ | YPD | Growth in YPD up to $OD_{600\ nm}$ = 0.7 then incubate for 2 h with compound |
| 3G | Indicated in Figure 3G | YPD | Growth in YPD up to $OD_{600\ nm}$ = 0.5 then incubate for 2 h with compound |
| 4F | WT+*UME6* = MaV208+pAR128 = JOY134 | Complete = SC-LEU<br>No uracil = SC-LEU-URA<br>0.2% 5FOA = SC-LEU+0.2% 5FOA | Complete = $2x10^7$ cells/plate<br>No uracil = $5x10^9$ cells/plate<br>0.2% 5FOA = $2x10^7$ cells/plate |
| 4G, S3 | WT = MaV208<br>*rpd3Δ* = JOY111<br>*sin3Δ* = JOY116 | YPD | Growth in YPD up to $OD_{600\ nm}$ = 0.5 then incubate for 2 h with DMSO or compound |
| S1A | Indicated in Figure S1A | CHX = SC + CHX | Indicated in Figure S1A |
| S1B | WT+empty = MaV208+pRS415<br>WT+*UME6* = MaV208+pAR128 = JOY134 | Complete = SC-LEU<br>No uracil = SC-LEU-URA | Complete = $4x10^8$ cells/plate<br>No uracil = $4x10^8$ cells/plate |
| S1C | WT = MaV208<br>WT+*UME6* = MaV208+pAR128 = JOY134 | YPD | Growth in YPD up to $OD_{600\ nm}$ = 0.5 then incubate for 2 h |
| S1D | WT+*UME6* = MaV208+pAR128 = JOY134 | Complete = SC-LEU<br>No uracil = SC-LEU-URA<br>0.2% 5FOA = SC-LEU+0.2% 5FOA | Complete = $2x10^7$ cells/plate<br>No uracil = $5x10^9$ cells/plate<br>0.2% 5FOA = $2x10^7$ cells/plate |
| S1E | Indicated in Figure S1E | Complete = SC<br>No uracil = SC-URA<br>0.2% 5FOA = SC+0.2% 5FOA | Complete = $1x10^8$ cells/plate<br>No uracil = $1x10^8$ cells/plate<br>0.2% 5FOA = $1x10^8$ cells/plate |
| S1F | WT+*UME6* = MaV208+pAR128 = JOY134 | Complete = SC-LEU<br>No uracil = SC-LEU-URA<br>0.2% 5FOA = SC-LEU+0.2% 5FOA | Complete = $2x10^7$ cells/plate<br>No uracil = $5x10^9$ cells/plate<br>0.2% 5FOA = $2x10^7$ cells/plate |
| S1G | WT2+*UME6* = JOY200+pAR128 = JOY201 | Complete = SC-LEU<br>No uracil = SC-LEU-URA<br>0.2% 5FOA = SC-LEU+0.2% 5FOA | Complete = $2x10^7$ cells/plate<br>No uracil = $5x10^9$ cells/plate<br>0.2% 5FOA = $2x10^7$ cells/plate |
| S2F | WT+*UME6* = MaV208+pAR128 = JOY134<br>WT2+empty = JOY200+pDEST-DB<br>*sin3Δ*+empty = JOY116+pDEST-DB<br>*sin3Δ+SIN3* = JOY116+pAR124<br>*sin3Δ+SIN3-(HA)$_3$* = JOY116+YEplac181-Sin3-$(HA)_3$ | Complete = SC-LEU<br>No uracil = SC-LEU-URA<br>0.2% 5FOA = SC-LEU+0.2% 5FOA | Indicated in Figure S2F |
| S2G | *sin3Δ+SIN3-(HA)$_3$* = JOY116+YEplac181-Sin3-$(HA)_3$ | YPD | Growth in YPD up to $OD_{600\ nm}$ = 0.7 then incubate for 2 h with compound |

**Table S7.** Primers used in this study

| Oligonucleotides | Source |
|---|---|
| Primers for RT-qPCR and ChIP-qPCR: *URA3*:<br>Forward: TGACATTGCGAAGAGCGACA<br>Reverse: GAGACCACATCATCCACGGT | This paper |
| Primers for RT-qPCR and ChIP-qPCR: *INO1*:<br>Forward: TATATGAAGCCCGTCGGGGA<br>Reverse: TCGATGATCAAGGGCGTAGC | |
| Primers for RT-qPCR: *SPO13*:<br>Forward: AACGGCCGGAGTTGCTTTAT<br>Reverse: CTGCCCTTGGTGTCTCTTGG | |
| Primers for RT-qPCR: *TRK2*:<br>Forward: GCGCTTATGGTACAGTGGGT<br>Reverse: CCAGTTTGTCACTTGGCAGC | |
| Primers for RT-qPCR: *IME2*:<br>Forward: GGTGATGCCTCTTTAGGCGA<br>Reverse: TGAGGCTCGAACTTTTCCCG | |
| Primers for RT-qPCR: *CAR1*:<br>Forward: ATACGGCATCAACGCTGTCA<br>Reverse: GGTCAACCCACCTCTCACTG | |
| Primers for RT-qPCR: *UBC6*:<br>Forward: AATTGGATGAGGGGGATGCG<br>Reverse: CGCTTGTTCAGCGCGTATTC | |
| Primers for RT-qPCR: *TAF10*:<br>Forward: AACAACAGTCAGGCGAGAGC<br>Reverse: AACAGCGCTACTGAGATCGT | |
| Primers for RT-qPCR: *RPS11*:<br>Forward: ACATCCGCAAGTACAACCGCTT<br>Reverse: TGGTGACCTTGAGCACGTTGAAG | |
| Primers for RT-qPCR: *SNRPD3*:<br>Forward: GGCCTGCTGGGATTATAGGTGT<br>Reverse: AGGGAGATGGGTGAGAGGAAGTAAAT | |
| Primers for RT-qPCR: *CDKN1A*:<br>Forward: GGAGTATTGGGGTCTGACCCCAAA<br>Reverse: TTGAGCACCTGCTGTATATTCAGCATT | |
| Primers for RT-qPCR: *SOCS3*:<br>Forward: GTTTACAATCTGCCTCAATCACTCTGTCTT<br>Reverse: GGATTTTGTTGAGTTCTTCAAGCATCTCCTAA | |
| Primers for RT-qPCR: *EGR1*:<br>Forward: TTGCTGATGGCTTGACATGTGCAATT<br>Reverse: CGAAGCTCAGCTCAGCCCTCTT | |
| Primers for ChIP-qPCR: *Homer1*:<br>Forward: CTGTCAAAGTTGCACCAGCA<br>Reverse: GACGCTGAAGCTTCTGGAGG | |

**Note S1.** Agar diffusion assay to select for small molecules

The design of the assay is such that WT *URS-URA3* yeast cells are plated onto a solid medium lacking uracil. Compounds are subsequently spotted onto this lawn of non-growing $Ura^-$ cells, and allowed to diffuse to produce a gradient of concentrations. After penetrating cells, compounds that can inhibit the Sin3/Rpd3L HDAC complex confer a $Ura^+$ phenotype which appears as a ring of growth around the original spot, following incubation for 2-7 days. This primary positive selection for growth has obvious advantages over settings using negative readouts, such as reduction of reporter signal or cell viability. It filters out compounds that are cytotoxic and non cell-permeable, while preventing the constraint of having to define a particular concentration for the entire screening because of the gradient that forms around the spotted compounds.

**Note S2.** Assessment of HDAC enzymatic activity

The single reagent addition, luminescent system developed by Promega allows one to measure deacetylase activity of HDAC class I/II enzymes using live cells, cellular extracts, or purified proteins. The assay uses a cell permeable, acetylated, luminogenic peptide substrate that can be deacetylated in the presence of functional HDAC enzymes to produce luminescence upon addition of luciferase. In summary, the experimental setting is such that normal HDAC activity is represented by relatively high luminescence values, while enzymatic inhibitors decrease the luminescence signals.
